# Beyond the Highlands: Climate Drives Evolutionary Connections Between Ancient Neotropical Mountains and Lowland Biomes

**DOI:** 10.64898/2026.02.10.705068

**Authors:** Yago Barros-Souza, Monique Maianne, Rafael F. Barduzzi, Leonardo M. Borges

## Abstract

**Aim:** The assembly of montane plant communities through time is underlain by historical and abiotic factors. However, the extent of evolutionary connectivity between ancient highland ecosystems and surrounding lowlands remains unclear. Here, we investigate the evolutionary connections between the *campos rupestres*, a hyperdiverse and fragmented montane vegetation complex in eastern South America, and lowland biomes surrounding it: savannas, rainforests, and seasonally dry tropical forests.

**Location:** Eastern South America.

**Time period:** Cenozoic.

**Major taxa studied:** Flowering plants.

**Methods:** Using phylogenetic beta diversity analyses for 13 angiosperm clades, we assess the degree of lineage dissimilarity between *campos rupestres* subregions and adjacent biomes. We also apply generalized dissimilarity modeling to determine the role of climate, soil, and geographic distance in shaping spatial patterns of phylogenetic composition.

**Results:** Our results reveal high lineage permeability between *campos rupestres* and surrounding biomes, with lineage sharing largely reflecting biome adjacency. This pattern is mainly driven by shared climatic conditions, which are the strongest predictors of phylogenetic dissimilarity.

**Main conclusions:** We highlight the importance of lineage exchange between lowland and montane environments for the assembly of highland floras. By showing that lineage movements across biome boundaries have been common over time and spatial scales, our study challenges the idea that ancient Neotropical mountains are isolated sky-islands. Instead, we emphasize the dynamic nature of montane plant diversity and the pivotal role of climate in shaping evolutionary connections between highlands and lowlands.

## 1 Introduction

The spatial distribution of plant lineages is shaped by historical contingencies and environmental gradients (Cai et al., 2024; Moulatlet et al., 2023; Ringelberg et al., 2023). Yet lineage distribution changes over time via dispersal and niche evolution (Chase, 2011). The likelihood that a lineage will overcome biome limits, for example, is strongly influenced by biome size and age, preexisting traits, geographic adjacency, and environmental similarity (Donoghue & Edwards, 2014). Indeed, transitions between vegetation types and long-distance dispersal to similar biomes are both common across plant lineages (Crisp et al., 2009; Dale et al., 2020; Zizka et al., 2020) and often play a key role in community assembly. For instance, in the Cape Floristic Region, transitions from forest to fire-prone heathlands were linked to functional trait evolution and diversification bursts (Onstein et al., 2014), while in the Central Andes, mountain uplift created novel environments where immigration and ecological sorting of pre-adapted clades largely structured community assembly (Linan et al., 2021). Nonetheless, given the often steep environmental gradients occurring within short distances, the evolutionary relationship between montane and lowland environments is one of the most intriguing yet least understood across plant evolution.

This is the case for the *campos rupestres* (CR), a montane vegetation complex in eastern South America surrounded by savannas, rainforests, and seasonally dry tropical forests (SDTFs; Figure 1). The CR are highly fragmented habitats that harbor an astonishing diversity of ca. 5000 plant species in an area smaller than Ireland (Silveira et al., 2016). Although embodying a rich vegetation mosaic, they are dominated by rock-dwelling plant communities and grasslands established on infertile soils and ancient geological formations (with ca. 640 My; Alkmim, 2012). Ranging between 900 to over 2000 m elevation (for details, see Fernandes, 2016), the mountains where CR abound are primarily the result of Paleoproterozoic sedimentary deposits uplifted during the Mesoproterozoic and Neoproterozoic, with additional contributions from later, lower-magnitude regional uplift (Saadi, 1995). From the Paleozoic to the Cenozoic, prolonged erosion and weathering shaped the current relief (Schaefer, 2016), favoring a landscape with altitudinal gradients not as steep and sharply defined as those in recently formed mountain chains, such as the Andes (Antonelli et al., 2018). The CR occur mainly in the Espinhaço Range, a mountain chain subdivided into Southern (SE) and Northern (NE) subregions (Silveira et al., 2016), located in eastern Brazil and with a plant species/area ratio 40 times that of the Amazon forest (Rapini et al., 2021; Cardoso et al., 2017). Other CR areas include the *Serra da Canastra* (SC; Canastra Range), the Brazilian Central Plateau (BCP; Figure 1), and a few small, disjunct patches scattered across South America (Silveira et al., 2016).

**Figure 1.**
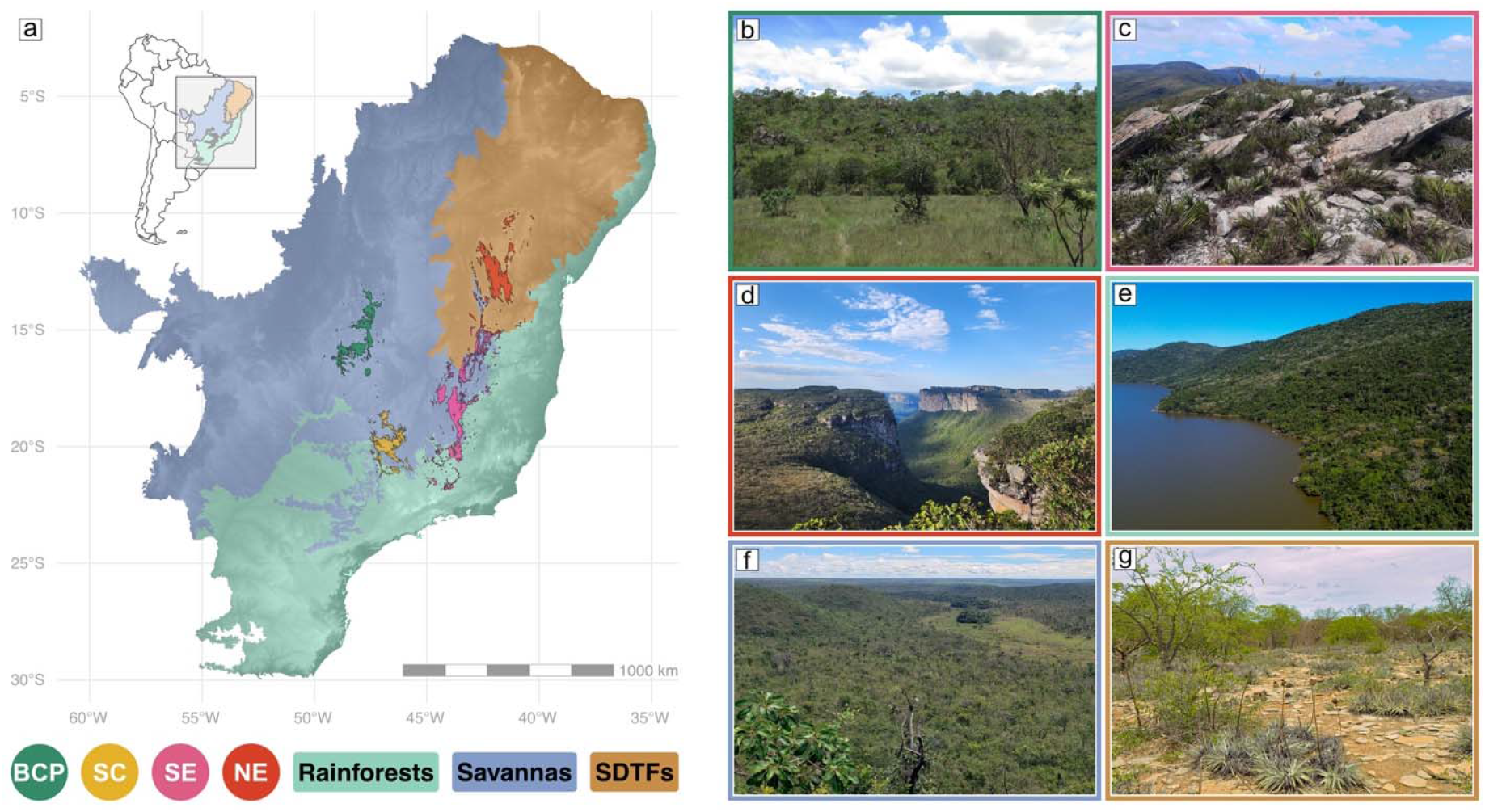
Documented distribution of *campos rupestres* subregions and surrounding biomes in eastern South America, and photographs depicting their typical landscapes. **a)** Subregions are shown in different colors and by filled circles. Surrounding biomes are likewise shown in distinct colors and by rounded rectangles. **b)** *Campos rupestres* at Parque Nacional da Chapada dos Veadeiros (BCP); **c)** *Campos rupestres* at Parque Nacional da Serra do Cipó (SE); **d)** *Campos rupestres* at Parque Nacional da Chapada Diamantina (NE); **e)** Atlantic rainforest at Lagoa do Peri; **f)** Savannas at Parque Nacional das Emas; **g)** Seasonally dry tropical forest at Gruta dos Brejões. Photos by Monique Maianne (**b**), Rafael Barduzzi (**c**), Marcelo Trovó (**d**), Elsimar Silveira da Silva (**e**), Leonardo Borges (**f**), and Domingos Cardoso (**g**). Abbreviations: BCP = Brazilian Central Plateau; SC = *Serra da Canastra* (Canastra Range); SE = Southern Espinhaço; NE = Northern Espinhaço; SDTFs = seasonally dry tropical forests

The astonishing plant diversity in the CR has often been attributed to a species-pump mechanism driven by Plio-Pleistocene climatic fluctuations (Vasconcelos et al., 2020; Giulietti et al., 1997; Harley, 1988). Particularly during the Pleistocene, the alternation between cooler and drier conditions on glacial and warmer and wetter climates on interglacial periods potentially promoted cyclic shifts in plant distributions along altitudinal gradients and episodic changes in connectivity among CR populations (Barbosa & Fernandes, 2016; Barres et al., 2019; Magri et al., 2024). Indeed, climatic dynamics during this period have influenced plant distributions and fast speciation in montane habitats more broadly (Flantua et al., 2019), and CR plant groups often bear signatures of recent diversification bursts despite examples of relatively older lineages (e.g., Alcantara et al., 2018; Rando et al., 2016; Vasconcelos et al., 2020). However, the extent to which climate-driven range expansions and contractions operated in the CR remains debated, as strong local constraints, particularly edaphic factors, may have limited dispersal and connectivity even during favorable climatic phases (Cunha-Blum et al., 2025; Oliveira et al., 2016; Rapini et al., 2021). In addition, the evolutionary trajectory of the CR flora may be further complicated by the influence of spatially adjacent biomes of varying ages on lineage and species composition (Barros-Souza & Borges, 2023; Neves et al., 2018; Rapini et al., 2021).

The CR are surrounded to the west by young savannas (the *Cerrado*), to the east and south by rainforests (the Atlantic rainforest), and to the north by ancient SDTFs (the *Caatinga*; Figure 1) (Giulietti et al., 1997). This adjacency to multiple biomes, combined with a heterogeneous gradient of environmental conditions across the CR (Silveira et al., 2016), has been acknowledged not only as a key contributing factor to outstanding levels of floristic diversity (Giulietti et al., 1997), but also as a driver of community turnover (Neves et al., 2018). In fact, woody plant communities in CR share several species with surrounding savannas (Neves et al., 2018), likely due to their similar fire-prone ecologies and mainly open vegetation physiognomies (Hughes et al., 2013). Also, phylogenetic evidence suggests a strong evolutionary relationship between the northern Espinhaço Range and the surrounding SDTFs (implicit in Souza et al., 2013), with incursions from SDTFs to the CR likely followed by surges in local speciation. However, despite the taxonomic evidence from woody communities (Neves et al. 2018), spatial patterns and drivers of plant phylogenetic composition across the CR and surrounding biomes are still elusive.

Here we assess the connectivity between ancient mountains and lowlands using the CR as a model system. To assess the extent of phylogenetic connectivity between CR and adjacent lowland biomes, we examine patterns of phylogenetic composition and their environmental drivers. Increasing evidence suggests that CR communities are evolutionarily connected to surrounding biomes–SDTFs, savannas, and rainforests–through repeated colonization events (Rapini et al., 2021; Souza et al., 2013) and taxonomic affinities (Neves et al., 2018) that reflect exchanges across the mountain-lowland gradient rather than long-term isolation. Thus, we hypothesize that (**H1**) CR communities do not share a unique evolutionary identity, but rather are segregated in distinct units sharing lineages with the surrounding biomes. Moreover, considering the widespread tendency for niche conservatism in plants (Crisp et al., 2009), the strong influence of climatic conditions on the biogeography and evolution of CR plants (Magri et al., 2024; Neves et al., 2018), and the relatively smooth altitudinal gradients between CR and surrounding lowlands (Silveira et al., 2016), we hypothesize (**H2**) that phylogenetic composition across montane-lowland gradients is primarily driven by shared climatic conditions.

## 2 Methods

To test our hypotheses, we used 1138 species from 13 angiosperm clades: *Calliandra* Benth. (Leguminosae) (52 spp.), *Cattleya* Lindl. (Orchidaceae) (30 spp.), *Chamaecrista* (L.) Moench (Leguminosae) (95 spp.), *Diplusodon* Pohl (Lythraceae) (74 spp.), *Dyckia* Schult. & Schult. f. (Bromeliaceae) (38 spp.), *Habenaria* Willd. (Orchidaceae) (107 spp.), Lychnophorinae Benth. (Asteraceae) (65 spp.), Marcetieae M.J.R. Rocha, P.J.F. Guim. & Michelang. (Melastomataceae) (56 spp.), *Mimosa* L. (Leguminosae) (203 spp.), *Myrcia* DC. (Myrtaceae) (134 spp.), *Paepalanthus* Mart. *s*.*l*. (Eriocaulaceae) (*sensu* Andrino et al., 2021) (134 spp.), Trimezieae Ravenna (Iridaceae) (42 spp.), and Velloziaceae J. Agardh (108 spp.). These groups are diverse in CR—comprising around 20% of CR diversity based on our account—encompass a variety of life forms, and span both monocots and eudicots.

Our approach included three steps that were conducted for each plant group: (1) compilation and validation of occurrence and phylogenetic data sourced from geo-referenced databases and the literature; (2) spatial analyses of phylogenetic beta diversity to assess patterns of phylogenetic composition; and (3) generalized dissimilarity modelling (GDM) to identify drivers of phylogenetic beta diversity.

### 2.1 Occurrence and spatial data

Occurrence data for each group comes from Global Biodiversity Information Facility (GBIF, 2020), Integrated Digitized Biocollections (*iDigBio*, 2024), and *species*Link (CRIA, 2020). While GBIF and iDigBio are among the largest and most comprehensive geo-referenced occurrence databases, *species*Link includes data from regional, smaller Brazilian herbaria. See Appendix 5 in Supporting Information for dataset reference numbers.

We first removed records lacking species-level identification, not based on preserved herbarium specimens, or with invalid coordinates. Problematic records were flagged for removal using ‘CoordinateCleaner’ (Zizka et al., 2019) and the following criteria: (1) within a 10,000 m radius of capital coordinates; (2) within a 1,000 m radius of country centroids; (3) with equal absolute longitude and latitude; (4) within a 100 m radius of biodiversity institutions; (5) located in oceans; and (6) with plain zero coordinates, equal latitude and longitude, or within a 0.5 m radius of the point 0/0.

Before removing duplicate records and outliers, we updated taxonomic names and corrected any typographical errors using ‘TNRS’ (Boyle et al., 2013) and WCVP (Govaerts et al., 2021). Any name with an overall TRNS score equal to or below 0.95 was manually verified and corrected based on the GBIF backbone taxonomy database (GBIF, 2020).

Lastly, we removed duplicate records and geographical outliers using ‘CoordinateCleaner’ with standard parameters. Occurrence data points and spatial polygons were projected into the SIRGAS 2000/Brazil Mercator coordinate system (EPSG:5641). Records located outside the documented distribution of the CR (Silveira et al., 2016; Vasconcelos et al., 2020) and their surrounding biomes (IBGE, 2019) were removed.

The study area was divided into hexagonal grid cells of 50 km edge-to-edge distance, and each record was assigned to its correspondent grid cell in order to build presence-absence matrices. A total of 2176 cells were created, of which 274 were CR cells. To mitigate common biases in presence-only data and avoid the artificial inflation of dissimilarities (Mokany et al., 2022), we removed species that occur in a single site only (singletons) and grid cells with a single species. This filtering step was applied to all downstream analyses, except for inter-area phylogenetic beta diversity (see below).

Each grid cell was assigned to a biome or CR subregion based on intersection tests available in the ‘sf’ package (Pebesma, 2018). Grid cells could overlap with multiple biomes, but if a grid cell intersected with any CR subregion, it was assigned exclusively to that subregion and not to any other spatial polygon. This approach ensured that CR grid cells were exclusive, while biome grid cells remained non-exclusive (Figure S1.1 for more details). In principle, this binary assignment could introduce bias by assigning non-CR taxa to CR cells (Figure S2.1). However, given the fragmented and discontinuous nature of the CR, as well as the uncertainty among published CR distribution maps (e.g., Colli-Silva et al., 2019; Silveira et al., 2016; Vasconcelos et al., 2020), a more restrictive approach would likely remove genuine CR occurrences. Therefore, our approach acts as a buffer that accommodates spatial uncertainty while remaining compatible with downstream analyses. To evaluate the robustness of this decision, we repeated all analyses using a more restrictive grid system in which records located in CR-assigned cells but falling outside CR polygons were filtered out (Appendix S2). All conclusions and key results remained consistent with those obtained under the binary assignment (Figures S2.2-S2.16).

It is important to note that biome boundaries are not strictly defined and do not correspond precisely to the spatial polygon limits used to represent them. Additionally, the spatial distribution of each biome, as defined here based on IBGE (2019), includes vegetation physiognomies that are not typically associated with that biome. For instance, altitudinal grasslands can be found within areas classified as rainforests, while gallery forests occur within savanna regions. Although such caveats are difficult to resolve with currently available data, the broad geographic extent and relative environmental homogeneity of each biome (IBGE, 2019) support their use as meaningful ecological units.

### 2.2 Phylogenetic trees

Phylogenetic trees needed for our analyses come from the most recent and comprehensively sampled phylogenetic hypotheses available (Alcantara et al., 2018; Amorim et al., 2019; Andrino et al., 2021; Batista et al., 2013, 2016; de Souza et al., 2019; Gustafsson et al., 2010; Inglis & Cavalcanti, 2018; Krapp et al., 2014; Loeuille et al., 2015; Lovo et al., 2018; Michelangeli et al., 2013; Pinangé et al., 2017; Rando et al., 2016; Rocha et al., 2016; Simon et al., 2011; Souza et al., 2013; van den Berg et al., 2009; Vasconcelos et al., 2020). Trees covered from 26% to 78% of the described taxonomic diversity across groups, with an average of 55%. Phylogenetic trees and presence-absence matrices were pruned to include only species present in both phylogeny and spatial data. Taxonomic names were corrected and updated using the same approach described above for occurrence data, but all names with an overall score below 1 (exact matching) were manually checked. As infraspecific names are not applied consistently to herbarium specimens, we conducted all analyses at the species level by collapsing infraspecific taxa into species both in spatial and phylogenetic data. Resultant duplicate tips were randomly removed from phylogenetic trees.

### 2.3 Phylogenetic beta diversity

Any scenario in which communities are not identical can be described with turnover, nestedness, or a combination of both. While nestedness reflects a non-random process promoting the orderly disaggregation of assemblages, turnover implies replacement of lineages <u>(Baselga, 2010)</u>. Because we were interested in the overall dissimilarity between communities, we quantified unpartitioned phylogenetic beta diversity rather than turnover or nestedness individually. All phylogenetic beta diversity and related analyses were performed using the R packages ‘betapart’ (Baselga & Orme, 2012) and ‘recluster’ (Dapporto et al., 2013).

#### 2.3.1 Grid-cell pairwise phylogenetic beta diversity analyses and phyloregionalization

We calculated phylogenetic beta diversity as the Sørensen derived pairwise phylogenetic dissimilarity (Koleff et al., 2003) for each grid cell pair, and the resultant distance matrices were used to conduct phyloregionalization analyses in order to test whether the CR have a cohesive evolutionary identity (**H1**). We used the Ward clustering algorithm because it performs better than other hierarchical clustering methods in phyloregionalization analyses, yielding phyloregions with a more balanced number of grid cells and the lowest possible within-cluster variance (Ringelberg et al., 2023).

### 2.3.2 Inter-area pairwise phylogenetic beta diversity analyses

To quantify phylogenetic beta diversity between CR and the different surrounding biomes, we also calculated Sørensen derived pairwise phylogenetic dissimilarity for all possible pairwise comparisons between CR subregions and surrounding biomes, each treated as a single area. Grid cells were first assigned as either CR subregions or surrounding biomes (Figure S1.1). Then, occurrence matrices were built for each area rather than at the grid-cell level. This inter-area approach was complementary to the inter-cell approach described above, and allowed us to evaluate whether lineage dissimilarity varies across different combinations of area pairs.

### 2.4 Generalized dissimilarity modelling (GDM)

Generalized dissimilarity modeling (GDM) is a powerful technique for evaluating the influence of multiple predictor variables on biological composition (e.g., taxonomic, phylogenetic, or functional) across spatial gradients (Mokany et al., 2022). Unlike other modeling approaches, GDM does not assume linear relationships between predictors and response variables, allowing for a more realistic assessment of complex ecological patterns. Moreover, GDM enables precise quantification of the contribution of individual abiotic variables to phylogenetic beta diversity.

To evaluate the influence of climate, soil, and geographic distance on the distribution of plant lineages across the CR (**H2**), we applied GDM using the ‘gdm’ R package (Mokany et al., 2022), including only pairwise phylogenetic dissimilarities with at least one CR grid cell as the response variable. This ensured that GDM results were exclusively CR-related and did not involve lowland-lowland pairwise dissimilarities. As predictors, we considered 19 bioclimatic variables from WorldClim 2.1 (Fick & Hijmans, 2017), and 9 edaphic variables from SoilGrids 2.0 (Poggio et al., 2021) averaged across 5-, 15-, and 30-cm sampling depths (Figures S3.1-S3.28).

To minimize multicollinearity, we assessed correlations among variables within the same category (precipitation, temperature, and soil) using Pearson correlation coefficient (*r*), and selected uncorrelated subsets (|*r*| ≤ 0.75). This resulted in a final set of 15 predictors (Figure S3.29). Additionally, we calculated geographic distance internally in the ‘gdm’ package as the linear Euclidean distance between grid cell centroids. Although least-cost distances are more realistic representations of geographic distance for organisms (e.g., Cai et al., 2024), we used linear distances because the study area lacks clear geographic barriers needed to define least-cost distances.

To disentangle the joint effects of variables within each category (climate, soil, and distance) on phylogenetic beta diversity, we performed partitioned GDM analyses. This involved running GDM models with all predictor categories together and each predictor category individually to assess the relative importance of each category.

## 3 Results

### 3.1 Phylogenetic beta diversity

#### 3.1.1 Grid-cell pairwise phylogenetic beta diversity analyses and phyloregionalization

Overall, phyloregionalization analyses show lack of cohesion between all the different CR subregions. Instead, communities in each subregion in general tend to be more similar to communities in the surrounding lowland biomes. The strength of these patterns vary among groups and areas.

A connection between sites in BCP, SC, SE and savannas is evident for *Mimosa*, which also demonstrates strong similarities between communities in NE and SDTFs. Similarly, lineage sharing between BCP and savannas are clear also for *Calliandra*, Lychnophorinae, and Marcetieae, for example. *Calliandra* and *Chamaecrista* reinforce the similarities between NE and SDTFs.

Patterns of lineage sharing between CR and rainforests are particularly evident for *Habenaria*, Velloziaceae, and especially *Myrcia*. For *Habenaria*, a single phyloregion includes most CR cells but also rainforest sites. This connection is also observed for Velloziaceae, although the relationship between SE and rainforests is stronger. Phyloregionalization for *Myrcia* reveals a large rainforest phyloregion including most cells of all CR subregions. However, *Myrcia* phyloregions including communities in the savannas and SDTFs also include, albeit to a lesser extent, cells from all CR subregions (especially from BCP), suggesting that the CR hosts *Myrcia* lineages from both dry and moist surrounding biomes.

Also, as expected from previous evidence pointing to strong phylogenetic structuring in CR (Barros-Souza & Borges, 2023), our results showed that communities are more similar within the boundaries of each subregion (NE, SE, SC, BCP) or a combination of them (Figure 2). This is particularly evident in *Diplusodon* and Lychnophorinae, for which distinct phyloregions align with the general spatial distribution of each CR subregion. *Diplusodon* in particular shows stepwise similarity following the distribution of the subregions to form a large CR group. However, for both *Diplusodon* and Lychnophorinae, the phyloregions including BCP also include communities from surrounding savannas and, in Lychnophorinae, groups nesting NE or SE also contain a few SDTF, rainforests, and savanna grid cells. Similarly, in *Cattleya* and Velloziaceae, SE and NE belong to distinct phyloregions, though for Velloziaceae, SE and NE also include grid cells from broader phyloregions.

**Figure 2.**
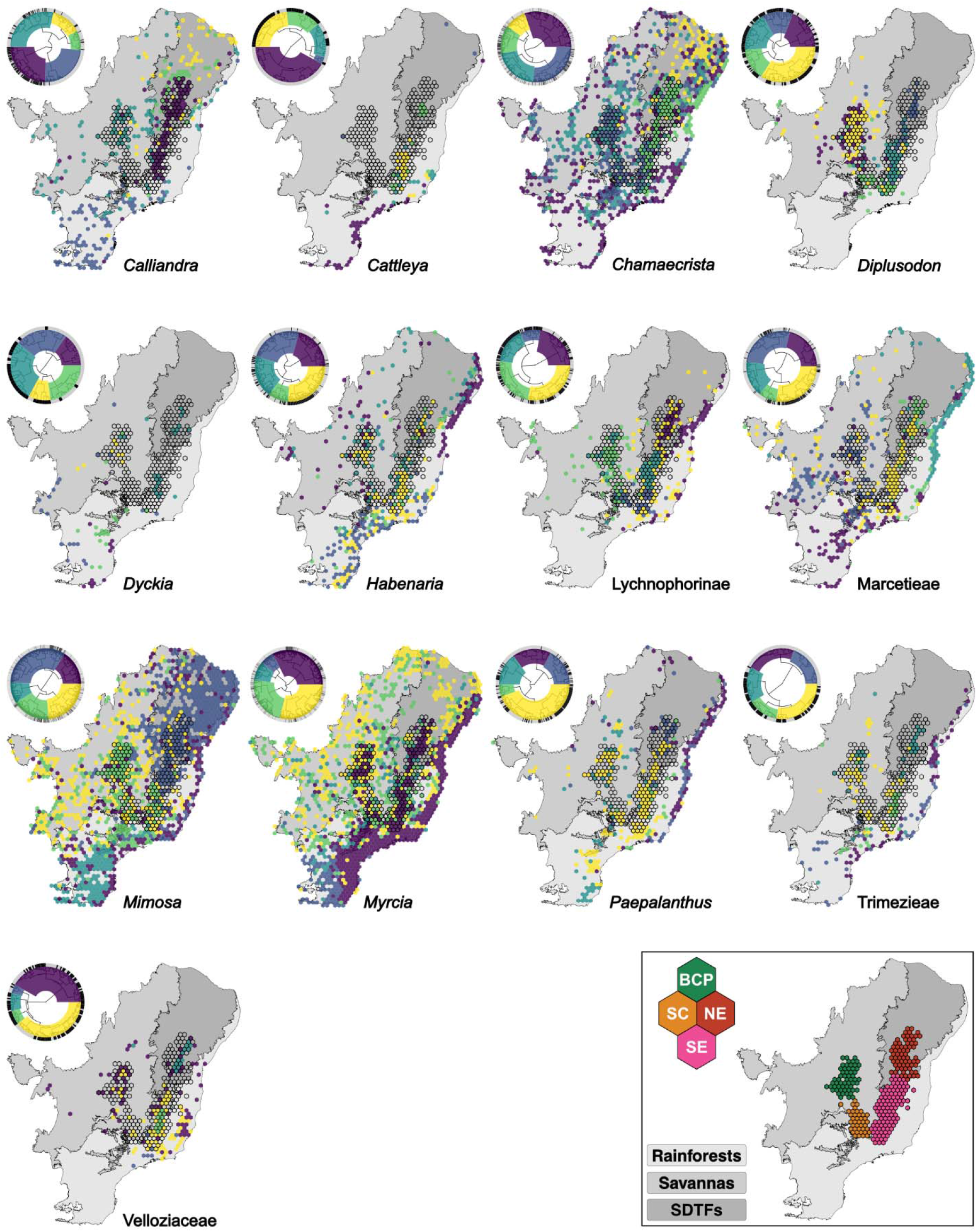
Phyloregionalization and hierarchical clustering for 13 clades in the *campos rupestres* and surrounding biomes, based on Sørensen-derived pairwise phylogenetic dissimilarity and Ward’s algorithm. In each panel, cell colors match cluster colors in the corresponding dendrogram. Tips in the dendrograms are shown in black for *campos rupestres* cells (hexagons outlined in maps) and in grey for surrounding biome cells (not outlined). Surrounding biomes are represented in the maps with different shades of grey. Uncolored cells indicate absence of occurrence data. The bottom right plot shows the distribution of *campos rupestres* subregions (colored hexagons) and surrounding biomes (colored in shades of grey). Abbreviations: BCP = Brazilian Central Plateau; SC = *Serra da Canastra* (Canastra Range); SE = Southern Espinhaço; NE = Northern Espinhaço; SDTFs = seasonally dry tropical forests

#### 3.1.2 Inter-area pairwise phylogenetic beta diversity analyses

Inter-area pairwise phylogenetic beta diversity results indicate that different CR subregions are evolutionarily linked to distinct surrounding biomes (Figure 3). Northern Espinhaço (NE) shows the lowest phylogenetic dissimilarity to SDTFs for nearly all clades (12/13), indicating greater lineage sharing between NE and SDTFs than between NE and savannas or rainforests (except for Trimezieae). Such strong evolutionary connection reflects the geographic embedding of NE within SDTFs, which has likely facilitated lineage exchanges between these areas.

**Figure 3.**
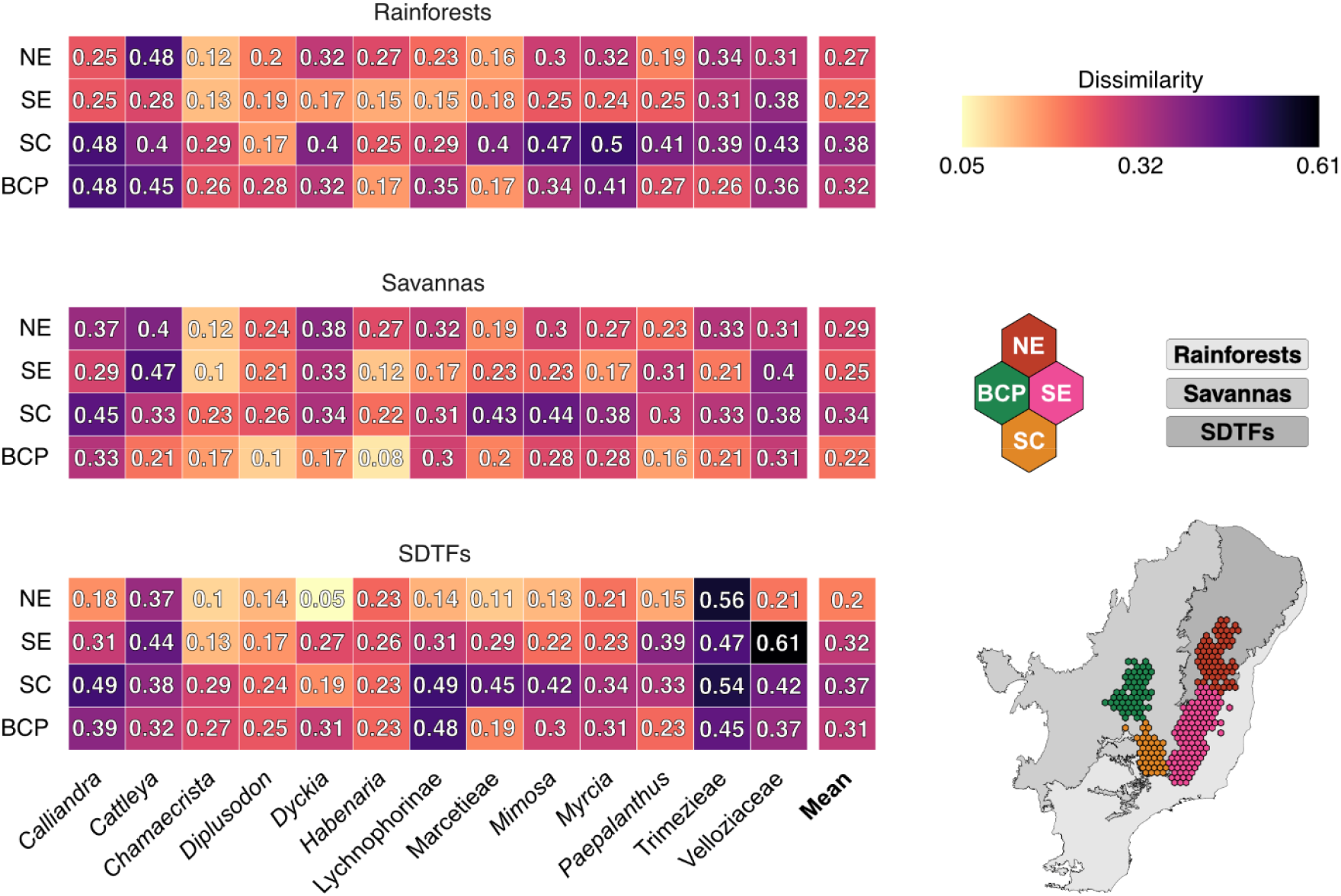
Inter-area pairwise phylogenetic beta diversity among *campos rupestres* (CR) subregions and surrounding biomes for 13 clades. **a)** Map of the study area, including *campos rupestres* subregions and surrounding biomes. Subregions are shown in different colors and by filled circles, while surrounding biomes are shown in distinct colors and by rounded rectangles. **b)** Each heatmap refers to a surrounding biome, with *campos rupestres* subregions in rows and clades (and their means) in columns. Cell values indicate lineage dissimilarity levels. Abbreviations: BCP = Brazilian Central Plateau; SC = *Serra da Canastra* (Canastra Range); SE = Southern Espinhaço; NE = Northern Espinhaço; SDTFs = seasonally dry tropical forests.

Despite forming a relatively continuous mountain chain with NE, Southern Espinhaço (SE) shows stronger evolutionary connections to rainforests or savannas. For most groups, SE shows the lowest dissimilarity with rainforests (7/13) and savannas (4/13) and the highest dissimilarity with SDTFs (7/13). Nevertheless, connections between northern SE and SDTFs are evident for some groups (e.g. *Chamaecrista* and *Mimosa*; Figures 2 and 3). These mixed patterns likely reflect the position of the SE at the intersection of all three biomes (Figure 1).

*Serra da Canastra* (SC) and the Brazilian Central Plateau (BCP) showed similar connectivity patterns, also reflecting their geographic context. As enclaves within savannas (particularly the BCP), they show the lowest dissimilarity to this biome (9/13 [SC] and 12/13 [BCP] groups). In contrast, for most groups, both subregions show higher phylogenetic dissimilarities with SDTFs (6/13 [SC] and 5/13 [BCP] groups) and with rainforests (8/13 [SC] and 7/13 [BCP] groups). Altogether, inter-area pairwise results indicate that evolutionary connections between all CR subregions and surrounding biomes largely mirror biome adjacency.

### 3.2 Drivers of phylogenetic beta diversity

Standard GDMs including climate, soil, and geographic distance explained between 6.58% (*Myrcia*) and 39.63% (*Cattleya*) of the spatial variation in phylogenetic composition (Figure 4), with a mean deviance explained of 19.17%. Partitioned GDM analyses excluding joint effects (Figure 4a) show that climate is the strongest predictor of phylogenetic dissimilarities for most groups, with relative variable importance values generally exceeding those of soil and geographic distance. Exceptions occur in *Dyckia*, in which soil is the strongest contributor, and *Paepalanthus*, where soil and geographic distance contribute equally. In Lychnophorinae, climate and geographic distance have nearly equivalent influence. When partitioning models to account for joint effects, climate remains the most influential single predictor category across groups, although soil continues to play a major role in *Paepalanthus* and *Dyckia* (Figure 4b; Table S4.1).

**Figure 4.**
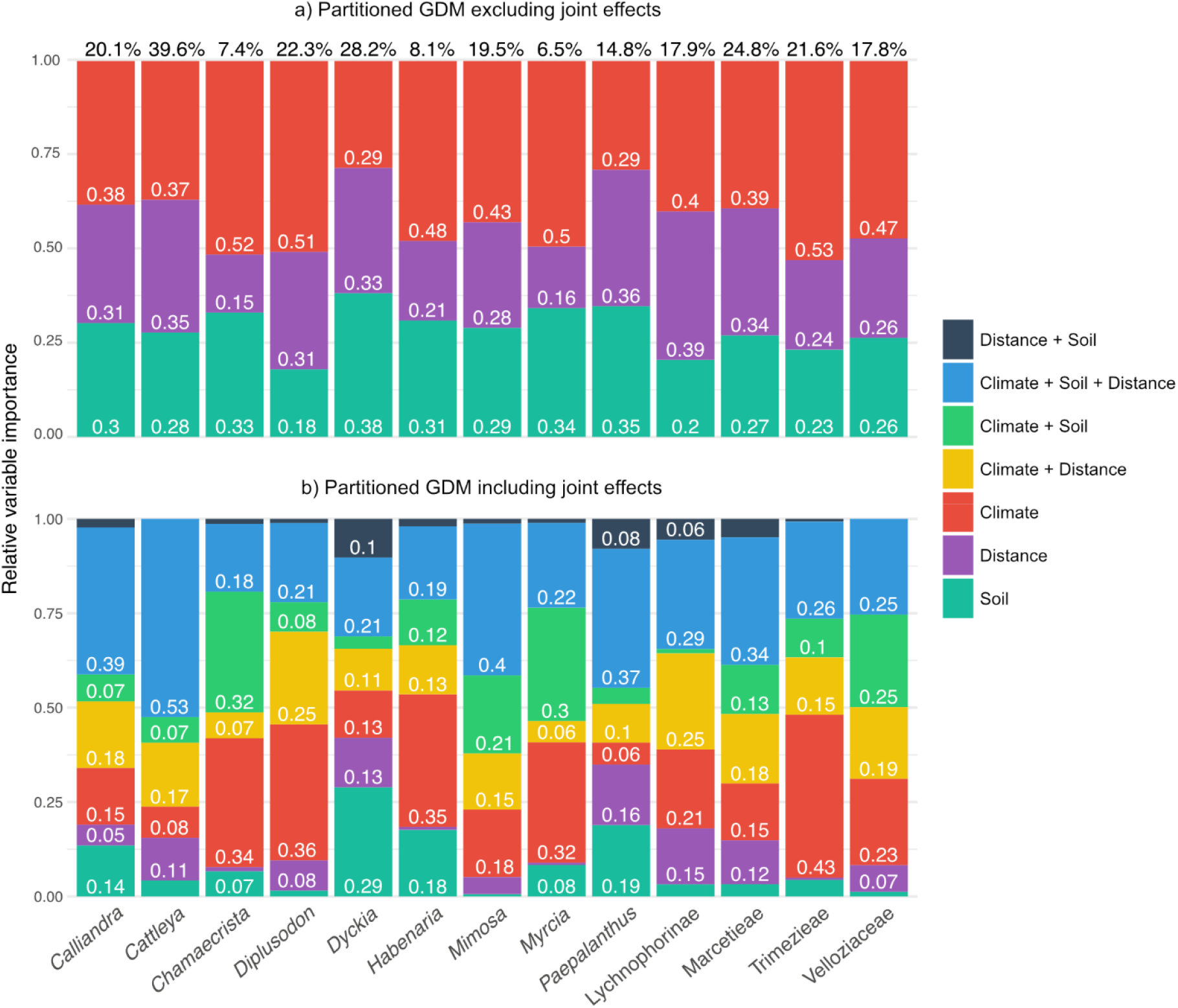
Generalized dissimilarity models (GDM) showing partitioned and total explained variance for 13 clades. Each stacked barplot corresponds to one clade, with numbers within bars indicating the relative fraction of phylogenetic beta diversity explained by each predictor category (partition). Only fractions larger than 0.05 are shown. **a)** Partitions excluding joint effects and showing the contributions of soil, geographic distance, and climate. **b)** Partitions include joint effects among predictor categories. Percentages above the bar in panel **a** indicate the total proportion of phylogenetic beta diversity explained by the full model.

At the level of individual abiotic factors, the most influential variables in each model, based on sums of coefficients, varied among plant groups. Phylogenetic dissimilarity was most strongly predicted by silt content in *Calliandra, Chamaecrista* and *Paepalanthus*, coarse fragments content in *Cattleya, Dyckia, Habenaria*, Lychnophorinae, and *Trimezieae*, precipitation of wettest quarter in *Diplusodon*, and annual mean temperature in Marcetieae, *Mimosa*, and Velloziaceae. No single abiotic factor consistently ranked among the top three drivers across all groups. However, geographic distance and coarse fragments content were among the three most important factors in 9 and 8 groups respectively (Table S4.2).

## 4 Discussion

Here we assessed the abiotic drivers and extent of evolutionary connectivity between montane ecosystems and their surrounding lowlands, using 13 angiosperm lineages that represent ca. 20% of the flora of one of the most diverse vegetation types in the Neotropics–the *campos rupestres* (CR). Our results show that phylogenetic dissimilarity is consistently low between CR subregions and surrounding biomes, suggesting that transitions between montane and lowland areas have been common across the evolution of different CR lineages (Figures 2 and 3). At the same time, CR subregions have different degrees of phylogenetic similarity with lowland biomes, which are in general linked to patterns of biome adjacency. Remarkably, Northern Espinhaço (NE) and Brazilian Central Plateau (BCP) show the highest similarities with biomes in which they are completely embedded–SDTFs and savannas, respectively–whereas Southern Espinhaço (SE), which mostly lays in between savannas and rainforests, in general shows stronger connections with these two biomes (Figures 2 and 3). Also, climate was consistently the strongest driver of phylogenetic beta diversity, suggesting that precipitation and temperature are the main factors shaping patterns of lineage sharing between CR and lowland biomes seen here (Figure 4). Soil and geographic distance play a secondary role for most groups (except for *Dyckia* and *Paepalanthus*), showing varying degrees of relative variable importance among different lineages (Figure 4).

### 4.1 Highland-lowland connections

Mountains are often described as sky-islands due to their relative isolation and limited permeability to surrounding lowlands (e.g., Barres et al., 2019). Nonetheless, isolation is not static. Sea-level fluctuations change the connectivity between islands, whereas in mountains, climatic oscillations dictate cycles of fragmentation and reconnection (Flantua et al., 2019). While elevation-driven isolation can promote high diversification and endemism (Hughes & Atchison, 2015), the sky-island analogy oversimplifies the dynamic nature of montane isolation. Unlike islands, which have sharply defined boundaries, mountains exhibit gradual environmental transitions and, thus, diffuse boundaries (Flantua et al., 2020). Additionally, erosion progressively reduces slope steepness, further blurring montane-lowland boundaries over time (Antonelli et al., 2018). These effects are particularly pronounced in ancient mountain systems (Antonelli et al., 2018) such as those harboring the CR (Alkmim, 2012).

Here we show that, across evolutionary time, CR plant communities are strongly connected to surrounding lowland biomes, supporting **H1**. Consistent with their geography (Figure 1), different CR subregions align primarily with their neighboring biomes. The BCP is strongly associated with savannas, corroborating taxonomic bioregionalization (Colli-Silva et al., 2019). In contrast, links between NE and SDTFs not detected in previous taxonomic analyses, are a novel phylogenetic pattern highlighted here. This suggests that, although microendemism at the species level is highly prevalent in many CR subregions (Giulietti et al., 1997), endemism in the NE appears less marked when considering deeper lineages, which are connected to surrounding SDTFs, as also evidenced from phylogenetic inferences (Souza et al., 2013).

Connections between CR, savannas, and SDTFs are consistent with expectations based on the sharing of a dry or mesic environment. Yet our results show that montane-lowland relationships are not restricted to dry formations. Phyloregionalization and inter-area phylogenetic beta-diversity in *Myrcia, Habenaria*, and Velloziaceae show extensive lineage sharing between CR and the Atlantic rainforest. Although explicit tests are needed, such connections likely reflect the composition of gallery forests interspersed within the CR (Nicholls et al., 2025; Silveira et al., 2016) or open vegetation embedded in the Atlantic rainforest, such as inselbergs (Cabral et al., 2021). Together, these findings support a model in which CR are not isolated “sky-islands”, but part of a broader evolutionary montane-lowland mosaic. Explicitly linking functional traits and niche evolution in analysis of trait-niche correlated evolution (e.g., Velásquez-Puentes et al., 2023) could shed light on mechanisms behind the connections between CR and surrounding environments.

By providing strong phylogenetic evidence for evolutionary connections between montane and lowland communities, we question the “sky-island” analogy for the CR. Instead, we expand the idea that CR communities are closely linked to surrounding biomes (Neves et al., 2018). The connections we unveiled encompass both dry and wet biomes, operate across evolutionary timescales, and are widespread among taxa with different life forms. Although our taxonomic sampling spans multiple lineages, future research including major CR plant groups not included here (e.g., Poaceae Barnhart, Xyridaceae C.Agardh, Rubiaceae Juss., Lamiaceae Martinov; Silveira et al., 2016) as well as denser sampling within species-rich clades, will be essential to test our generalizations. Nonetheless, the geological context of the CR (Alkmim, 2012; Saadi, 1995) further supports our results. Unlike younger montane systems such as the Andes, protracted erosion in these ancient mountains blurs the limits between montane and lowland biotas. This long-term connectivity invites the question of which environmental gradients mediate lineage exchange across the montane-lowland mosaic of eastern South America.

### 4.2 Drivers of compositional change

Evidence across different taxonomic groups and biogeographic regions consistently highlights the role of precipitation and temperature in structuring species and lineage composition (Naidoo et al., 2022; Ringelberg et al., 2023). Water availability and temperature extremes, which impose strong constraints on plant growth and reproduction (Segovia et al., 2020), sort and limit the establishment and persistence of lineages across environmental gradients. Although dispersal limitations also constrain distributions, dispersal is often more common than rapid niche evolution, as illustrated by evolutionary connections between similar environments in different continents (e.g., Morley, 2003; Renner, 2004). The widespread tendency of plant lineages to retain ancestral biome preferences during range expansions (i.e., high phylogenetic niche conservatism; Crisp et al., 2009) may shape both species distributions and phylogenetic composition of communities.

In line with global patterns, our findings suggest that phylogenetic conservatism of climatic niche explains the affinities between CR and adjacent lowland biomes, supporting **H2**. Water availability is one of the key selective pressures in the CR (Conceição et al., 2016), particularly during the dry season and in rocky outcrops, where soils are shallow and retain little moisture (Oliveira et al., 2016). Drought-tolerant species are well documented in the CR (Conceição et al., 2016; Oliveira et al., 2016; Teodoro et al., 2021), and the ability to persist under desiccation is arguably a key feature of their assemblages, even at the cost of reduced productivity (Teodoro et al., 2021). This is consistent with our analyses showing that climatic variables accounted for the largest share of variation in lineage composition for most plant groups (Figure 4). Such climate-driven variation is particularly relevant in a landscape where strong environmental gradients coincide with biogeographic boundaries. The CR spans a wide climatic range that mirrors the contrasting conditions of their dry (SDTFs), mesic (savannas), and wet (rainforests) adjacent biomes. Accordingly, we detected stronger evolutionary affinities between NE and SDTFs, between BCP/SC and savannas, and between SE and savannas/rainforests (Figures 2 and 3). As a whole, we show that climatic filters constrain lineage exchange across the mountain-lowland mosaic in eastern South America, reinforcing the role of niche conservatism in shaping biogeographic connectivity.

Geographic distance, by contrast, played a weaker role than expected. Although it was one of the strongest predictors for a few groups (e.g., *Cattleya*, Lychnophorinae, and *Paepalanthus*; Figure 4), its overall influence was modest compared to climate. Considering that autochory (dispersal not mediated by external agents) predominates in CR plants (Conceição et al., 2016) and that only ∼2% of species are shared among subregions (Colli-Silva et al., 2019), composition similarity should strongly decay as geographic distance increases. However, the relatively small contribution of distance suggests that dispersal may have been frequent enough over evolutionary timescales.

Soil emerged as the least influential driver for most groups (except for *Dyckia* and *Paepalanthus*), even though edaphic features have been shown to structure CR communities at local scales (e.g., Cunha-Blum et al., 2025; Zappi et al., 2017). Indeed, according to the OCBIL hypothesis (old, climatically buffered, infertile landscapes; Hopper, 2009), the infertile landscapes of the CR (Silveira et al., 2016) should promote the evolution of specializations in nutrient acquisition and the emergence of edaphic endemics. While such species are well documented (e.g., Oliveira et al., 2015), our findings suggest that edaphic transitions have perhaps been more frequent in evolutionary history than previously assumed (Rapini et al., 2021), leading to closely related lineages differing in soil preferences. Nonetheless, soil coarse fragments appear as a strong predictor of phylogenetic dissimilarity for many groups. As a whole, our results show that soil contrasts between assemblages may not be strong enough to constrain phylogenetic composition at the broad regional and temporal scales considered here.

A potential caveat to our findings comes from the fact that climate and soil databases are interpolated products derived from heterogeneous observations (Fick & Hijmans, 2017; Poggio et al., 2021) and therefore unlikely to capture microclimatic or fine-scale edaphic conditions. Although this limitation is inherent to large-scale biogeographical analyses, such as the ones presented here, refinement of climate and soil databases are paramount to test our conclusions that climate, not soil, is the major abiotic factor structuring phylogenetic community composition across the CR and adjacent biomes.

### 4.3 Evolutionary identity of an ancient mountain vegetation

Although recognized as a major center of plant diversity and endemism (Colli-Silva et al., 2019; Giulietti et al., 1997), and one of the most singular and important Neotropical ecosystems (Hughes et al., 2013), the CR remain underexplored, with few studies addressing their deep-time biogeographic and evolutionary dynamics (Alcantara et al., 2018; Barros-Souza & Borges, 2023; Vasconcelos et al., 2020). Traditionally, models to explain the assembly of the CR have relied on their apparent isolation and on the effects of geographic barriers such as the ∼300 km lowland stretch separating the SE and NE subregions (Harley, 1988). Although disjunct distributions are evident for several plant groups, the broader phylogenetic evidence provided here shows that many CR lineages also occur in lowlands. Climate mediates phylogenetic composition more consistently than geographic or vegetational barriers, challenging the view of the CR subregions as self-contained floristic units isolated by lowland areas (Giulietti et al., 1997; Harley, 1988). Evidence from other ancient ecosystems, including the Guiana highlands where temperature and moisture shaped floristic turnover across elevations (Liu & Smith, 2021; Rull, 2004), reinforces the idea that the climate-driven mechanisms identified here may be pervasive in evolutionary connections between lowland and ancient mountains.

A second, long-standing idea frames the CR as “species-pumps” that repeatedly generated novel lineages during recent climatic oscillations and resulting isolation-reconnection cycles between CR patches. While climatic oscillations promoted flickering connections in multiple montane systems (Flantua et al., 2019; Muellner-Riehl, 2019), an alternative hypothesis proposes that much of the CR endemic diversity arises in part by lineage exchange via niche shifts to and from surrounding biomes (Rapini et al., 2021). We provide partial support to this latter idea by showing extensive phylogenetic links between montane and lowland assemblages. Moreover, different subregions interconnect via the surrounding biomes, particularly those embedded in the savannas. Our results also indicate that lineage exchange between highlands and lowlands occurred over timescales that may exceed the time frame of the species-pump hypothesis, and, importantly, that it is bounded by climatic conservatism and environmental filtering. Future research focusing on drivers of diversification rates and directionality and timing of biome transitions will help to clarify these open questions on CR evolution.

As a whole, the CR are not isolated evolutionary “islands” nor solely the arenas of Pleistocene-driven speciation pulses. Instead, they are nodes in a long-term network of connectivity shaped by climatic buffering and geological stability (Silveira et al., 2016), episodic and localized diversification bursts (Barros-Souza & Borges, 2023; Vasconcelos et al., 2020), and lineage permeability. Although recent climatic oscillations evidently contributed to CR diversity (Bonatelli et al., 2014; Dantas-Queiroz et al., 2021), we highlight an additional, deeper layer: montane-lowland evolutionary links, driven by the persistence of ancestral climatic affinities on a background of ancient, eroded landscapes with soft environmental contrasts to neighboring biomes. Based on the evidence so far, we argue that the CR evolutionary identity has many faces. They are simultaneously persistent and fragile (Fernandes et al., 2018; Silveira et al., 2016), distinctive (Colli-Silva et al., 2019) and permeable, producing a flora that is uniquely prone to pump new lineages (Vasconcelos et al., 2020) and deeply integrated with surrounding biomes. Recognizing this complexity, which balances connectivity and singularity, reframes the role of the CR as one of the key weavers of the rich eastern South American evolutionary tapestry.

## 5 Conclusion

Assessing the extent to which lineages transcend environmental and geographic barriers is critical to understanding floristic assembly across spatial and evolutionary scales. Here we show that the ancient *campos rupestres* (CR) of eastern South America have been highly permeable to adjacent lowland biomes–savannas, rainforests, and SDTFs–over macroevolutionary timescales. Climate emerged as the dominant driver of phylogenetic composition, structuring montane-lowland links, while soil and geographic distance played variable and generally minor roles. We argue that the evolutionary history of the CR reflects a long-term, multi-faceted interplay between lineage permeability and diversification within an ancient, climatically buffered landscape. Finally, we underscore the critical role of the surrounding biomes in maintaining high diversity levels in the ecosystems of ancient mountains over evolutionary time.

## Acknowledgments

We thank Renske E. Onstein, Luis L. Valente, and Jens J. Ringelberg for comments on later versions of the manuscript, and members of the Taxonomy and Evolution of Plants (UFSCar), Evolution and Adaptation (iDiv), and Biodiversity Hotspots (Naturalis Biodiversity Center) groups for discussion during development of this research. This study was financed wholly or in part by Coordenação de Aperfeiçoamento de Pessoal de Nível Superior - Brasil (CAPES) – Finance Code 001 and the São Paulo Research Foundation (FAPESP), Brasil (Process Numbers: 2023/16170-4 and 2021/13031-8 to YB; 2021/12607-3 and 2024/00265-9 to MM; 2024/20286-0 and 2023/17978-5 to RFB; and 2022/03046-0 to LMB).

## Author Contributions

YB and LMB conceived the ideas, hypotheses, and analytical approach. YB, MM, and RFB collected, compiled, and curated the data. YB performed and interpreted the analyses, designed the figures, and wrote the manuscript. LMB supervised the project, provided critical feedback throughout, and reviewed multiple versions of the manuscript. MM and RFB provided critical comments on drafts of the manuscript and contributed equally to the revision of its final version.

## Data and Code Availability Statement

The data and code that support the findings of this study are openly available at https://zenodo.org/records/18474912, reference number https://doi.org/10.5281/zenodo.18474912.

