## Supporting Information for "Beyond the Highlands: Climate Drives Evolutionary Connections Between Ancient Neotropical Mountains and Lowland Biomes"

#### Appendix 1: Grid protocol

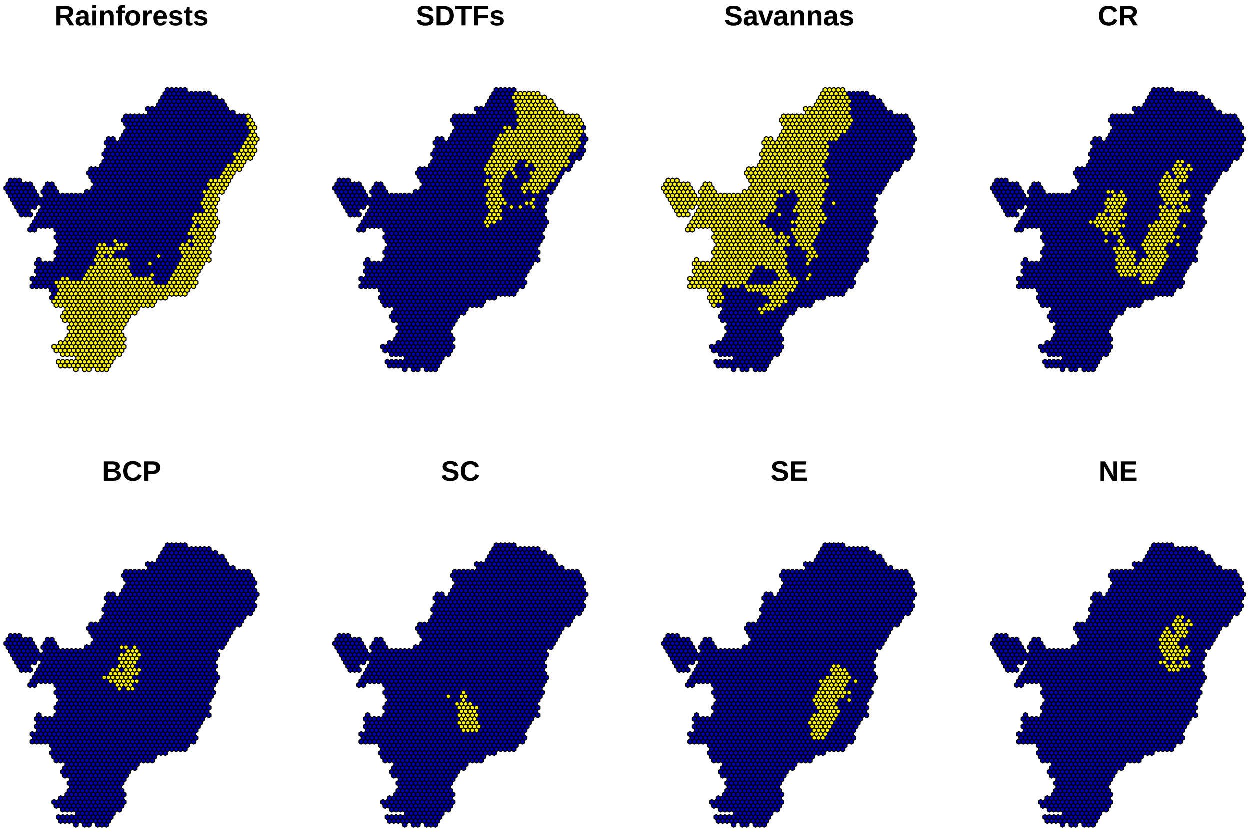


**Figure S1.1.**Grid cell system with a 50 km edge-to-edge resolution and biome/subregion assignment scheme. Cells intersecting *campos rupestres* (CR) subregions (BCP, SC, SE, and NE) were assigned exclusively to those subregions, whereas cells intersecting only biomes were assigned to one or multiple biomes. In each plot, yellow cells indicate areas assigned to the corresponding biome or CR subregion, and blue cells indicate unassigned areas. Abbreviations: SDTFs = seasonally dry tropical forests; CR = *campos rupestres*, BCP = Brazilian Central Plateau; SC = *Serra da Canastra* (Canastra Range); SE = Southern Espinhaço; NE = Northern Espinhaço.

### Appendix 2: Sensitivity analyses based on records overlapping the *campos rupestres* distribution

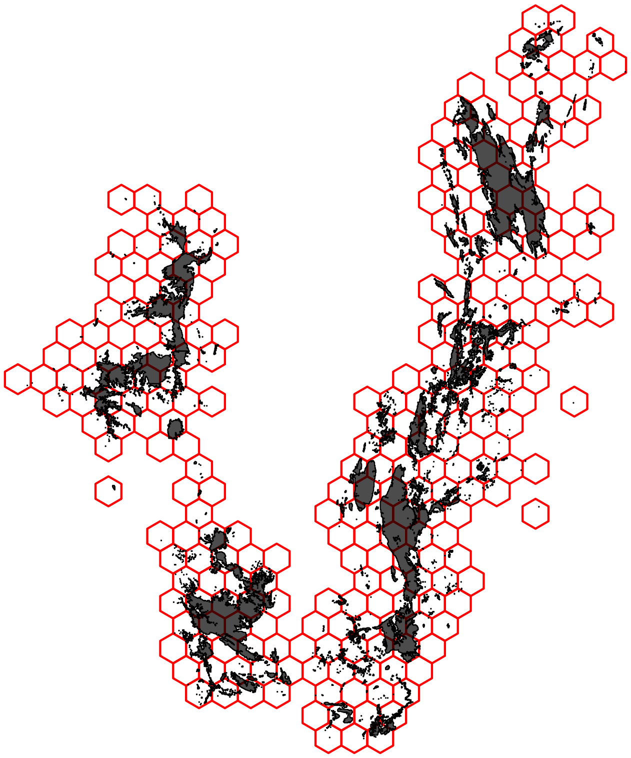

**Figure S2.1.** *Campos rupestres* (CR) documented distribution (shaded areas) overlapped with the CR assigned grid cells (red hexagons).

#### Phyloregionalization

Figure S2.2 to Figure S2.14 represents phyloregionalization and hierarchical clustering for 13 clades in the *campos rupestres* and surrounding biomes, based on Sørensen-derived pairwise phylogenetic dissimilarity and Ward's algorithm. In each panel, cell colors match cluster colors in the corresponding dendrogram. Tips in the dendrograms are shown in black for *campos rupestres* cells (hexagons outlined in maps) and in grey for surrounding biome cells (not outlined). Surrounding biomes are represented in the maps with different shades of grey. Uncolored cells indicate absence of occurrence data. The bottom right plot shows the distribution of *campos rupestres* subregions (colored hexagons) and surrounding biomes (colored in shades of grey).

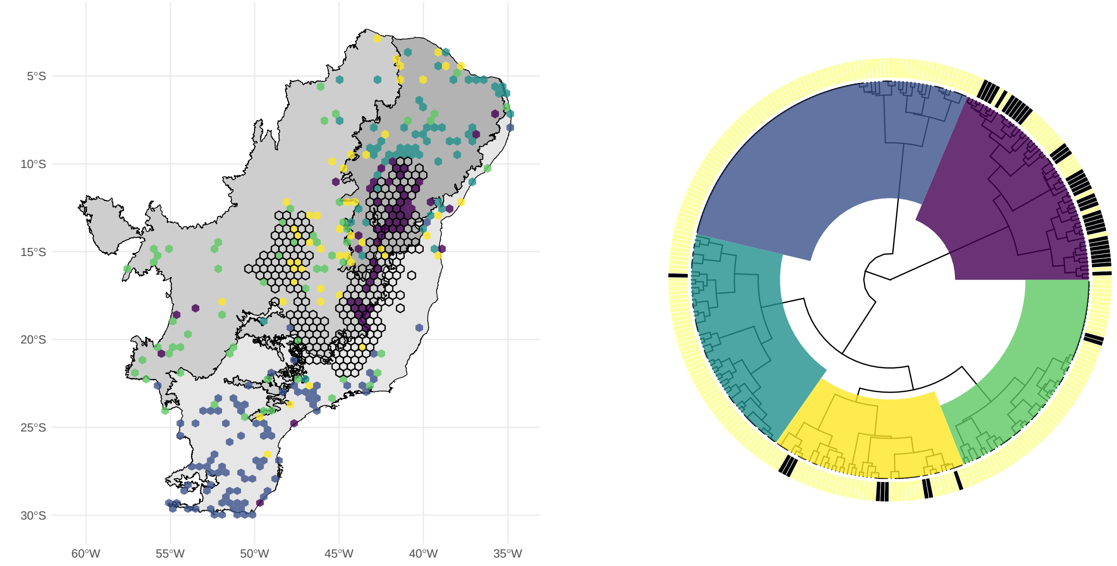


**Figure S2.2.**Phyloregionalization analysis for *Calliandra*.

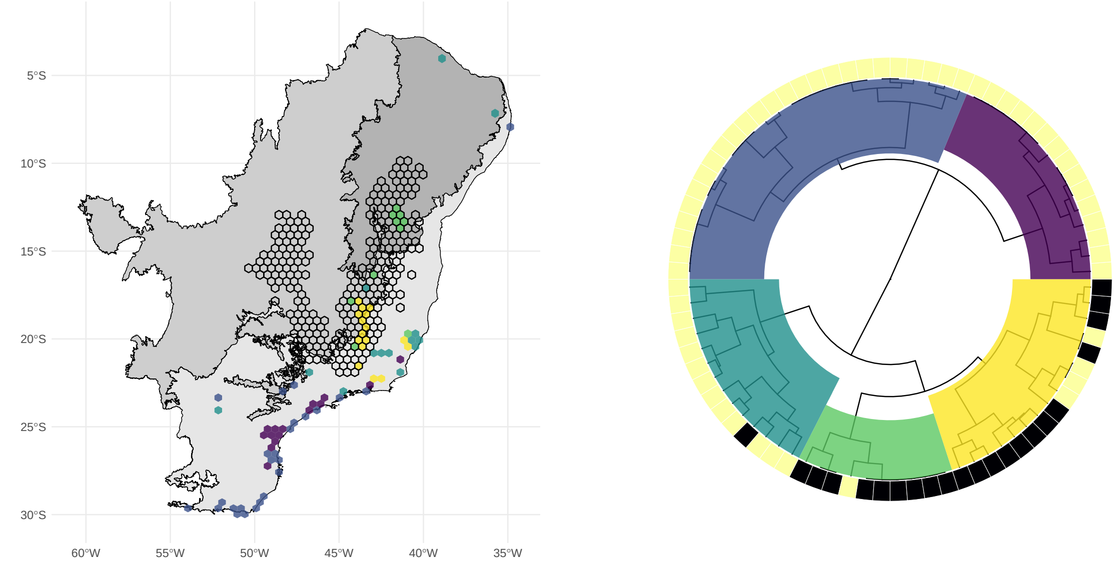


**Figure S2.3.**Phyloregionalization analysis for *Cattleya*.

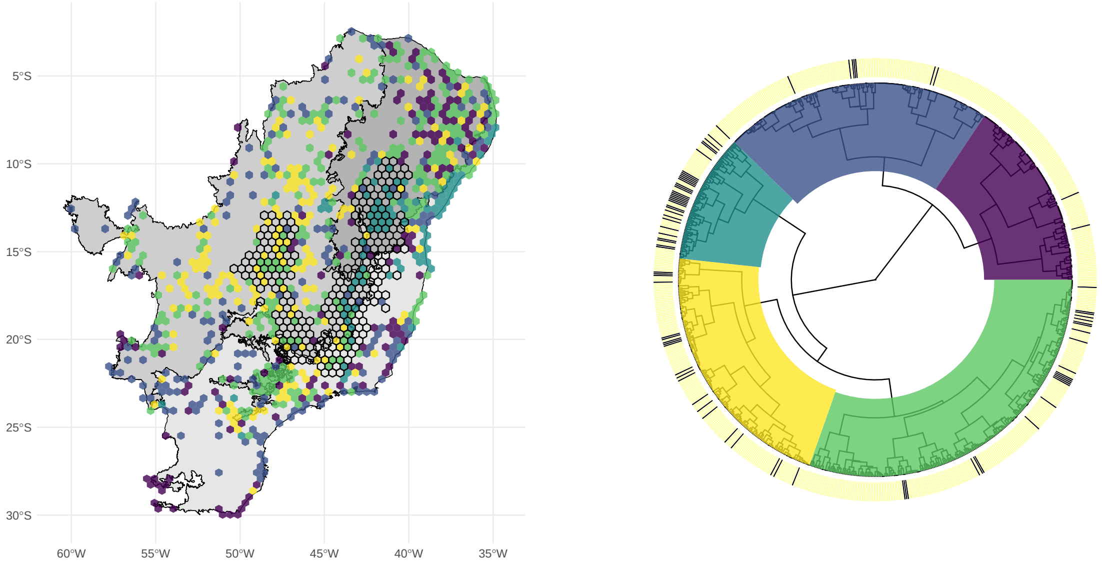


**Figure S2.4.**Phyloregionalization analysis for *Chamaecrista*.

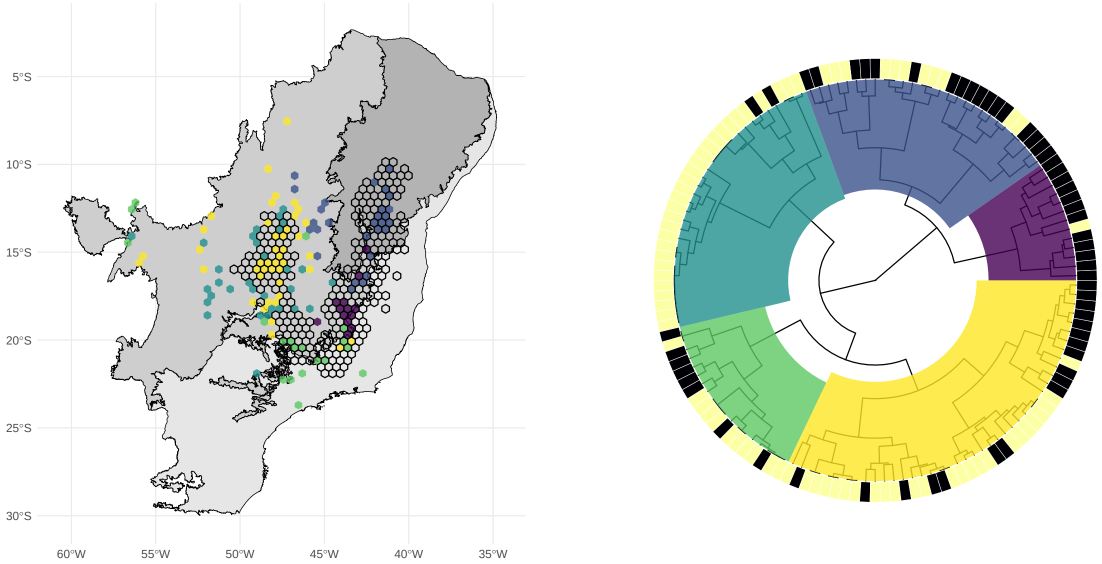


**Figure S2.5.**Phyloregionalization analysis for *Diplusodon*.

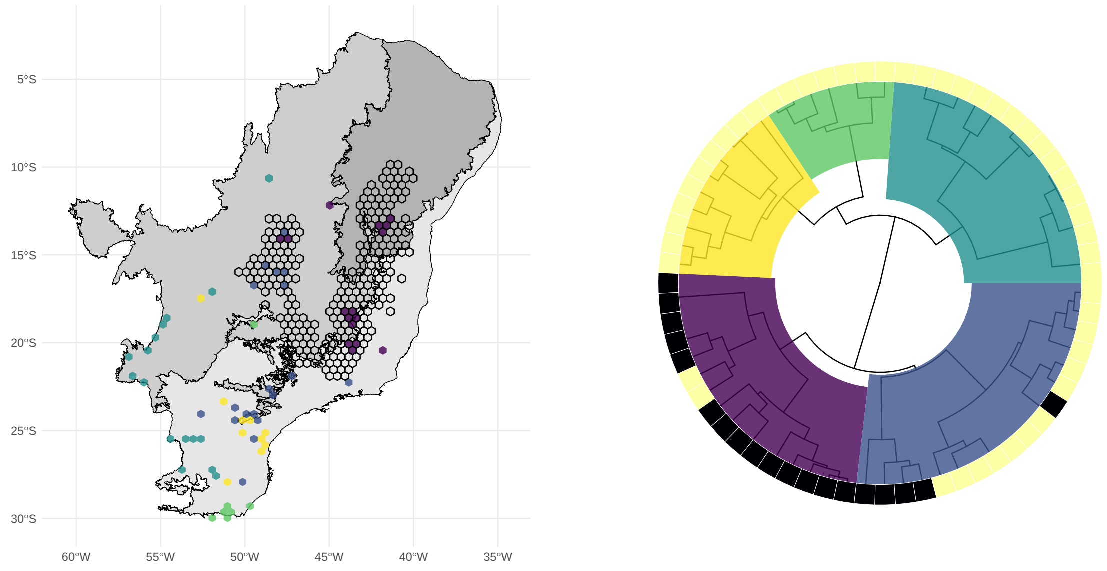


**Figure S2.6.**Phyloregionalization analysis for *Dyckia*.

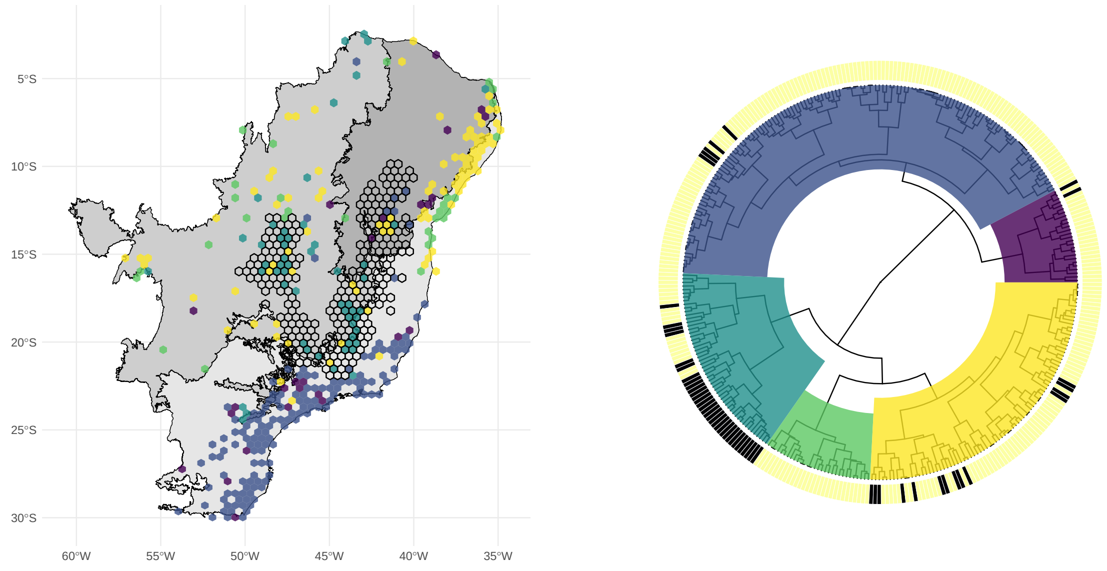


**Figure S2.7.**Phyloregionalization analysis for *Habenaria*.

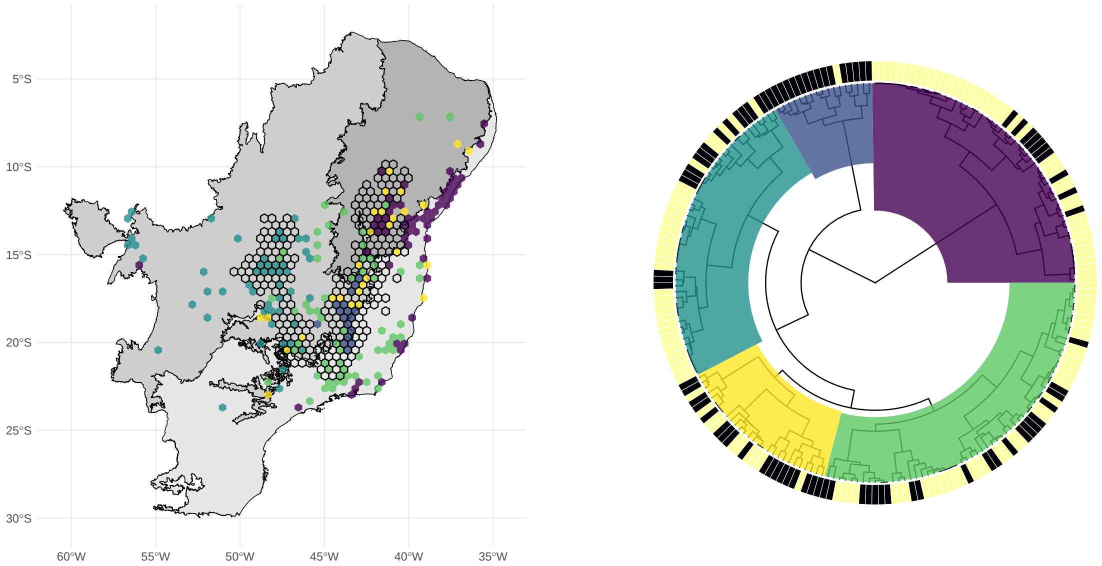


**Figure S2.8.**Phyloregionalization analysis for Lychnophorinae.

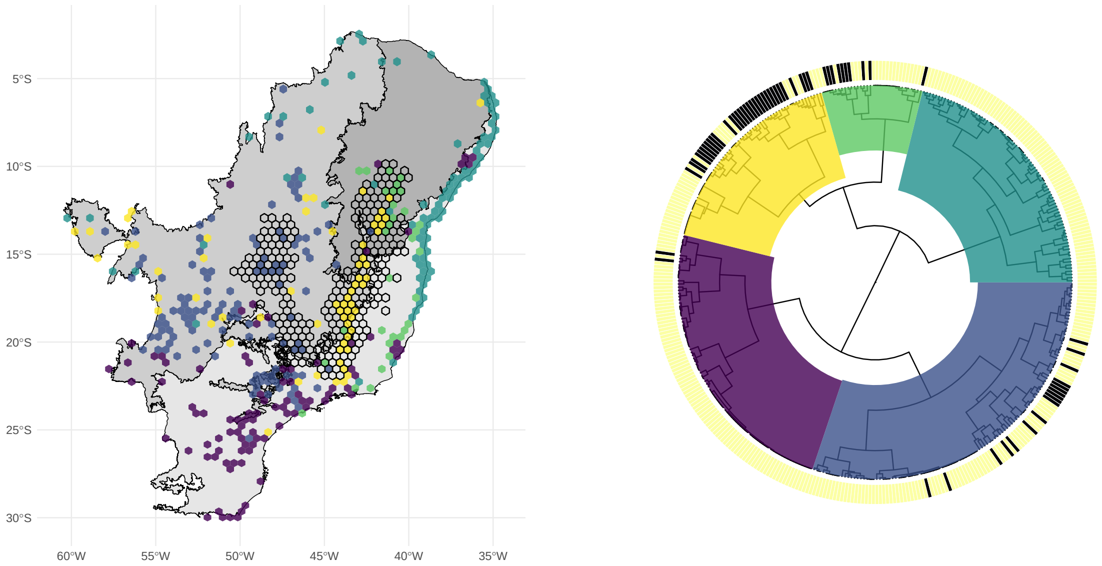


**Figure S2.9.**Phyloregionalization analysis for Marcetieae.

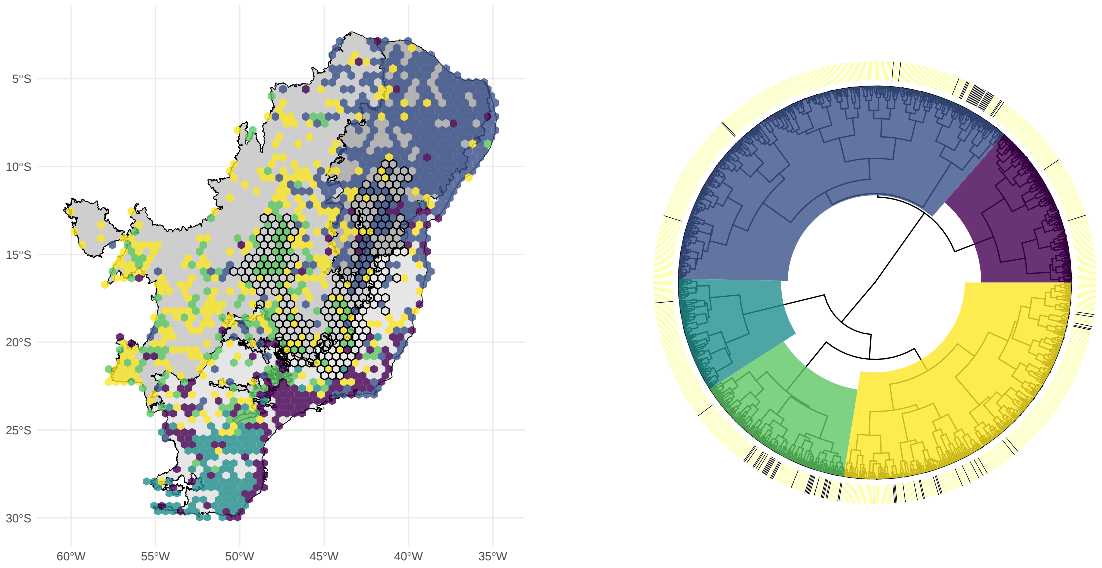


**Figure S2.10.**Phyloregionalization analysis for *Mimosa*.

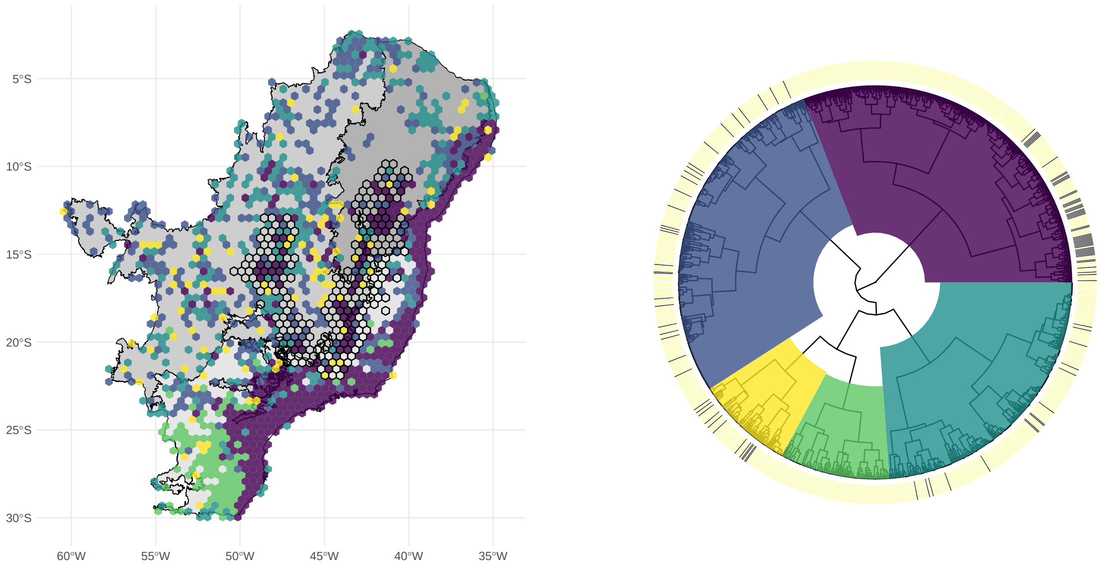


**Figure S2.11.**Phyloregionalization analysis for *Myrcia*.

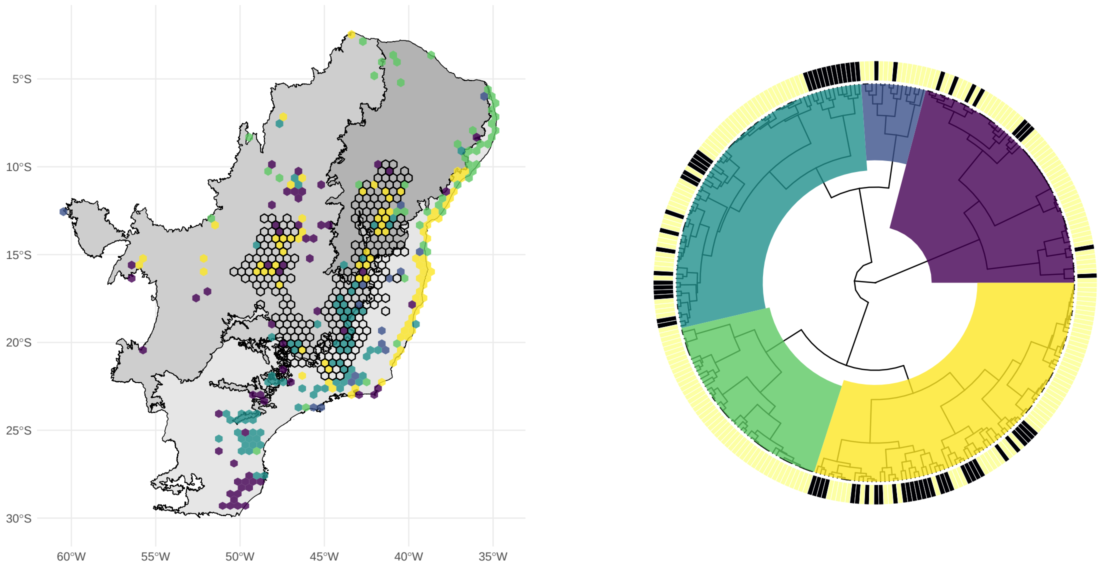


**Figure S2.12.**Phyloregionalization analysis for *Paepalanthus*.

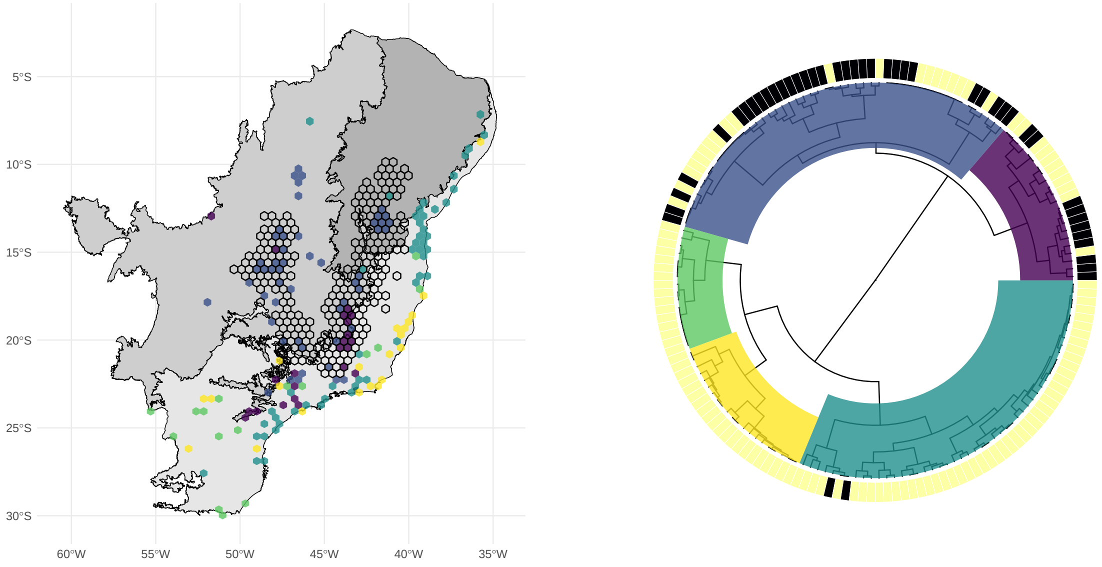


**Figure S2.13.**Phyloregionalization analysis for *Trimezieae*.

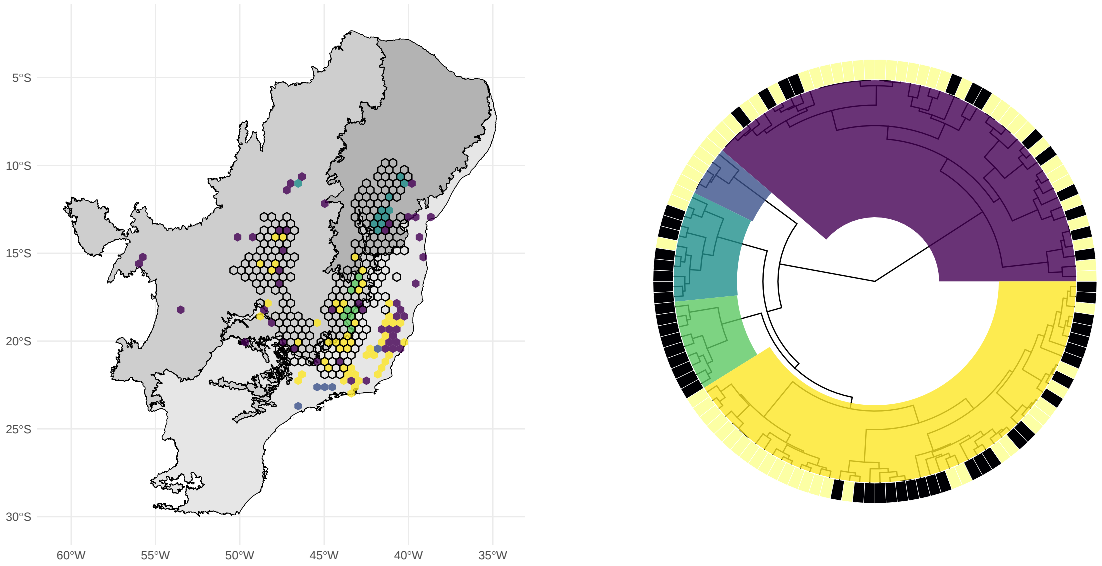


**Figure S2.14.** Phyloregionalization analysis for Velloziaceae.

#### Inter-area pairwise phylogenetic beta diversity

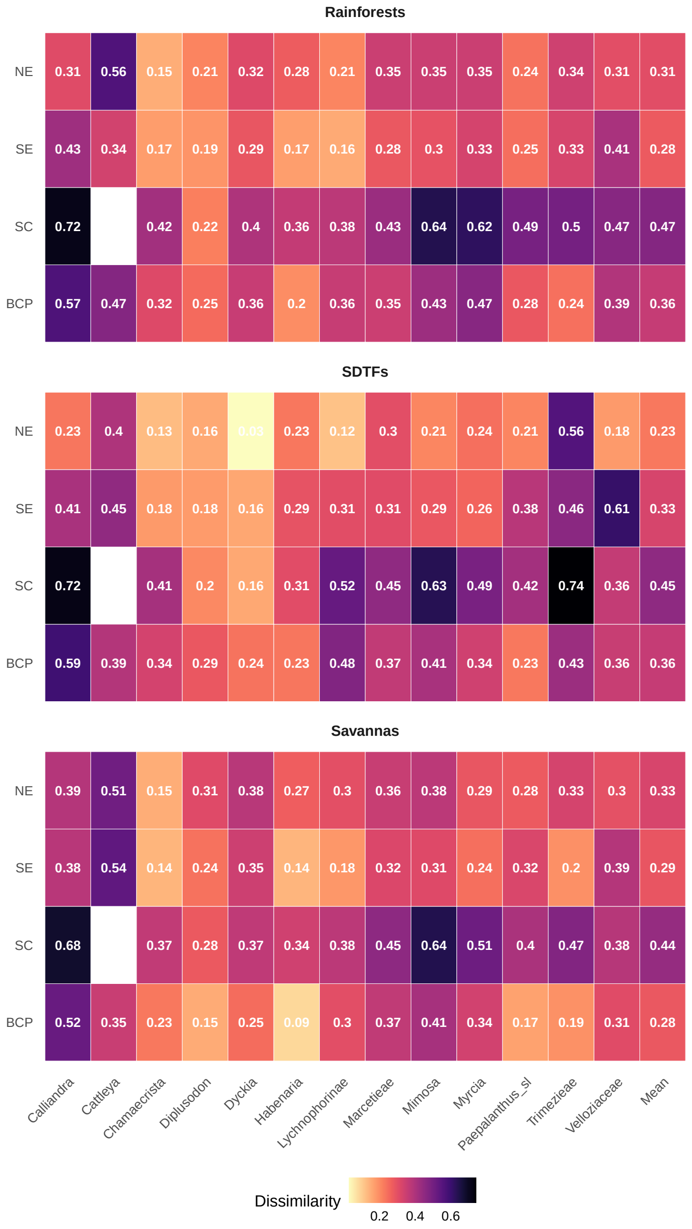


**Figure S2.15.**Inter-area pairwise phylogenetic beta diversity among *campos rupestres* (CR) subregions and surrounding biomes for 13 clades. Each heatmap refers to a surrounding biome, with CR subregions in rows and clades (and their means) in columns. Cell values indicate lineage dissimilarity levels. Abbreviations: BCP = Brazilian Central Plateau; SC = *Serra da Canastra* (Canastra Range); SE = Southern Espinhaço; NE = Northern Espinhaço; SDTFs = seasonally dry tropical forests.

**Partitioned GDM**

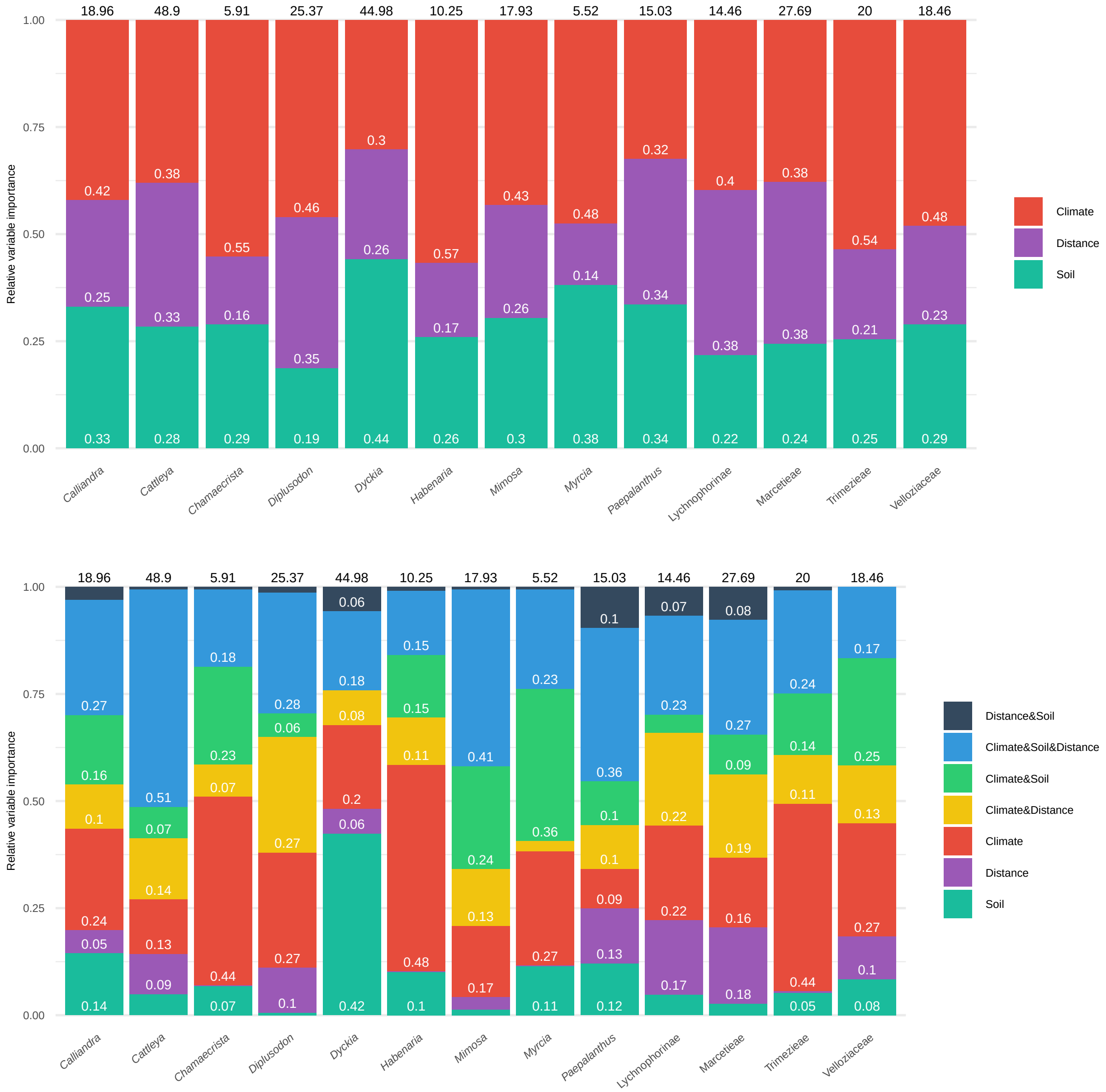


**Figure S2.16.**Generalized dissimilarity models (GDM) showing partitioned and total explained variance for 13 clades. Each stacked barplot corresponds to one clade, with numbers within bars indicating the relative fraction of phylogenetic beta diversity explained by each predictor category (partition). Only fractions larger than 0.05 are shown. Upper plot: Partitions do not account for and show the contributions of soil, geographic distance, and climate. Bottom plot: Partitions include joint effects among predictor categories. Percentages above the bars indicate the total proportion of phylogenetic beta diversity explained by the full model.

### Appendix 3: Distribution of abiotic variables and their pairwise correlations

#### Bioclimatic variables

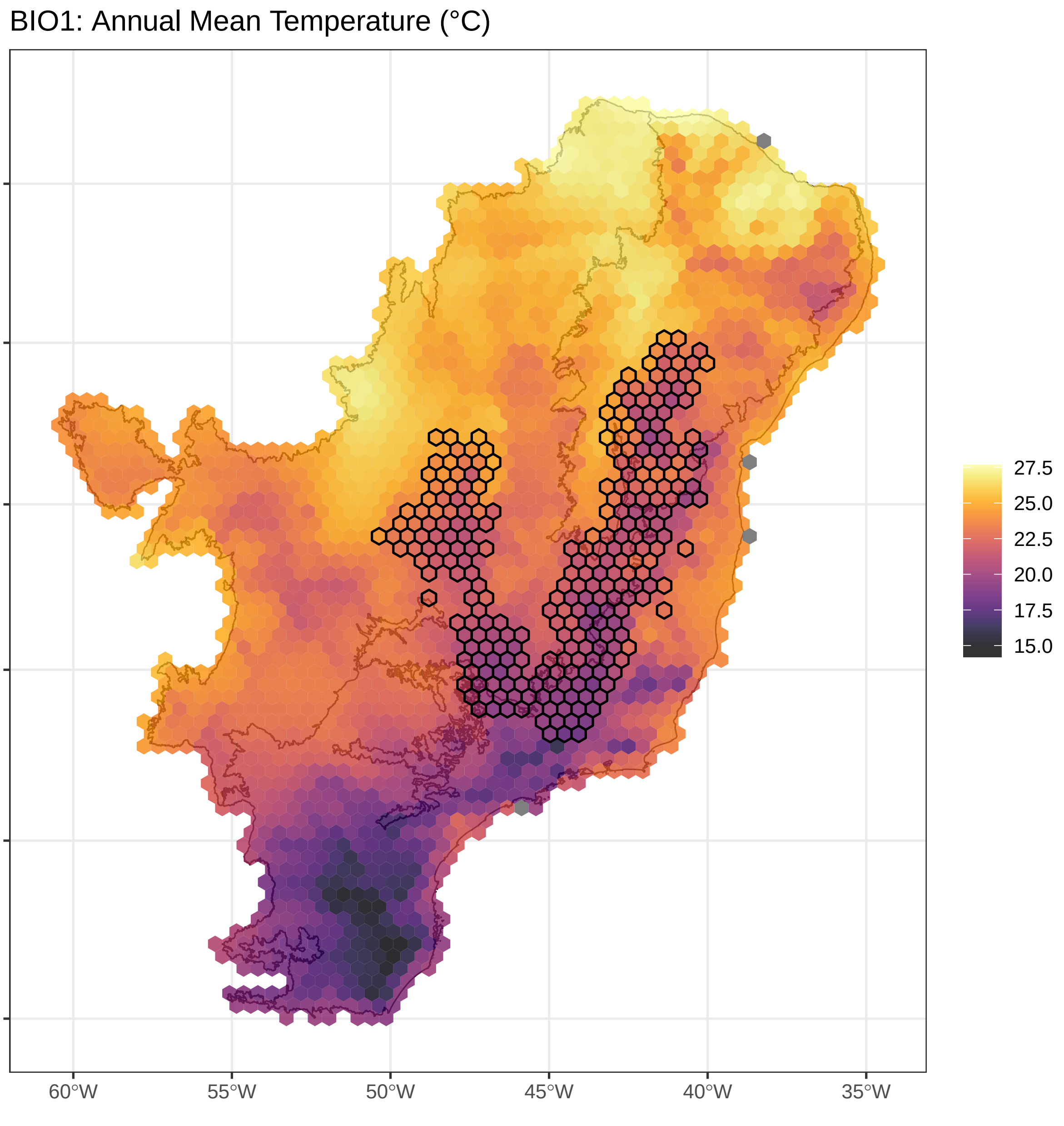


**Figure S3.1.**Annual mean temperature values assigned for each cell. Cells assigned to *campos rupestres* are outlined.

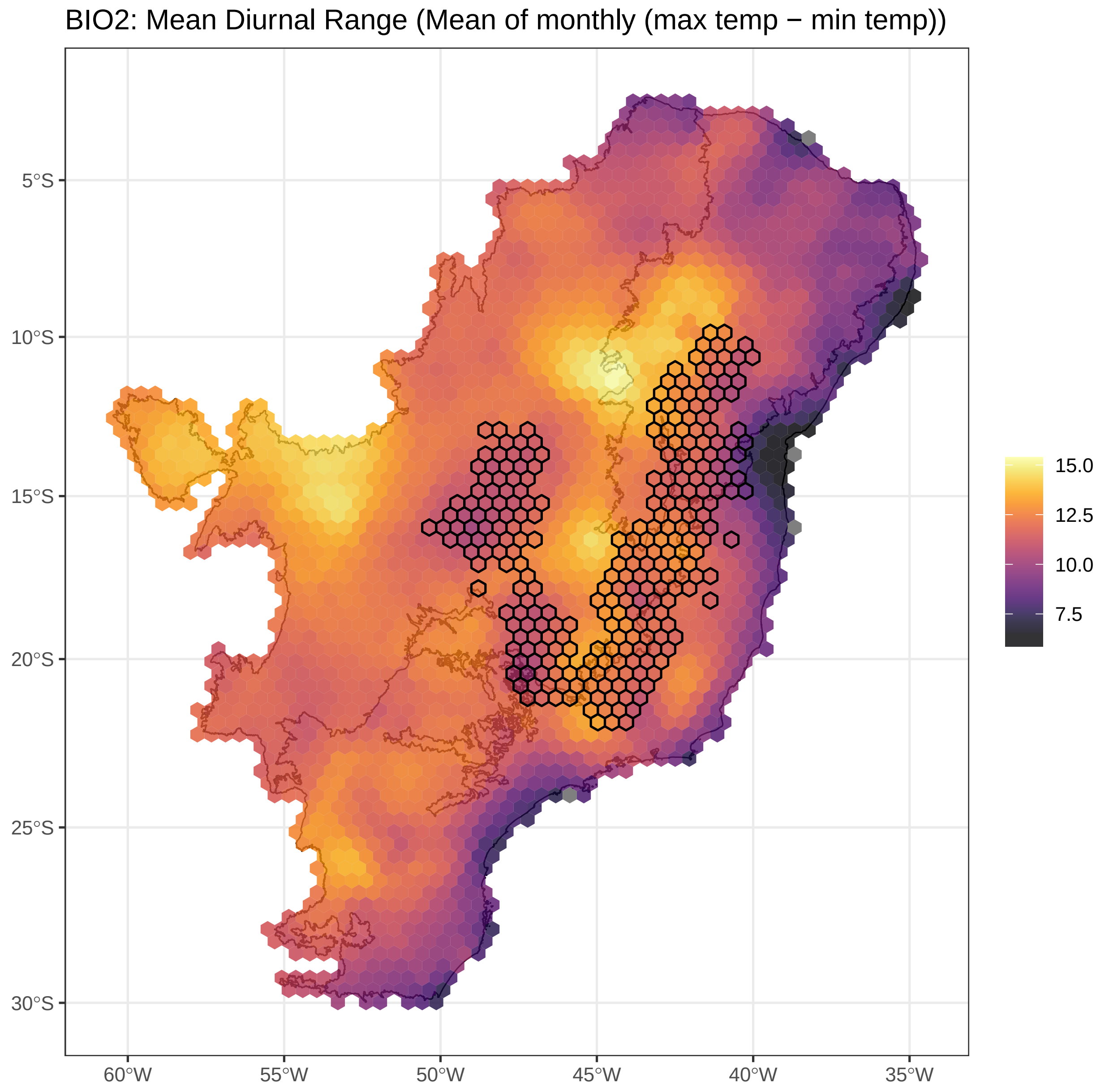


**Figure S3.2.**Mean diurnal range values assigned for each cell. Cells assigned to *campos rupestres* are outlined.

BIO3: Isothermality (BIO2/BIO7) (×100)

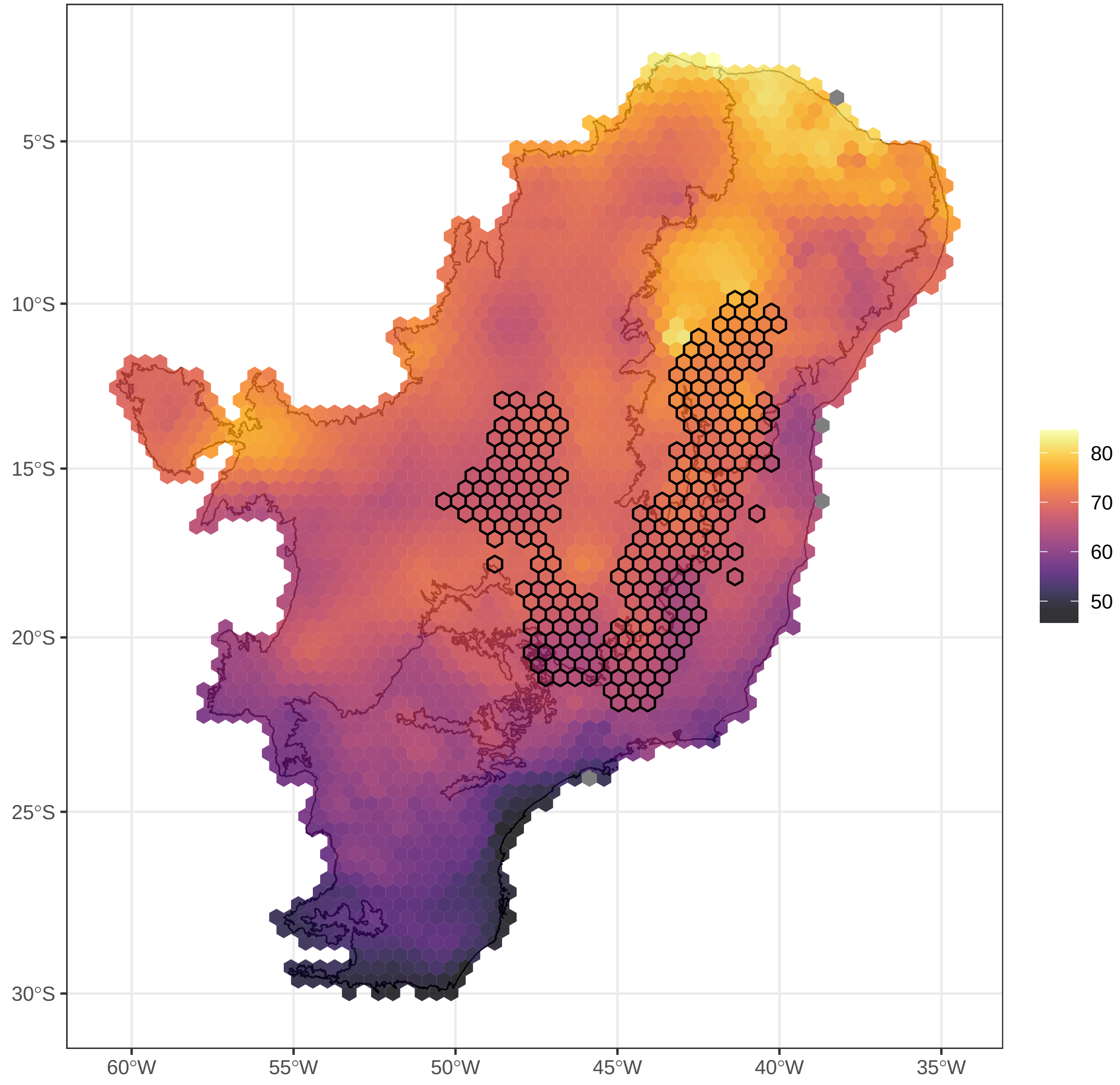


**Figure S3.3.**Isothermality values assigned for each cell. Cells assigned to *campos rupestres* are outlined.

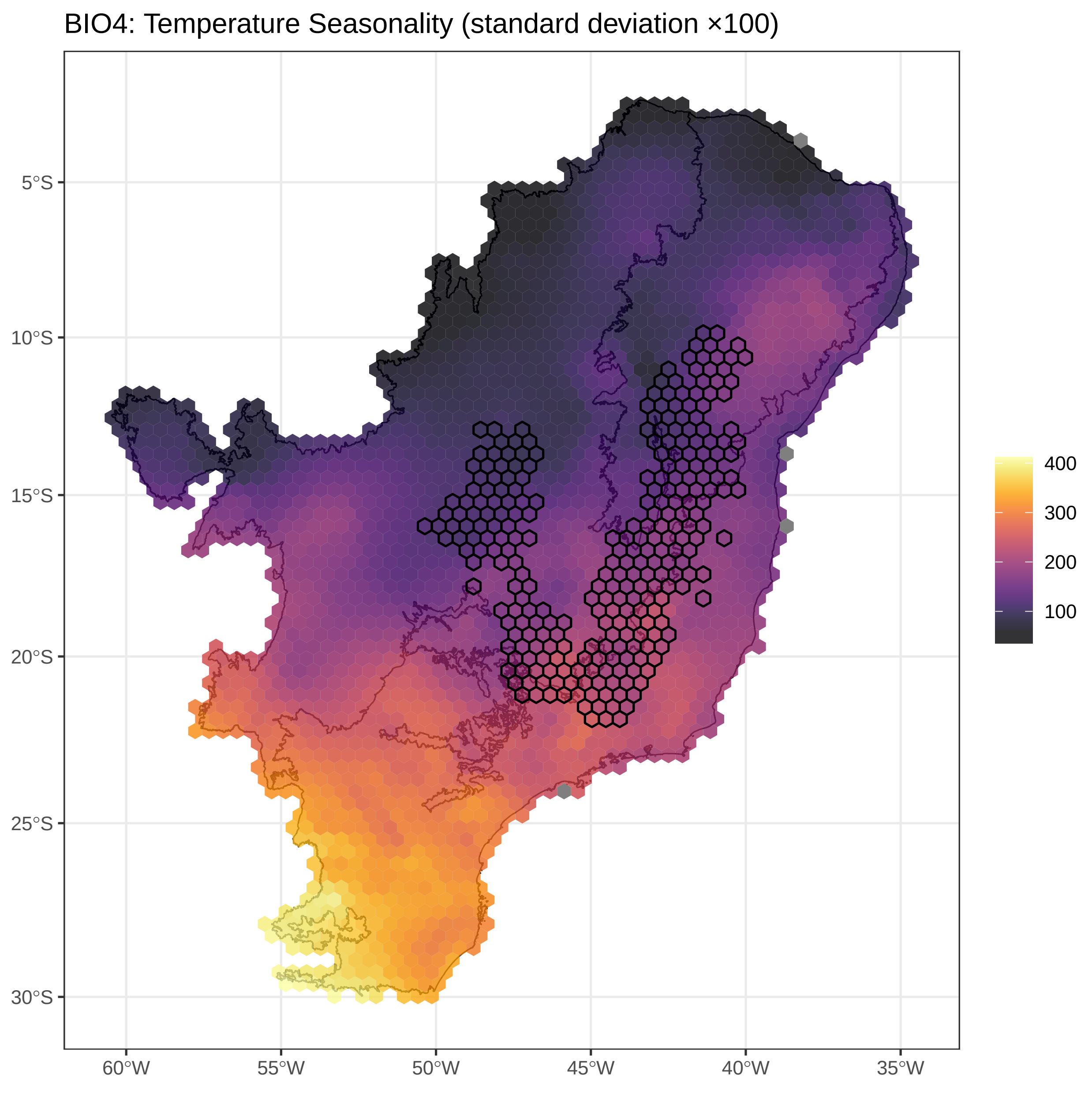


**Figure S3.4.**Temperature seasonality values assigned for each cell. Cells assigned to *campos rupestres* are outlined.

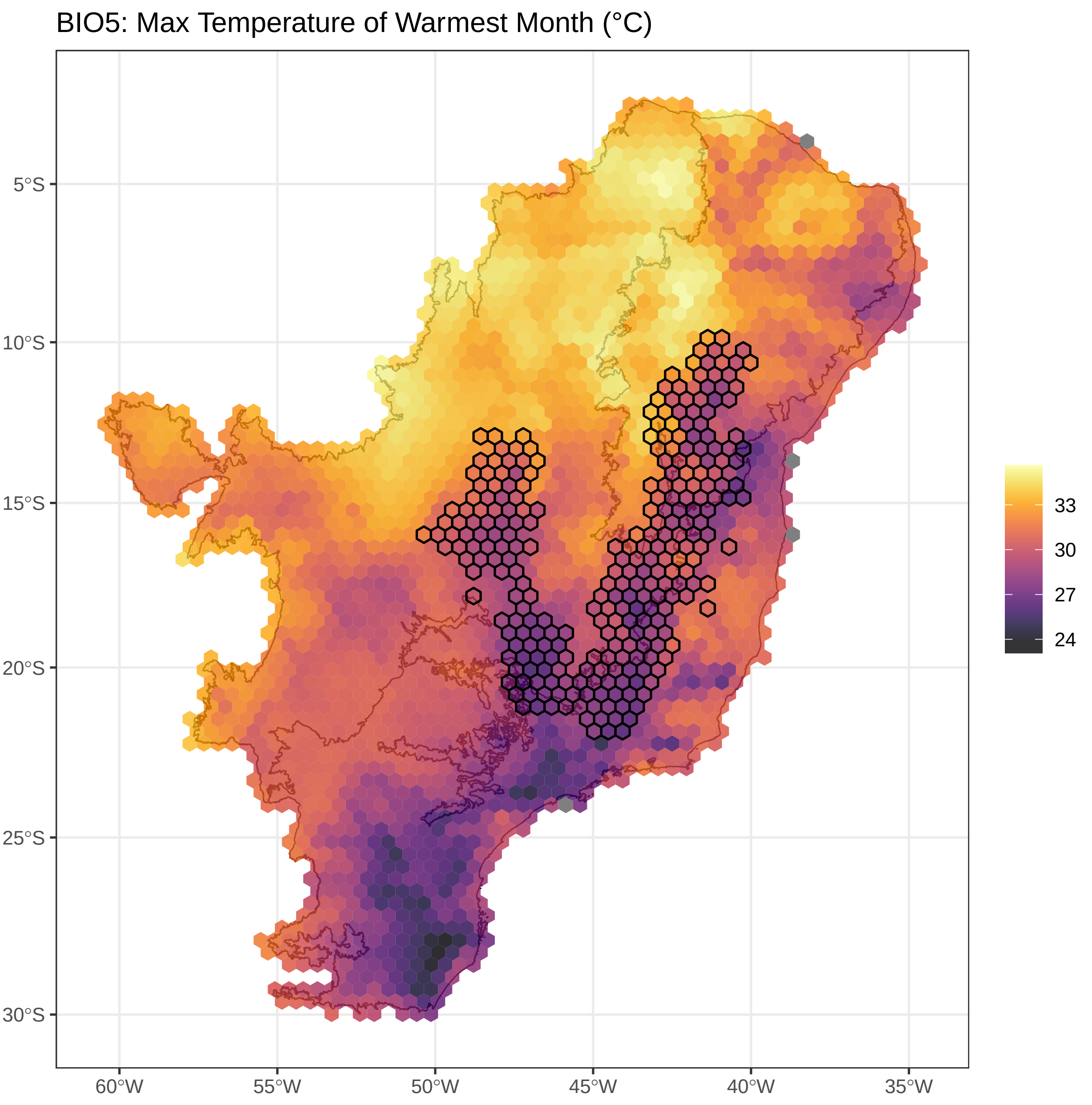


**Figure S3.5.**Maximum temperature of the warmest month values assigned for each cell. Cells assigned to *campos rupestres* are outlined.

BIO6: Min Temperature of Coldest Month (°C)

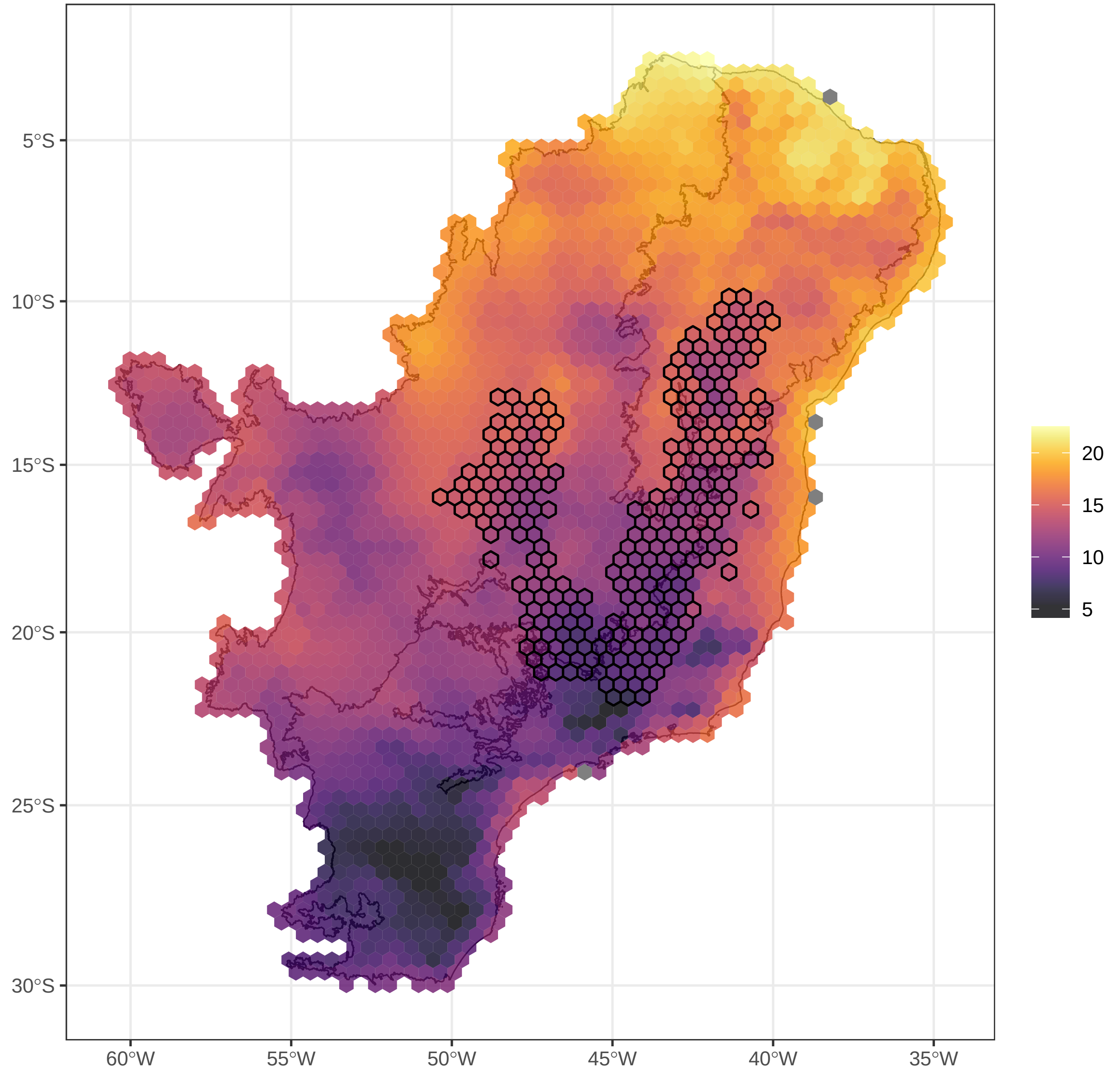


**Figure S3.6.**Minimum temperature of the coldest month values assigned for each cell. Cells assigned to *campos rupestres* are outlined.

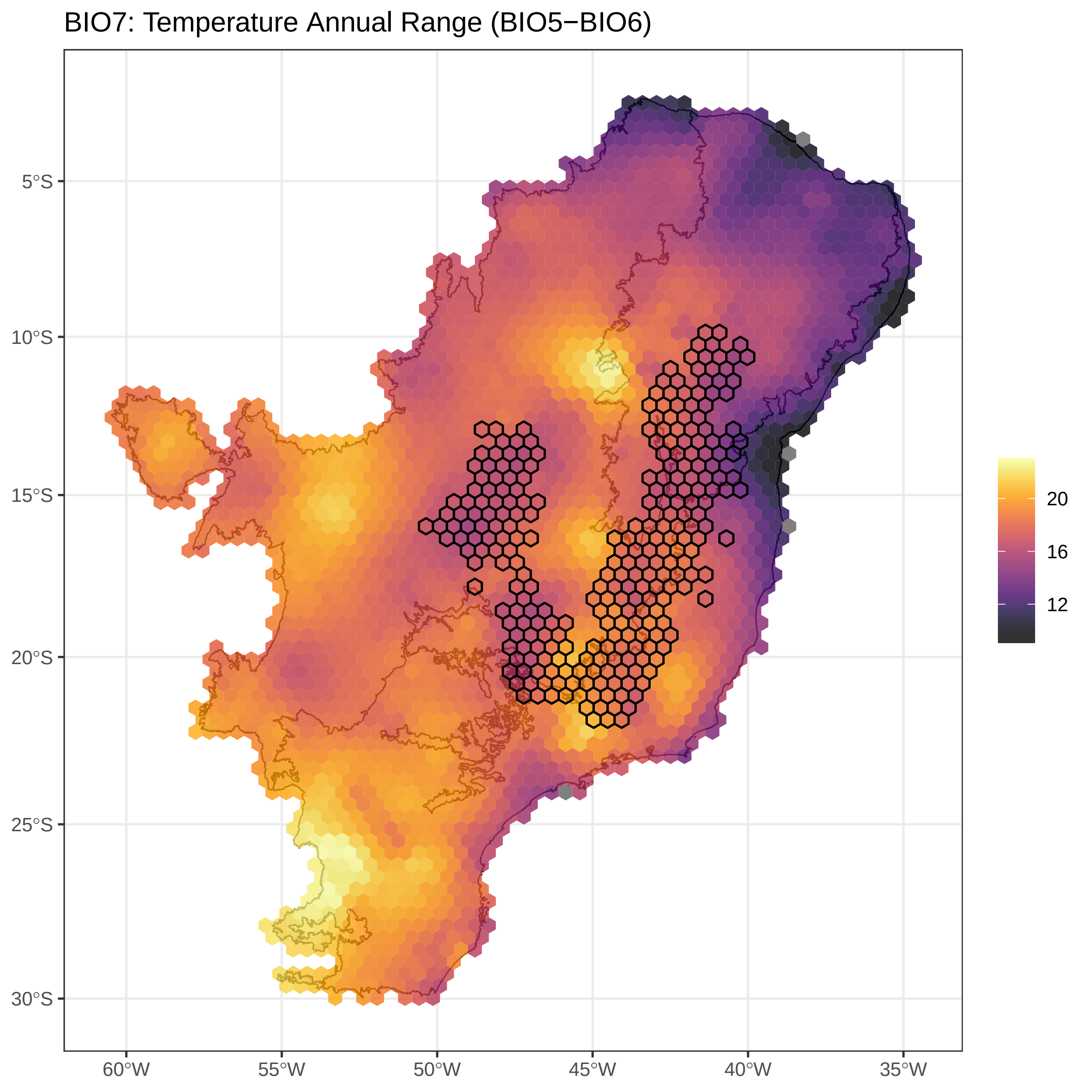


**Figure S3.7.**Temperature annual range values assigned for each cell. Cells assigned to *campos rupestres* are outlined.

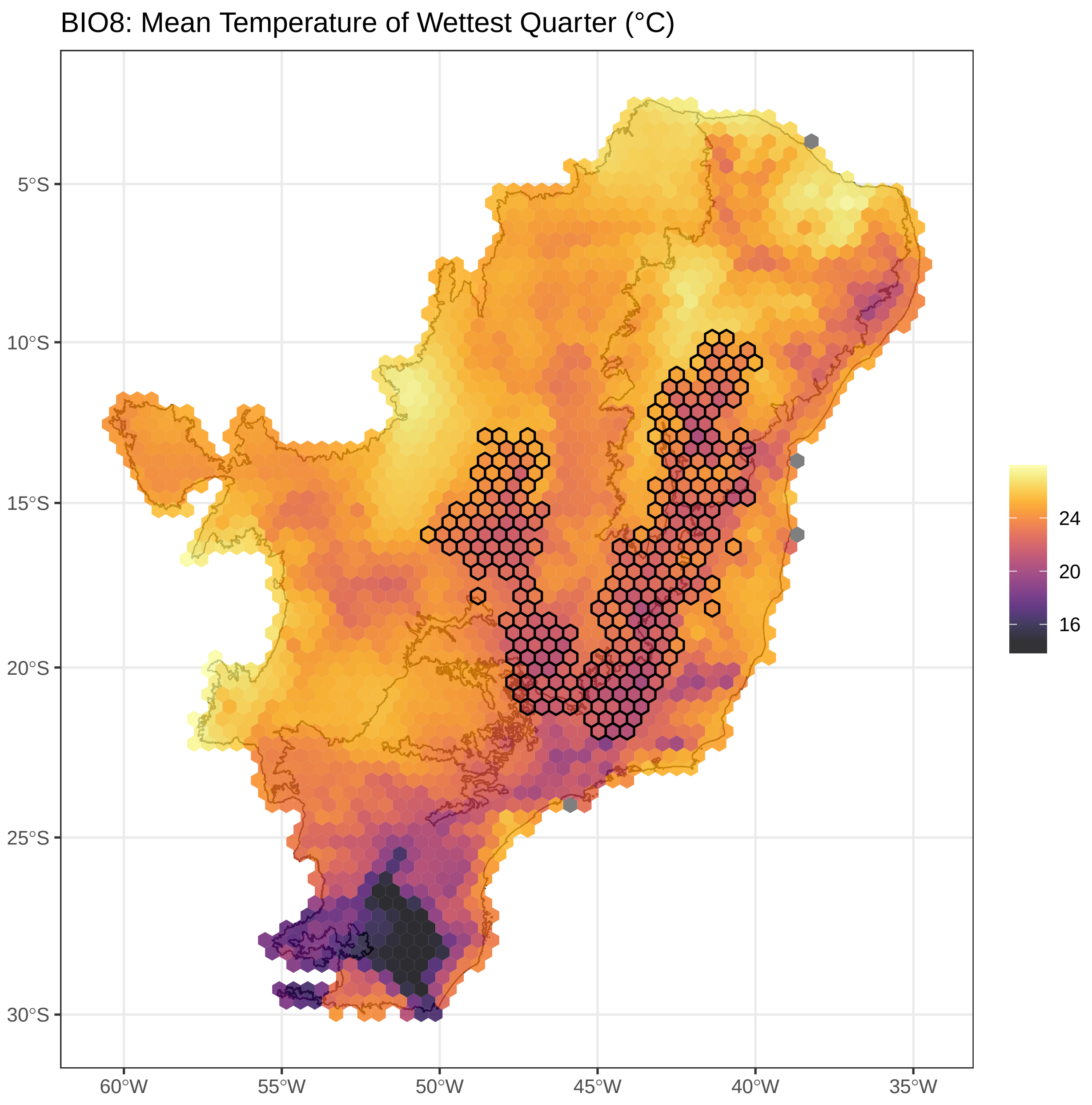


**Figure S3.8.**Mean temperature of the wettest quarter values assigned for each cell. Cells assigned to *campos rupestres* are outlined.

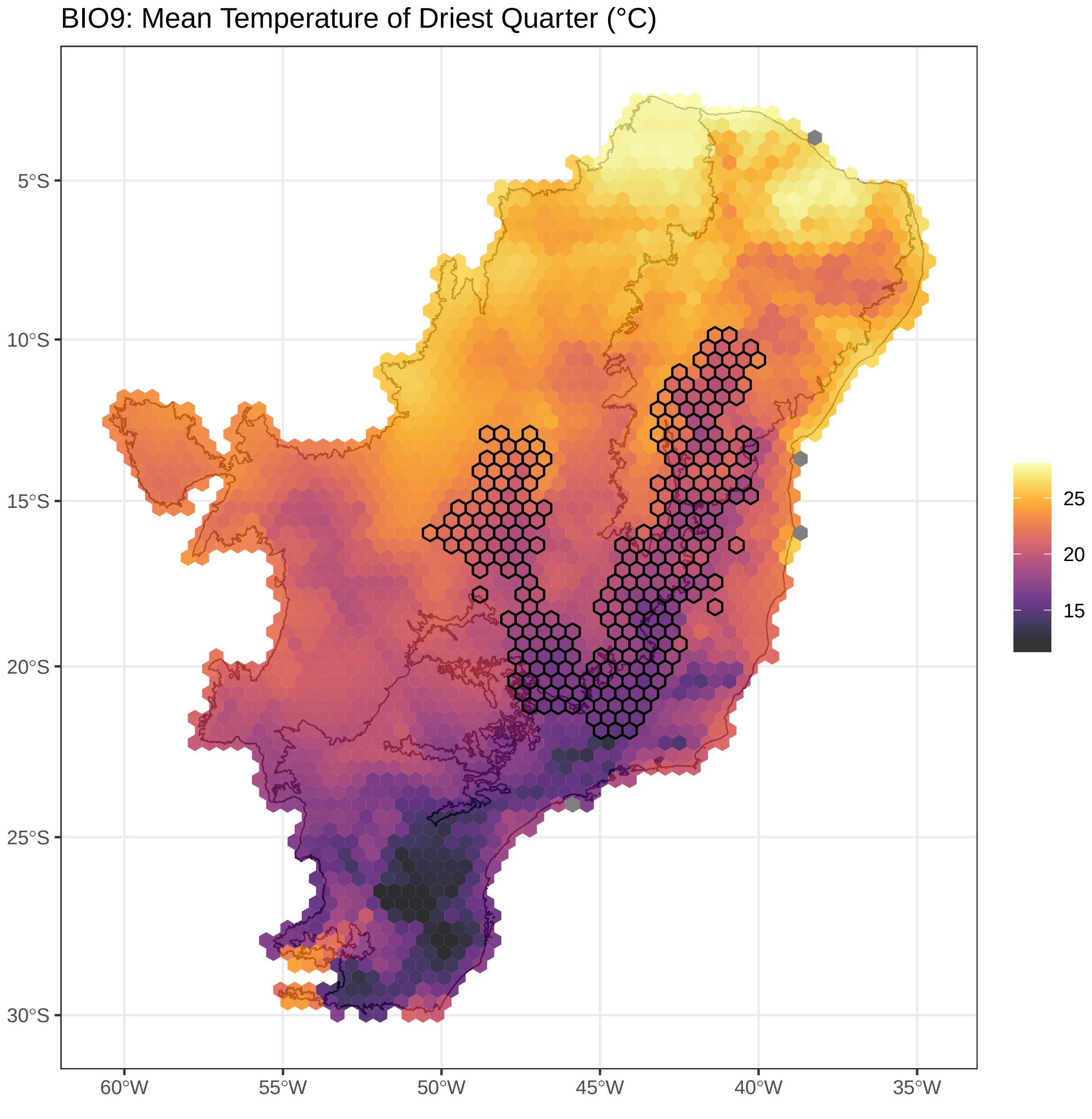


**Figure S3.9.** Mean temperature of the driest quarter values assigned for each cell. Cells assigned to *campos rupestres* are outlined.

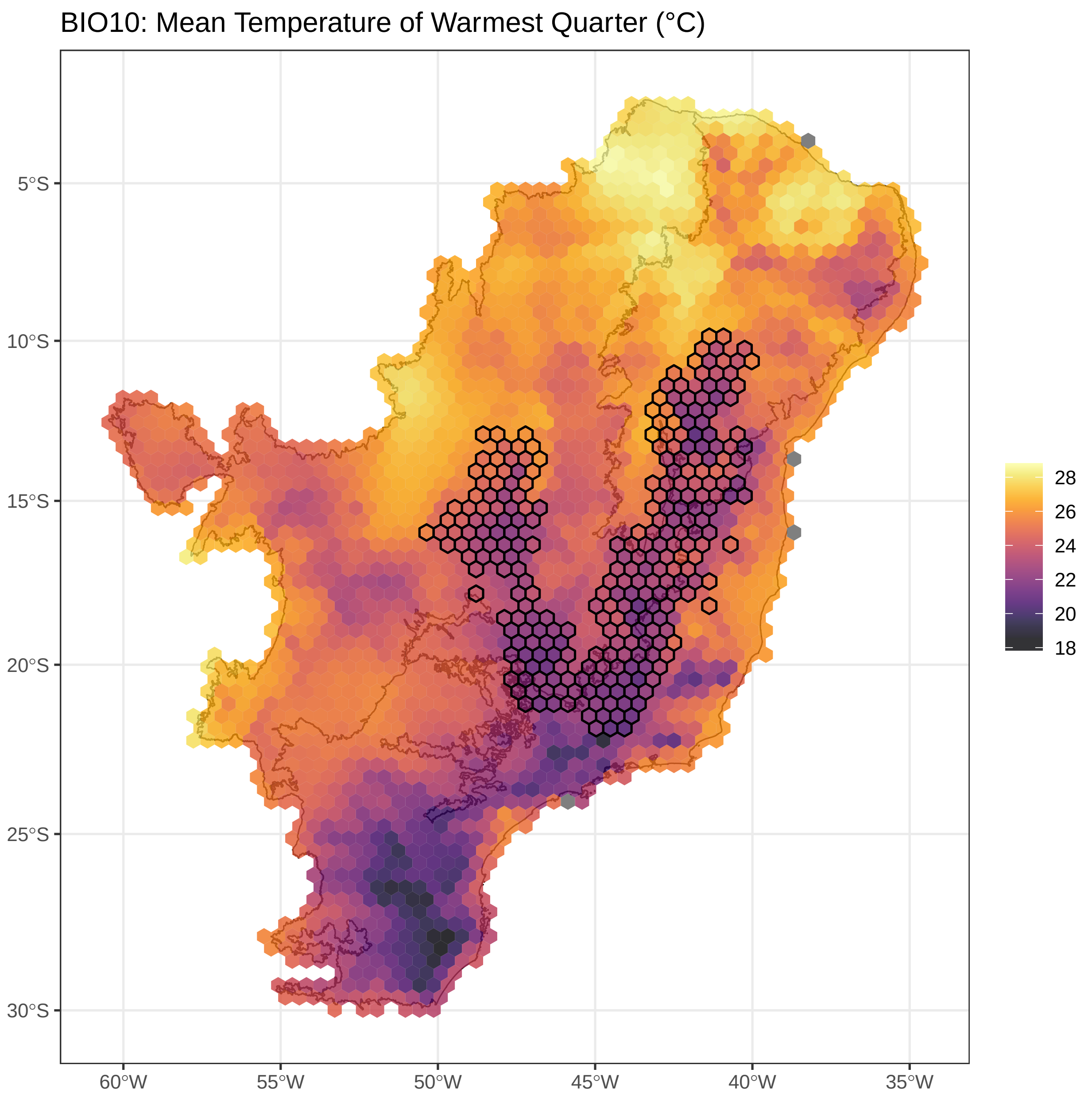


**Figure S3.10.** Mean temperature of the warmest quarter values assigned for each cell. Cells assigned to *campos rupestres* are outlined.

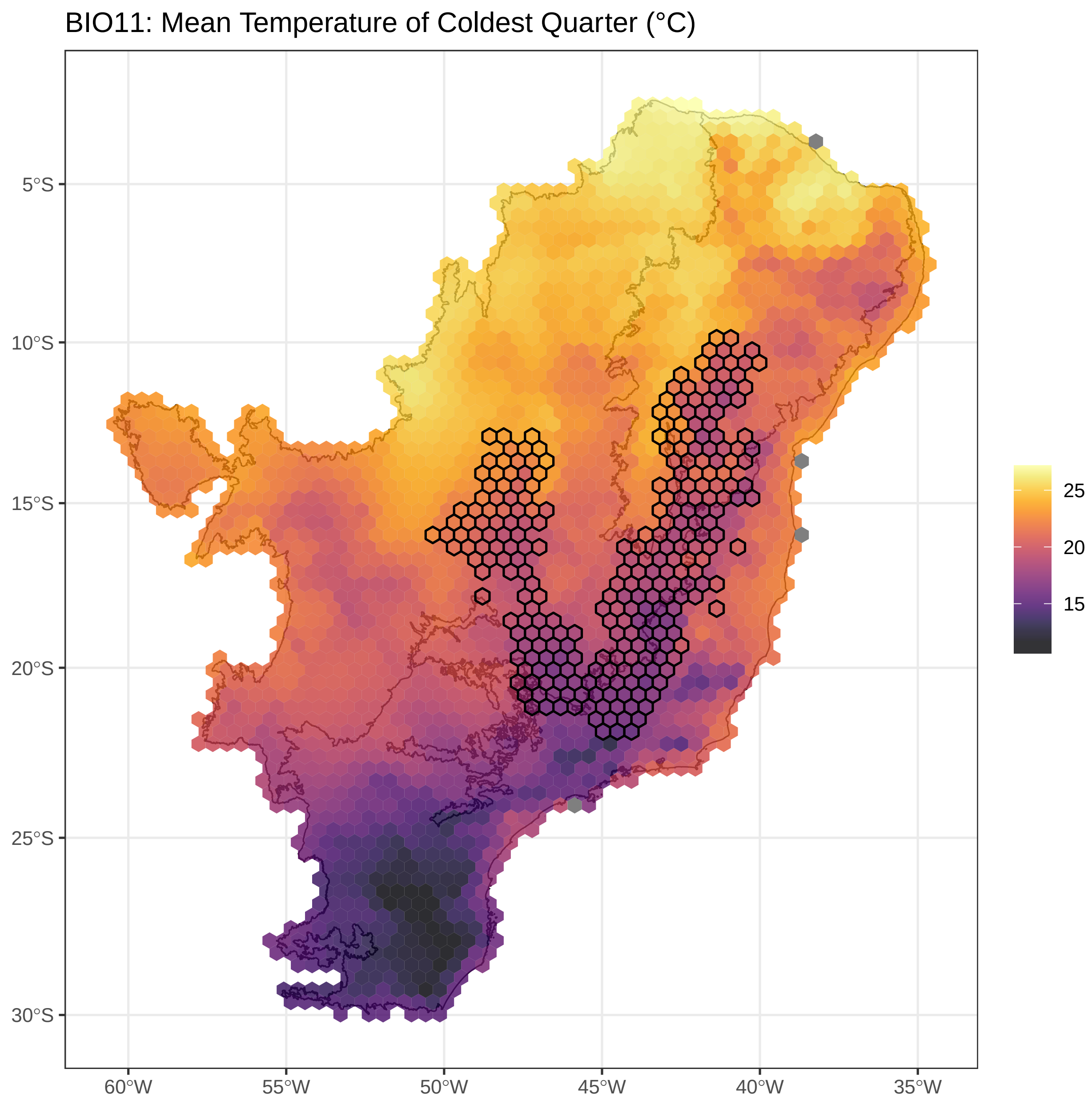


**Figure S3.11.**Mean temperature of the coldest quarter values assigned for each cell. Cells assigned to *campos rupestres* are outlined.

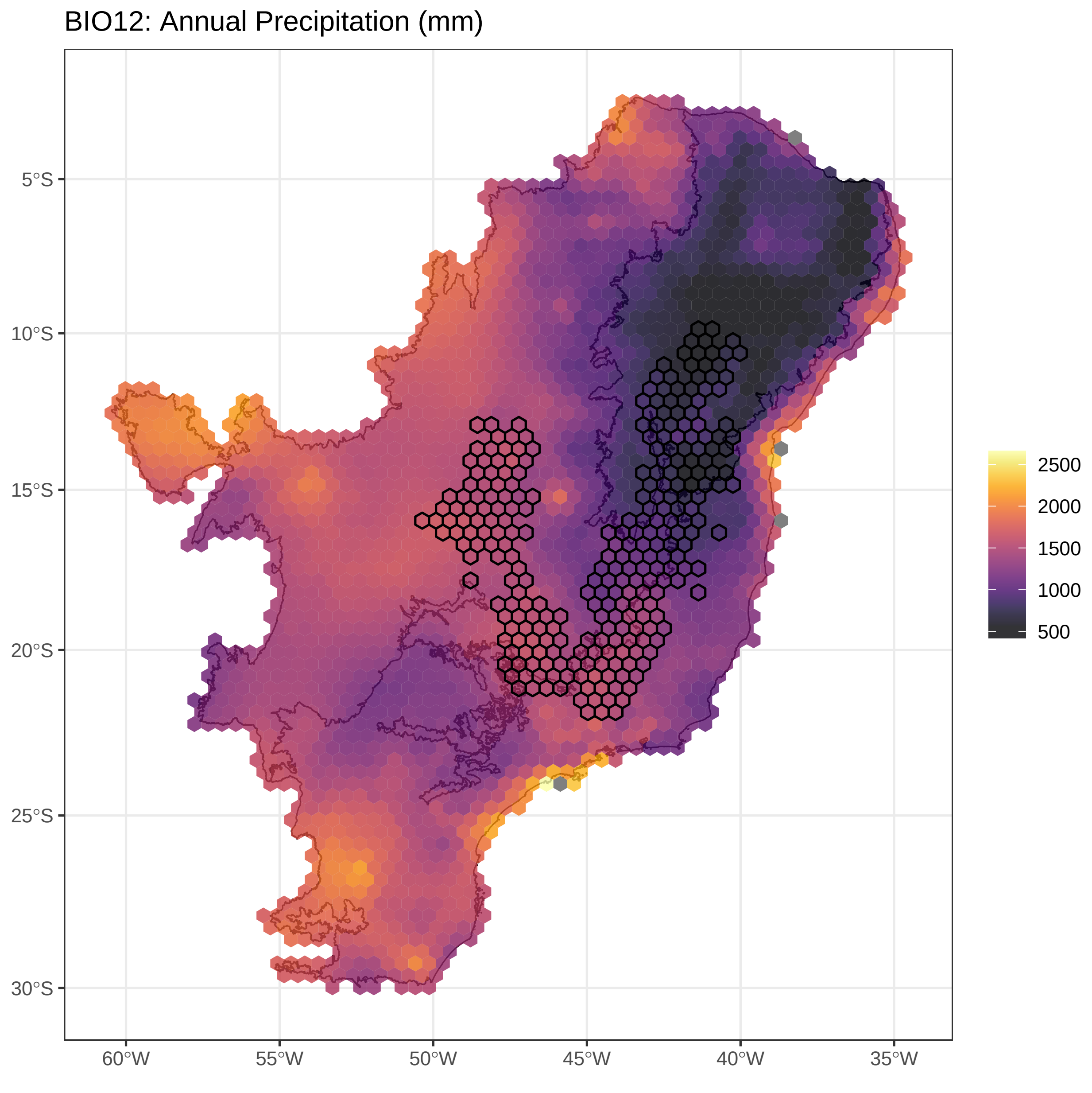


**Figure S3.12.**Annual precipitation values assigned for each cell. Cells assigned to *campos rupestres* are outlined.

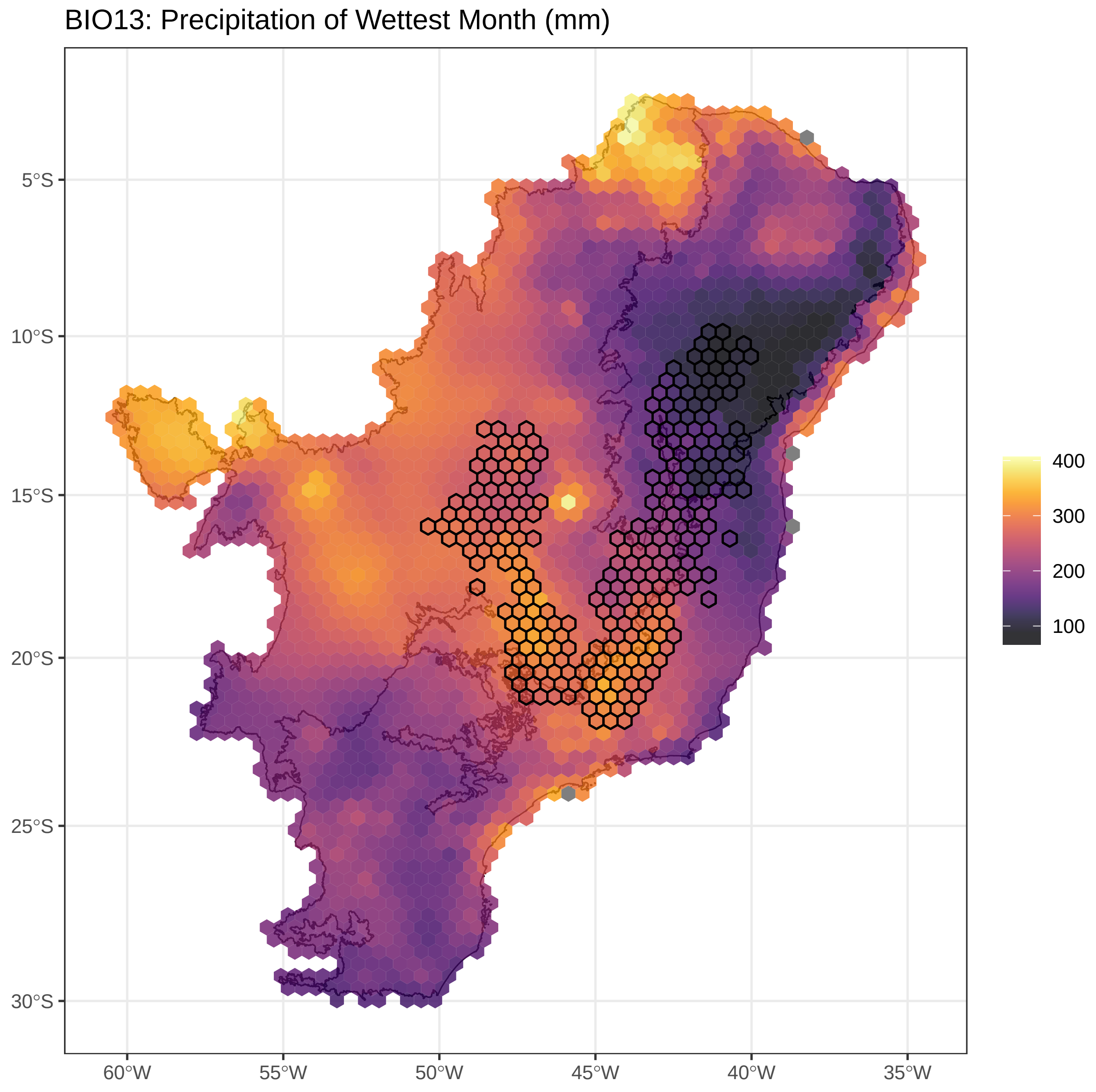


**Figure S3.13.**Precipitation of the wettest month values assigned for each cell. Cells assigned to *campos rupestres* are outlined.

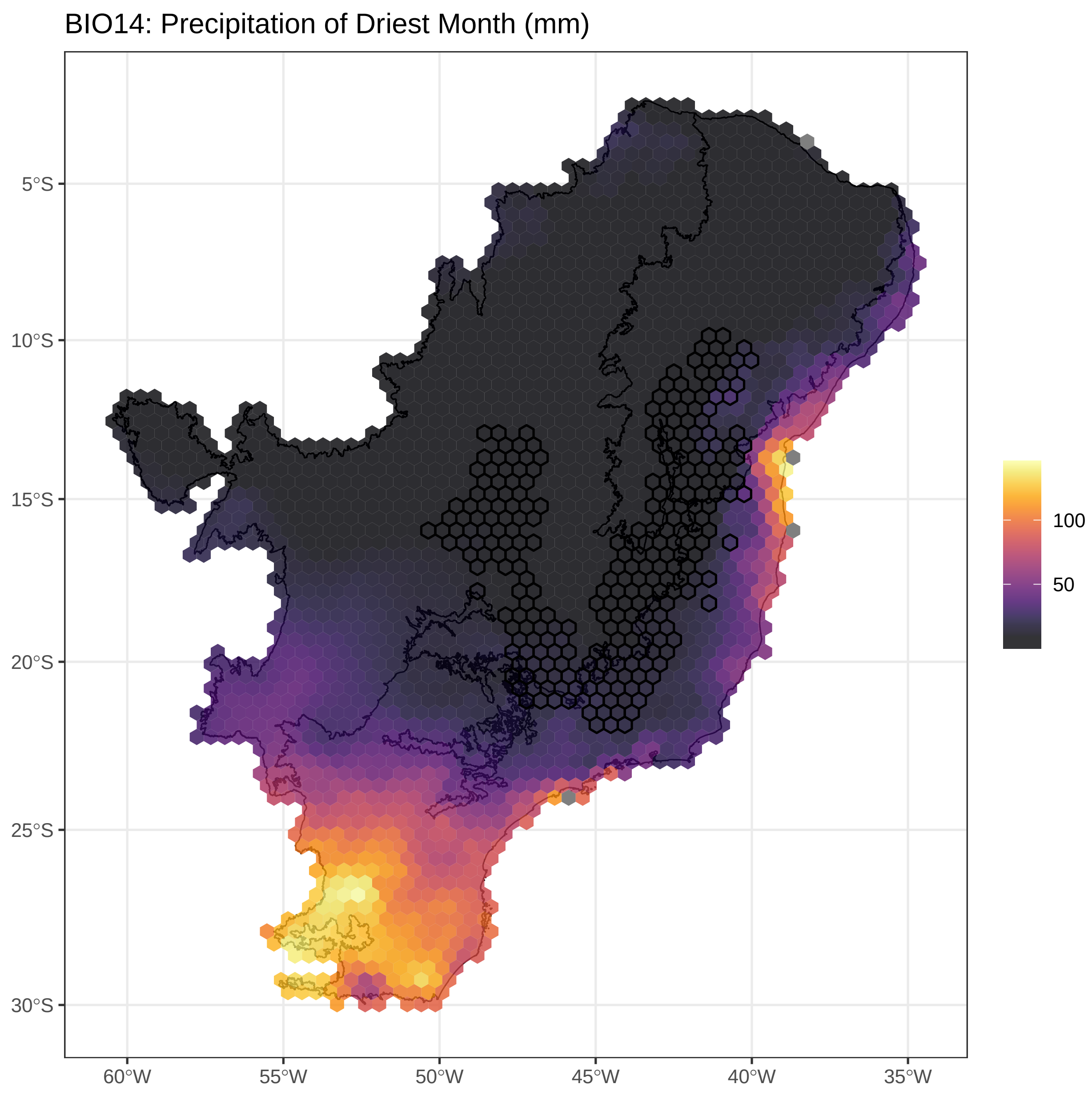


**Figure S3.14.**Precipitation of the driest month values assigned for each cell. Cells assigned to *campos rupestres* are outlined.

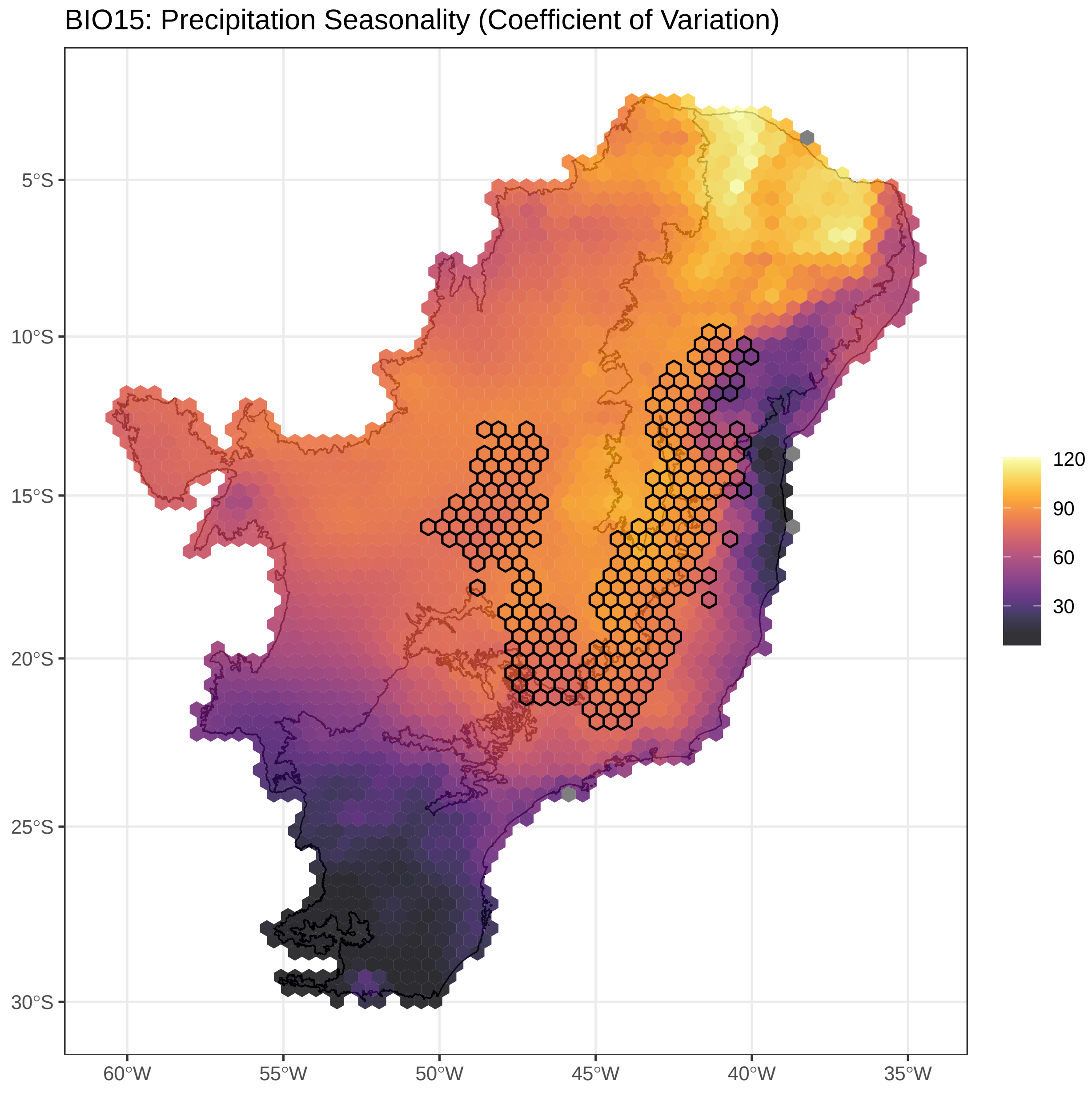


**Figure S3.15.**Precipitation seasonality values assigned for each cell. Cells assigned to *campos rupestres* are outlined.

BIO16: Precipitation of Wettest Quarter (mm)

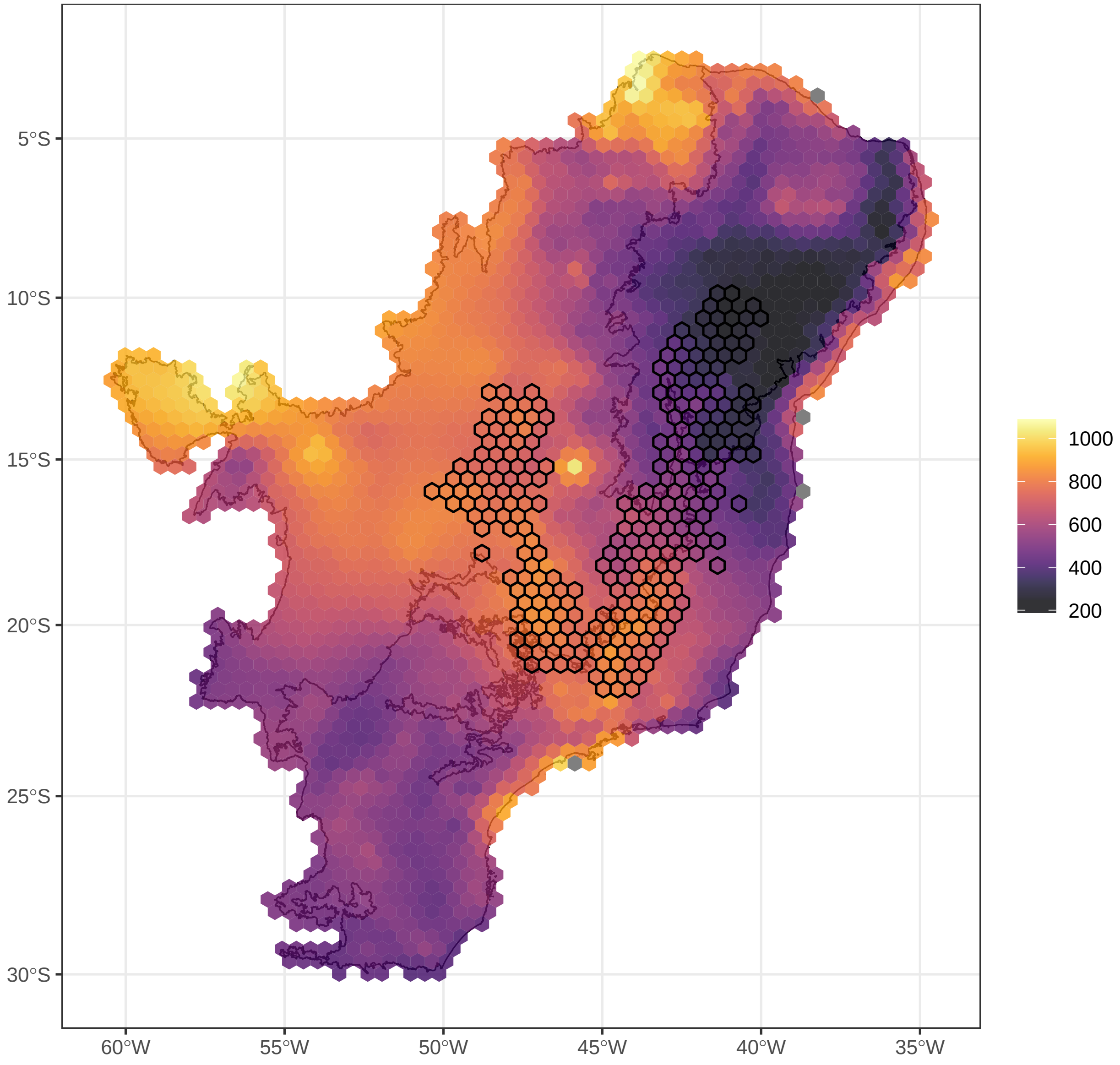


**Figure S3.16.**Precipitation of the wettest quarter values assigned for each cell. Cells assigned to *campos rupestres* are outlined.

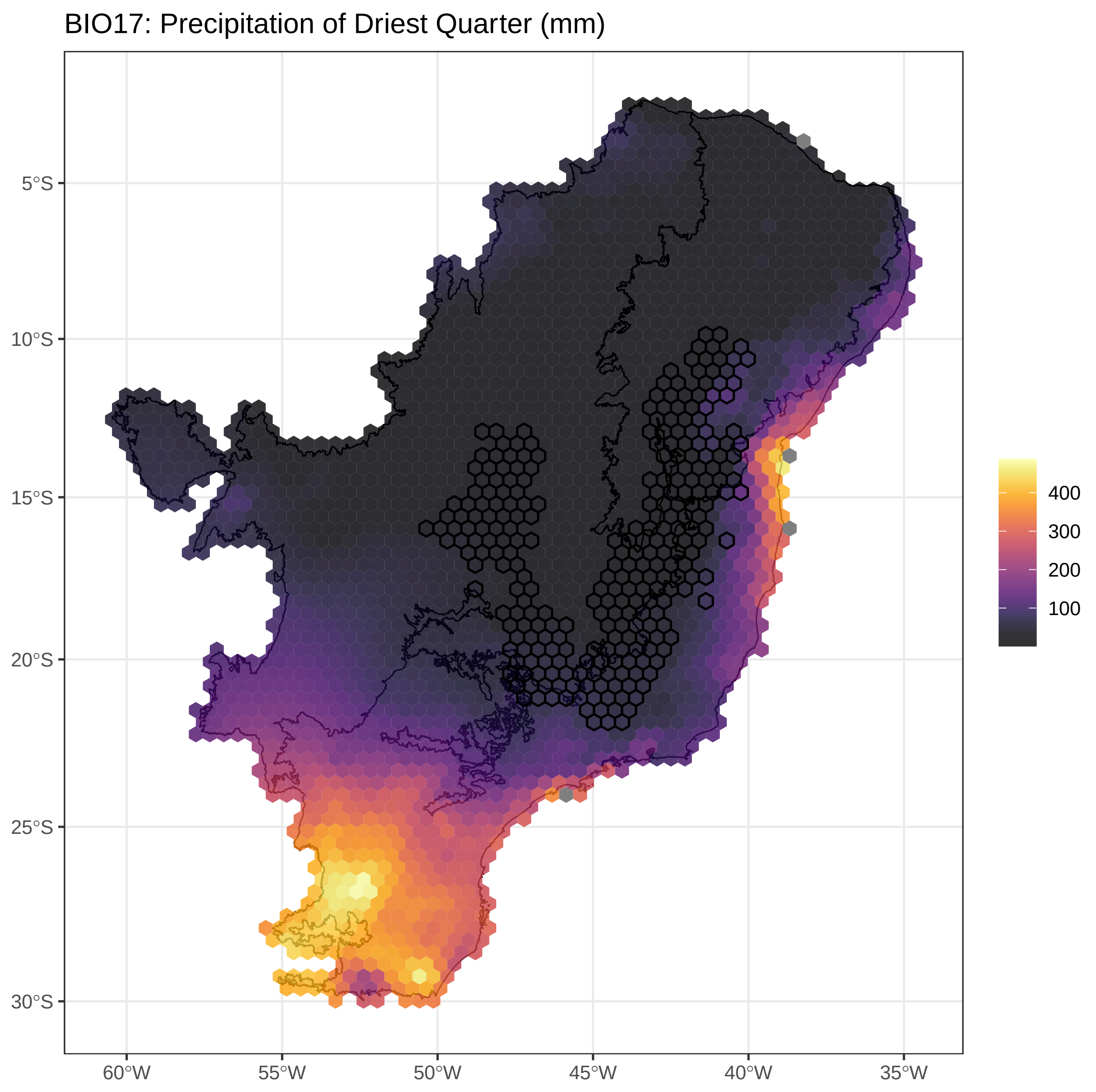


**Figure S3.17.**Precipitation of the driest quarter values assigned for each cell. Cells assigned to *campos rupestres* are outlined.

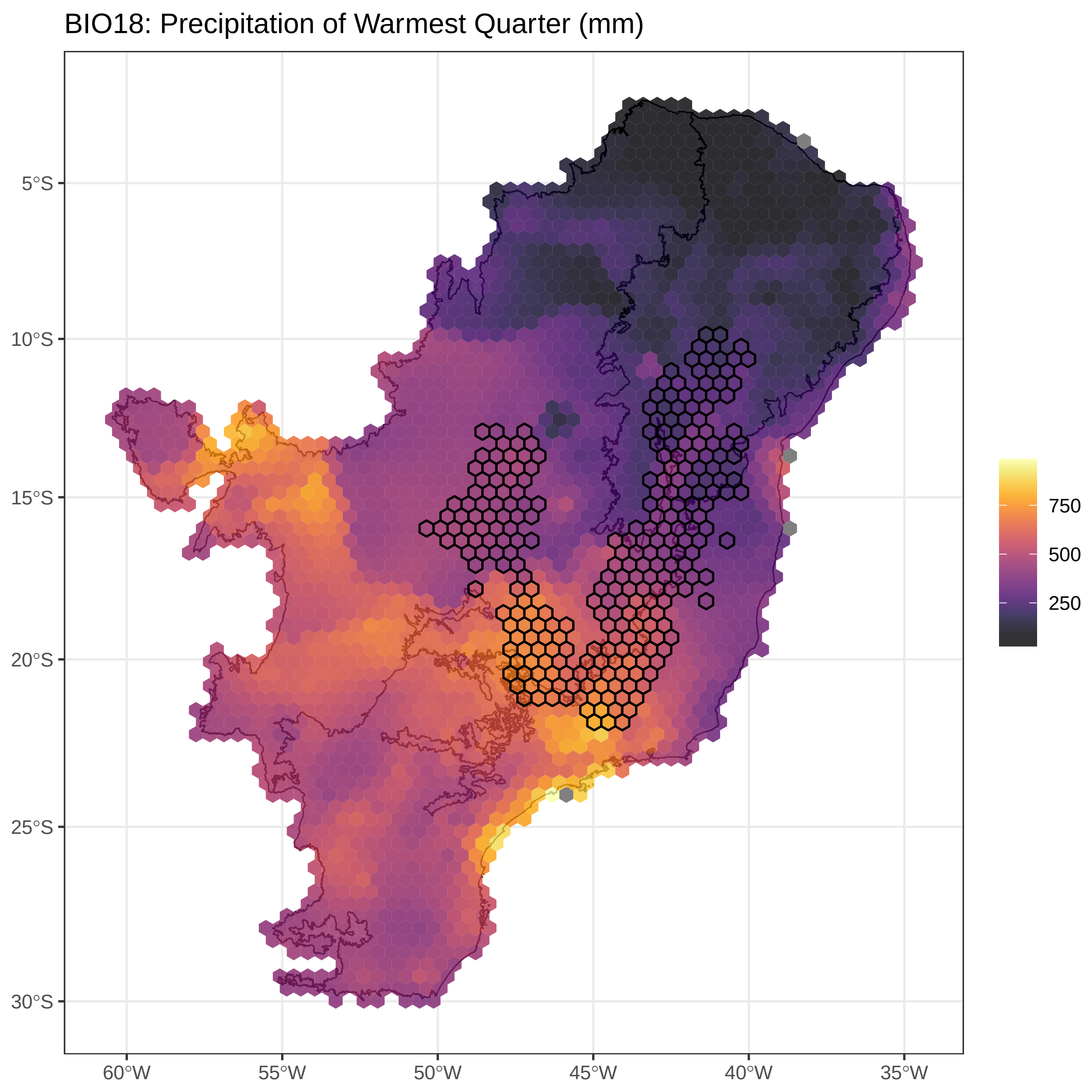


**Figure S3.18.**Precipitation of the warmest quarter values assigned for each cell. Cells assigned to *campos rupestres* are outlined.

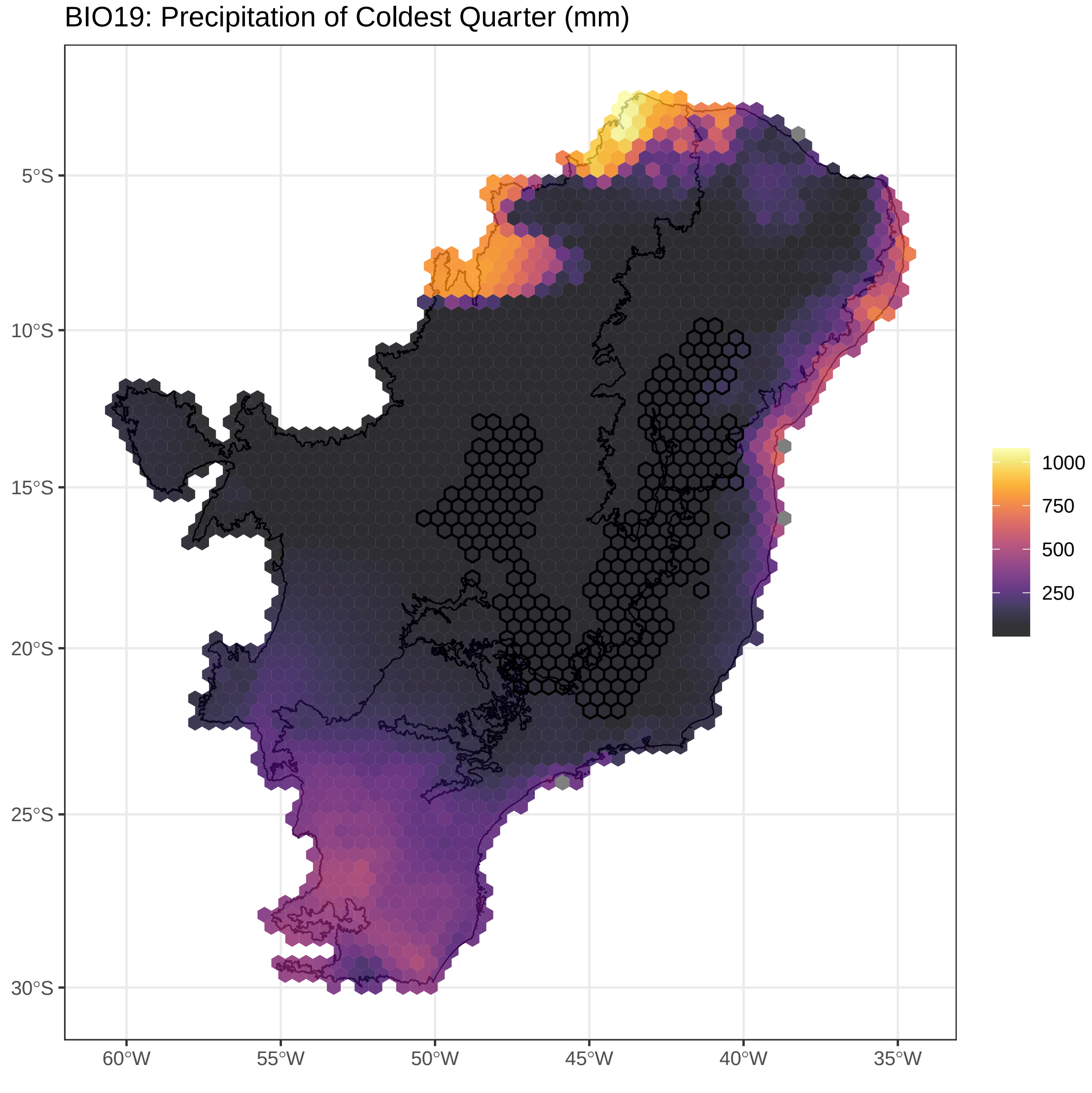


**Figure S3.19.**Precipitation of the coldest quarter values assigned for each cell. Cells assigned to *campos rupestres* are outlined.

#### Edaphic variables

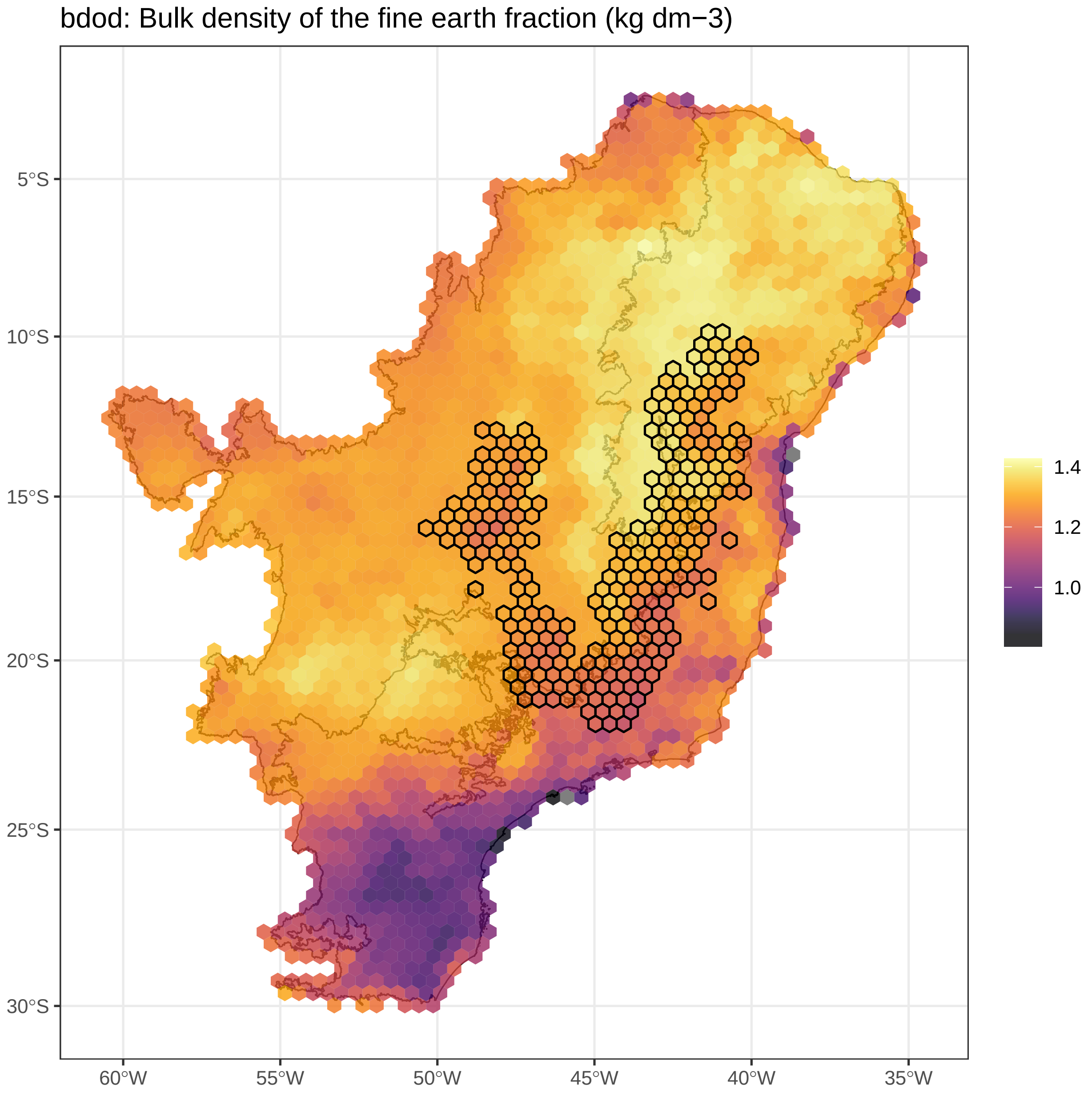


**Figure S3.20.**Bulk density values assigned for each cell. Cells assigned to *campos rupestres* are outlined.

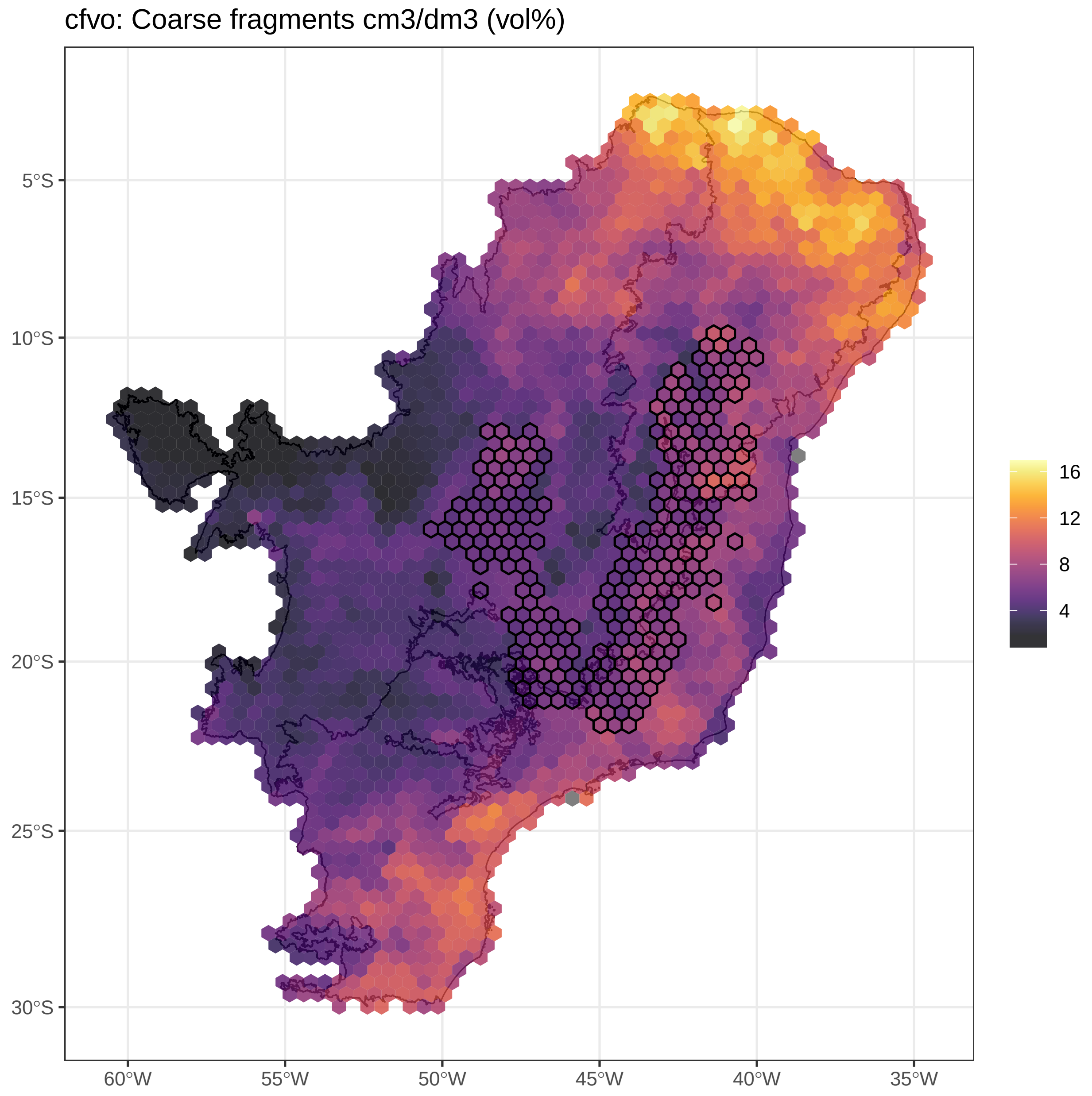


**Figure S3.21.**Coarse fragments values assigned for each cell. Cells assigned to *campos rupestres* are outlined.

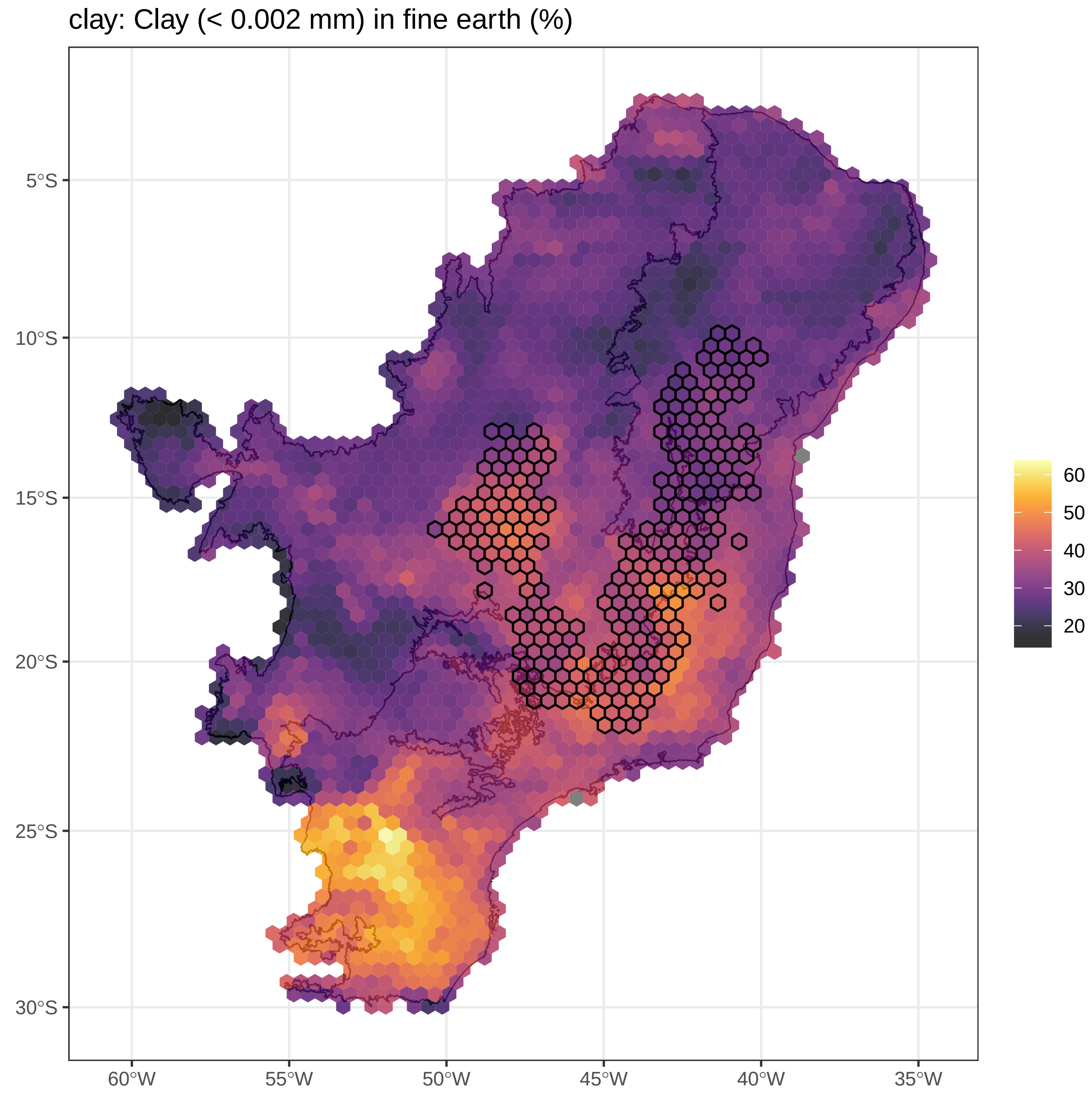


**Figure S3.22.** Clay values assigned for each cell. Cells assigned to *campos rupestres* are outlined.

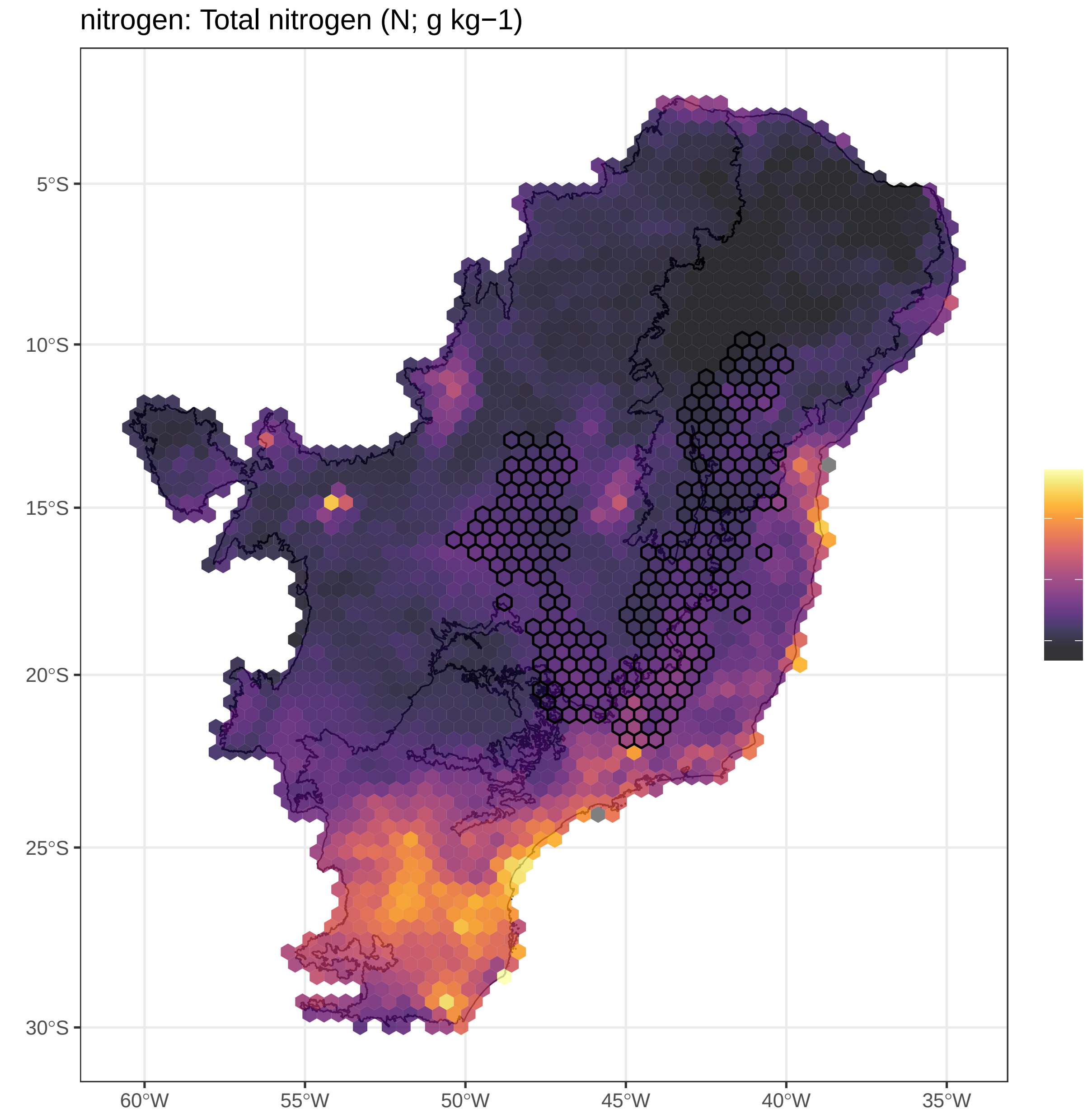


**Figure S3.23.**Total nitrogen values assigned for each cell. Cells assigned to *campos rupestres* are outlined.

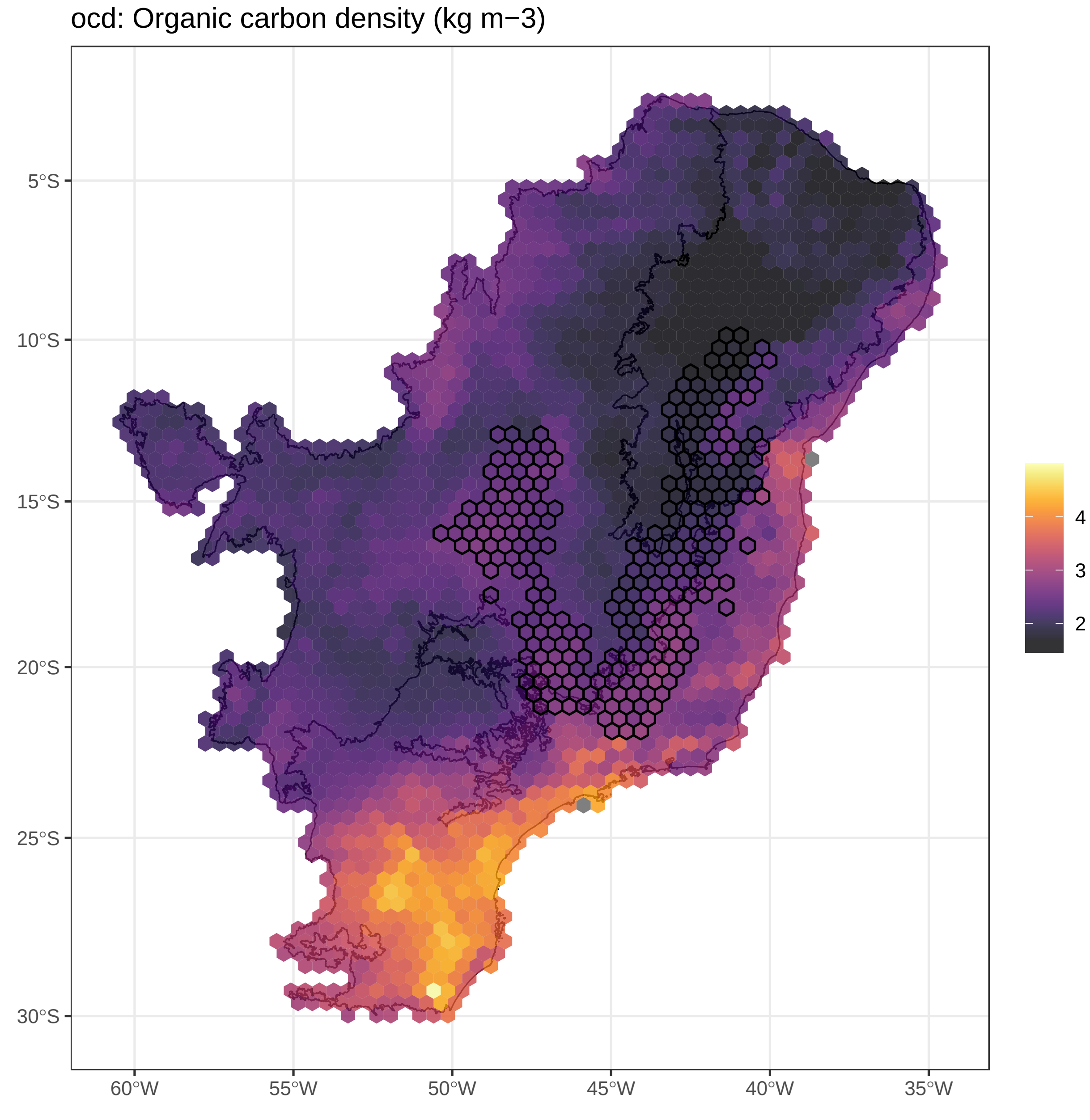


**Figure S3.24.**Organic carbon density values assigned for each cell. Cells assigned to *campos rupestres* are outlined.

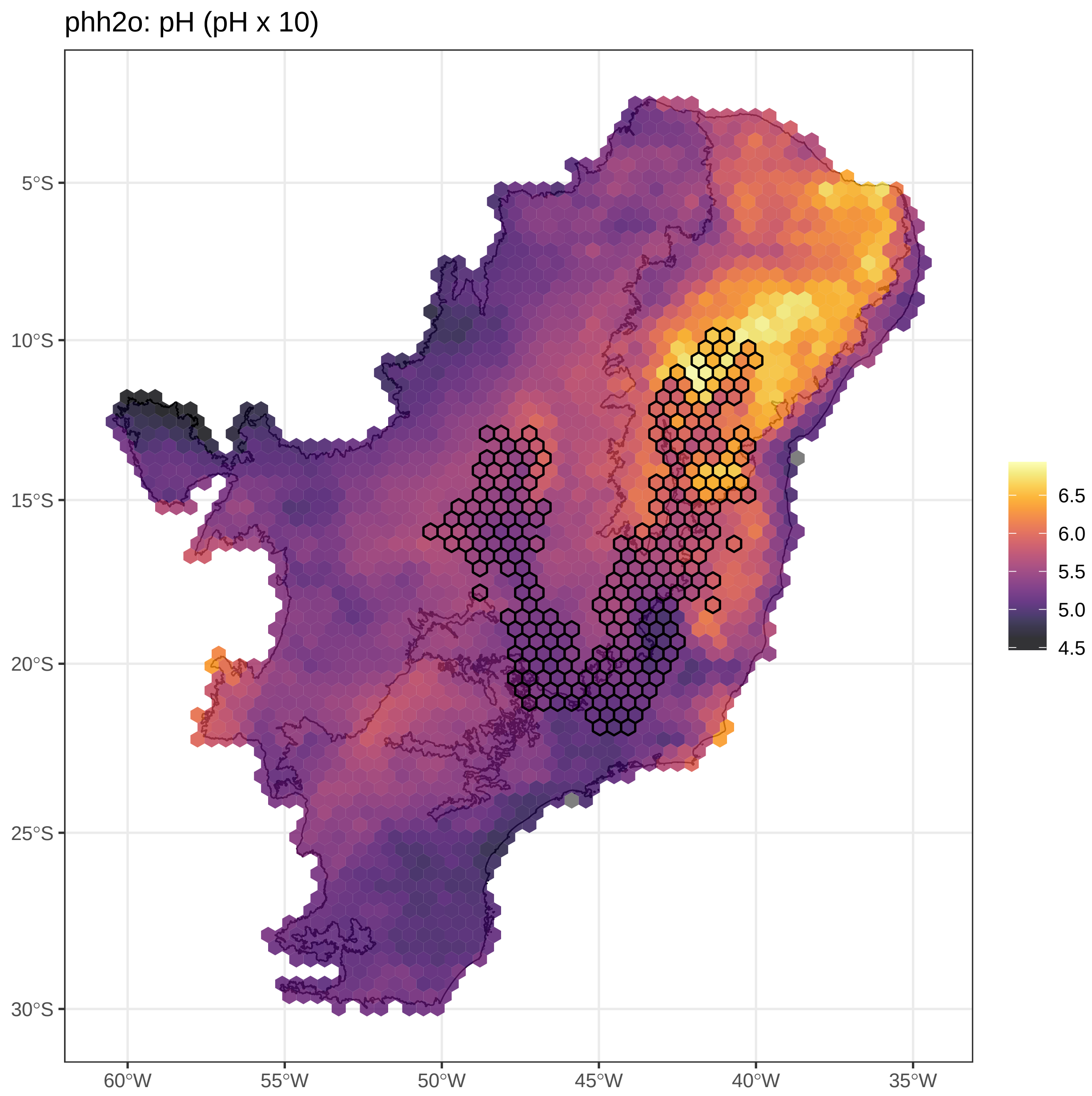


**Figure S3.25.**pH in H2O values assigned for each cell. Cells assigned to *campos rupestres* are outlined.

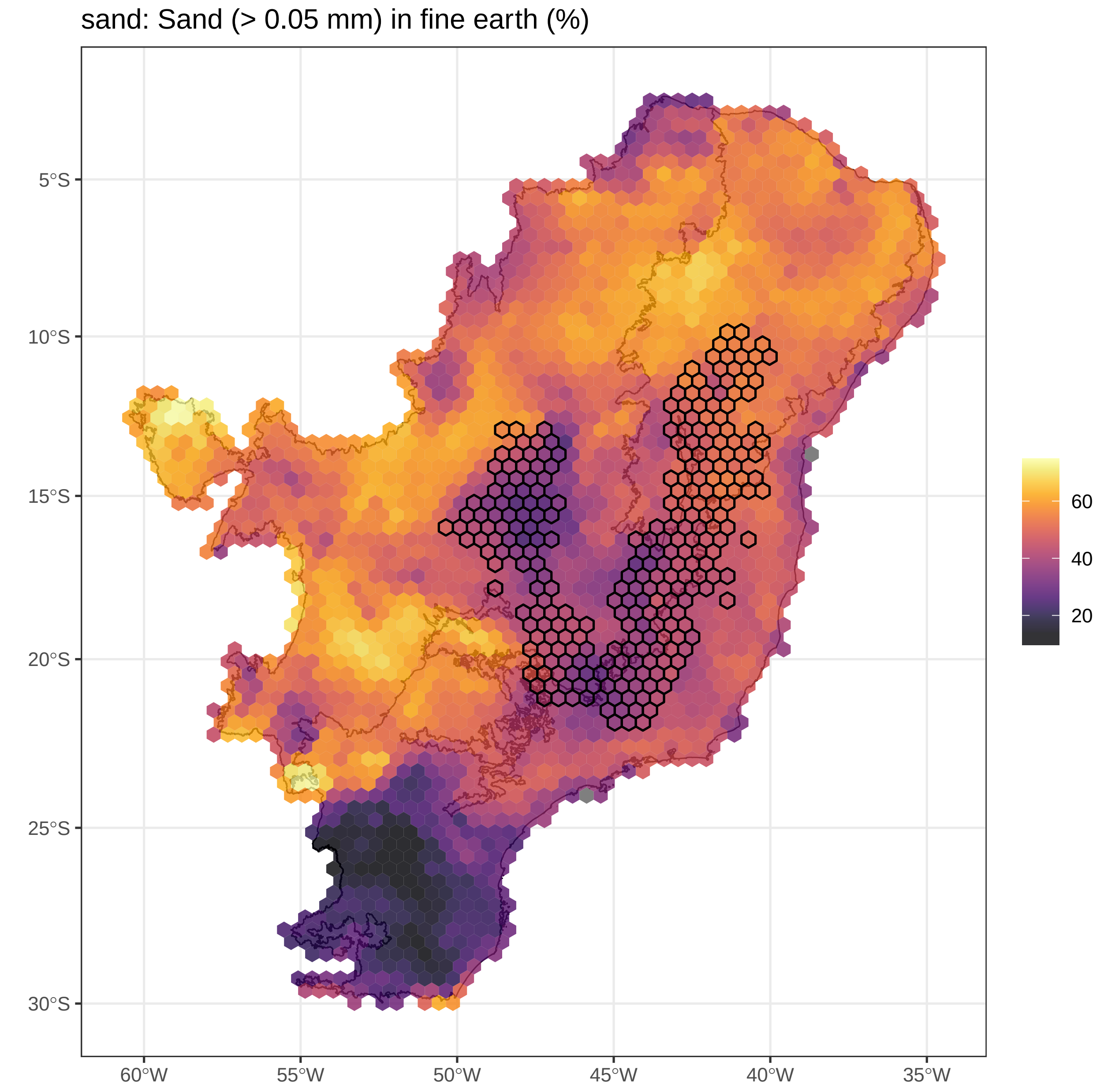


**Figure S3.26.**Sand values assigned for each cell. Cells assigned to *campos rupestres* are outlined.

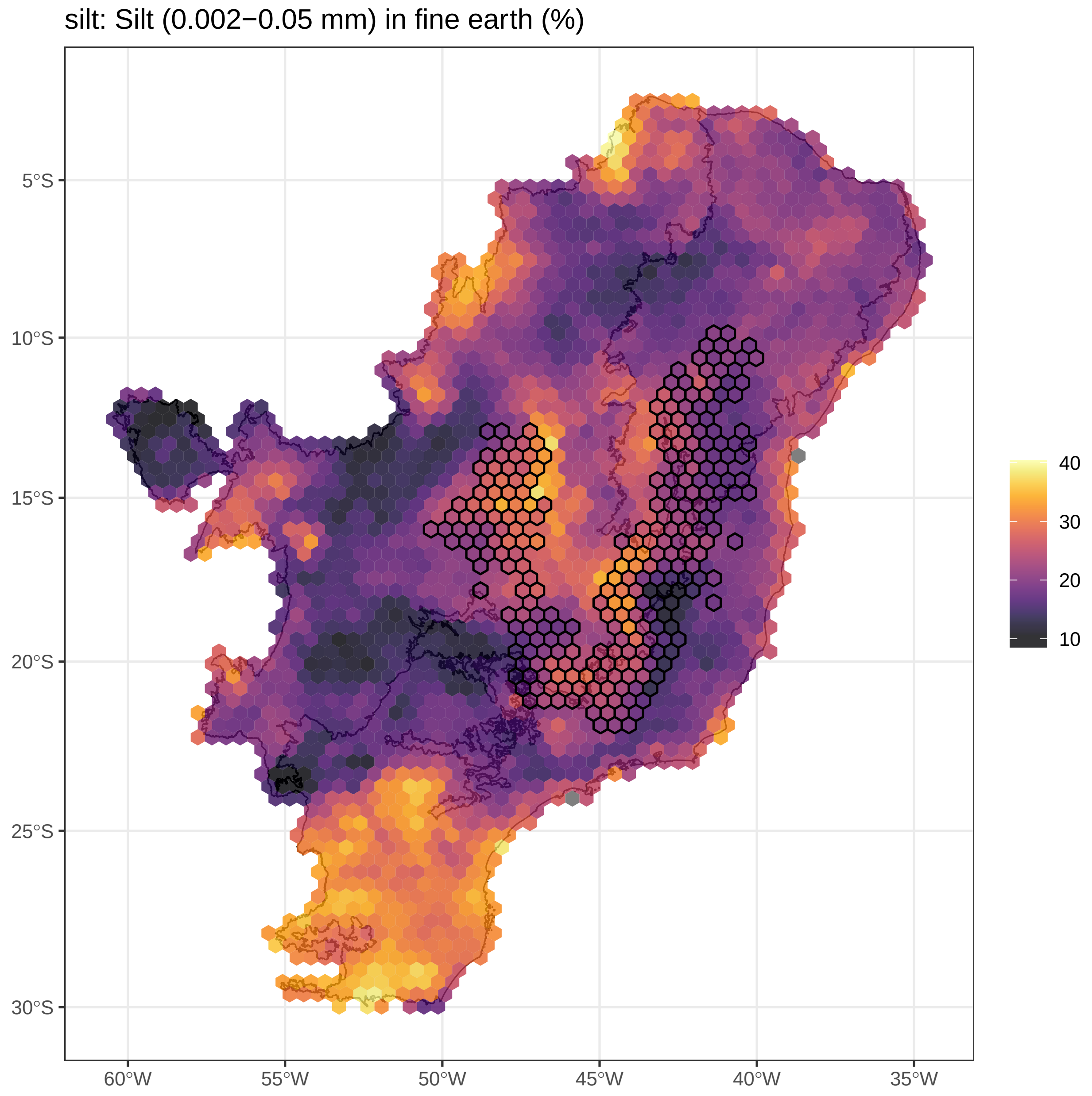


**Figure S3.27.**Silt values assigned for each cell. Cells assigned to *campos rupestres* are outlined.

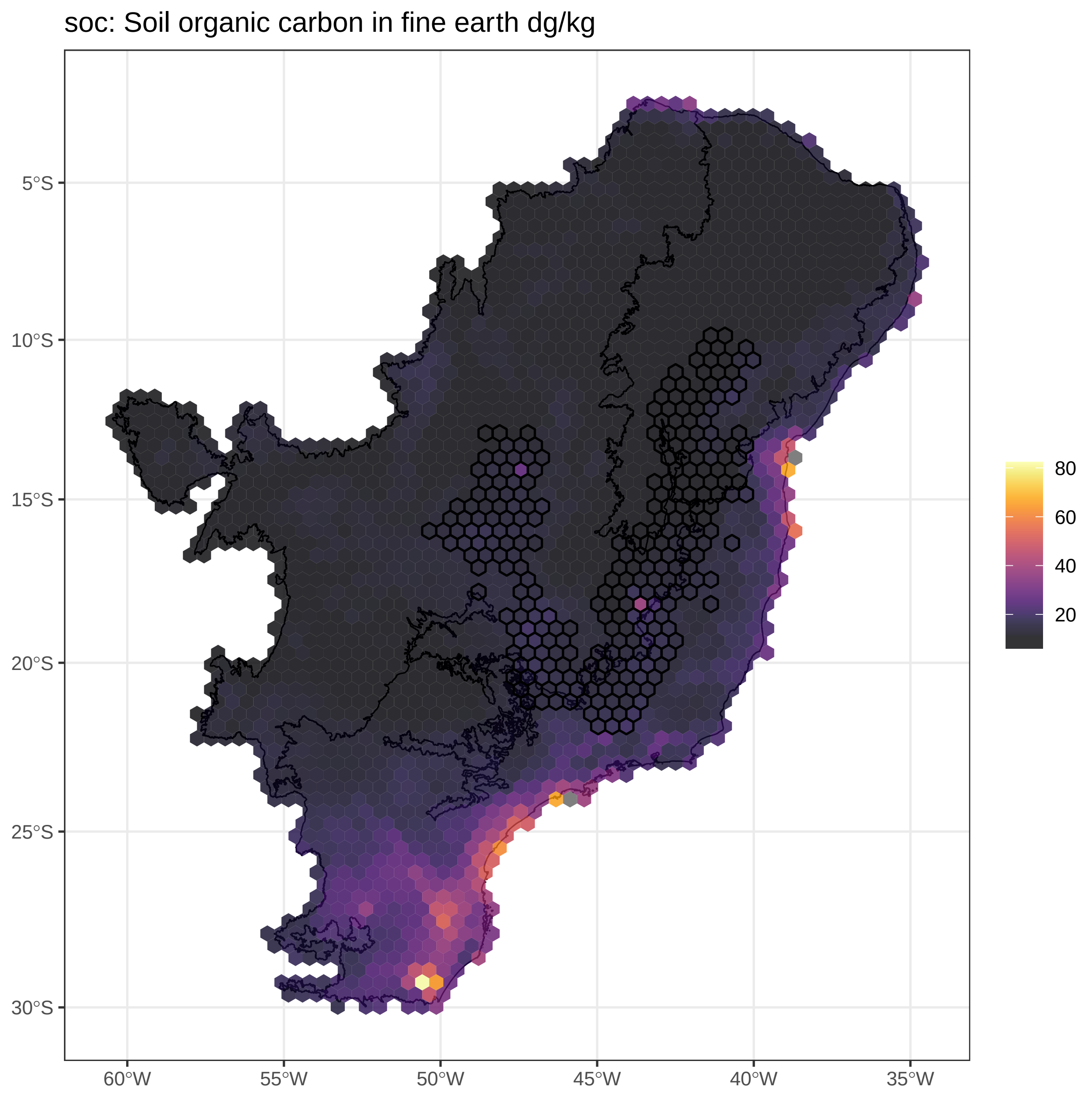


**Figure S3.28.**Soil organic carbon values assigned for each cell. Cells assigned to *campos rupestres* are outlined.

#### Correlation matrix

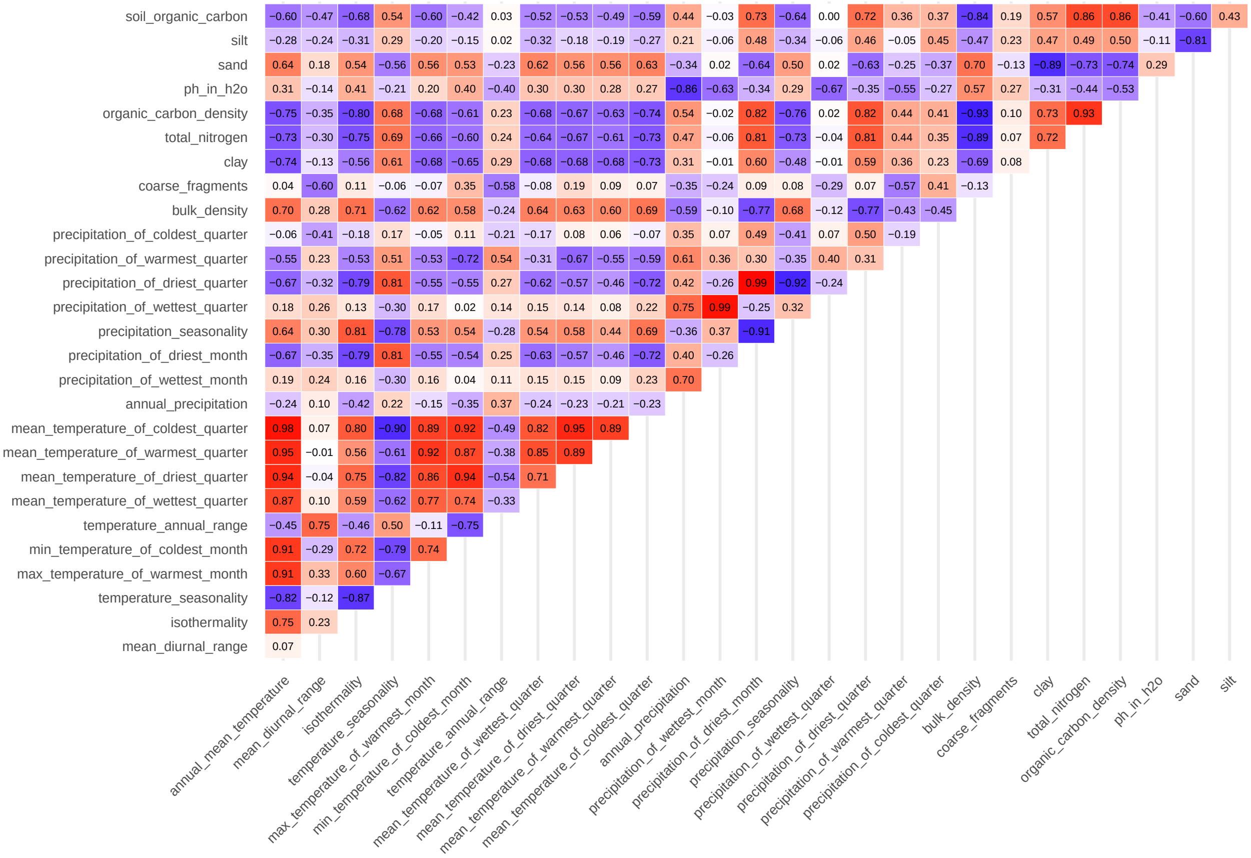


**Figure S3.29.**Heatmap showing pairwise correlations among abiotic variables, calculated using Pearson's correlation coefficient. To minimize multicollinearity between variables assigned to grid cells, we examined pairwise correlations among variables within each category (precipitation, temperature, and soil) and retained only uncorrelated subsets ($|r| \leq 0.75$). This yielded a final set of 15 predictors: (1) annual precipitation, (2) precipitation of the wettest, (3) driest, (4) warmest, and (5) coldest quarters, (6) annual mean temperature, (7) mean diurnal range, (8) isothermality, (9) temperature annual range, (10) coarse fragments, (11) clay, (12) sand, and (13) silt contents, (14) total nitrogen, and (15) soil pH. Although clay, sand, and silt contents are inherently correlated, we retained them because they represent proportions of distinct soil textures. Cell colors and values indicate the strength and direction of correlations, ranging from $-1$ (blue; negative) to 1 (red; positive).

### Appendix 4: GDM outputs

| | *Calliandra* | *Cattleya* | *Chamaecrista* | *Diplusodon* | *Dyckia* | *Habenaria* | Lychnophorinae | Marcetieae | *Mimosa* | *Paepalanthus* | *Myrcia* | Trimezieae | Velloziaceae |
|---|---|---|---|---|---|---|---|---|---|---|---|---|---|
| Deviance explained (%) | 20.110 | 39.630 | 7.460 | 22.380 | 28.280 | 8.150 | 17.970 | 24.860 | 19.530 | 14.870 | 6.580 | 21.610 | 17.850 |
| Intercept | 0.350 | 0.370 | 0.440 | 0.360 | 0.370 | 0.840 | 0.300 | 0.340 | 0.640 | 0.390 | 0.610 | 0.260 | 0.300 |
| Geographic | **0.390** | **0.990** | 0.050 | **0.290** | **0.460** | 0.080 | **0.260** | **1.200** | **0.260** | **0.660** | 0.050 | 0.110 | **0.220** |
| Annual Precipitation | **0.200** | - | **0.150** | **0.410** | 0.020 | - | - | - | **0.250** | **0.170** | 0.100 | - | 0.020 |
| Precipitation of Wettest Quarter | 0.160 | **0.250** | 0.110 | **0.560** | - | 0.270 | 0.160 | 0.240 | 0.130 | 0.140 | 0.050 | **0.540** | 0.130 |
| Precipitation of Driest Quarter | 0.110 | **0.290** | **0.170** | 0.220 | 0.050 | **0.410** | - | **0.350** | **0.180** | - | **0.370** | **0.580** | 0.070 |
| Precipitation of Warmest Quarter | 0.110 | - | 0.050 | - | - | - | - | 0.030 | 0.050 | **0.220** | - | - | 0.060 |
| Precipitation of Coldest Quarter | - | 0.260 | 0.110 | 0.120 | 0.190 | 0.120 | 0.100 | - | 0.050 | - | 0.030 | 0.130 | 0.020 |
| Annual Mean Temperature | **0.150** | **0.810** | 0.070 | 0.140 | 0.190 | **0.520** | - | **0.540** | **0.440** | 0.020 | 0.100 | - | **0.440** |
| Mean Diurnal Range | - | - | - | - | 0.060 | **0.190** | 0.050 | 0.070 | 0.140 | 0.000 | 0.100 | **0.190** | 0.140 |
| Isothermality | 0.060 | - | 0.090 | - | **0.460** | - | 0.010 | - | 0.100 | 0.000 | 0.060 | **0.360** | **0.260** |
| Temperature Annual Range | **0.220** | 0.060 | 0.080 | - | 0.230 | - | **0.140** | 0.170 | - | 0.000 | **0.100** | - | **0.240** |
| Coarse Fragments | - | **0.760** | 0.100 | 0.050 | **1.070** | **0.660** | **0.160** | **0.480** | 0.010 | **0.210** | **0.150** | **0.760** | - |
| Clay | 0.060 | 0.050 | - | - | 0.390 | 0.040 | 0.020 | - | 0.040 | - | 0.040 | - | - |
| Total Nitrogen | 0.020 | 0.140 | 0.040 | - | 0.040 | **0.190** | - | - | 0.030 | - | 0.020 | - | - |
| pH in H2O | - | 0.170 | - | - | 0.020 | - | - | 0.180 | 0.020 | - | 0.020 | - | **0.120** |
| Sand | - | - | 0.010 | - | - | - | - | - | - | 0.050 | - | - | - |
| Silt | **0.320** | 0.180 | **0.220** | 0.180 | - | **0.230** | 0.050 | 0.130 | 0.080 | **0.300** | **0.120** | - | 0.020 |

**Table S4.1.**GDM results for each plant group. First row shows the percentage of phylogenetic dissimilarity explained by the predictors (Deviance explained). Second row shows the mean dissimilarity when site-pairs have identical predictor values (Intercept). Remaining rows show the sum of coefficients of each predictor. The higher the sum of coefficients, the stronger the predictor effect. Absent values (-) indicate null influence of the predictor for a given focal group. Values in bold indicate the three most influential variables for each group.

| | *Calliandra* | *Cattleya* | *Chamaecrista* | *Diplusodon* | *Dyckia* | *Habenaria* | Lychnophorinae | Marcetieae | *Mimosa* | *Myrcia* | *Paepalanthus* | Trimezieae | Velloziaceae |
|---|---|---|---|---|---|---|---|---|---|---|---|---|---|
| Climate | 15.84 | 33.66 | 6.79 | 19.99 | 13.48 | 6.49 | 13.74 | 19.96 | 18.28 | 5.92 | 8.50 | 20.37 | 16.41 |
| Soil | 12.44 | 25.11 | 4.33 | 7.04 | 17.91 | 4.17 | 6.97 | 13.65 | 12.26 | 4.08 | 10.10 | 8.88 | 9.10 |
| Geographic | 12.93 | 31.93 | 2.02 | 12.22 | 15.62 | 2.85 | 13.43 | 17.06 | 11.87 | 1.95 | 10.55 | 9.12 | 9.14 |
| Climate & Soil | 19.01 | 35.14 | 7.39 | 20.59 | 24.56 | 8.09 | 15.31 | 21.98 | 18.66 | 6.54 | 12.49 | 21.48 | 16.59 |
| Geo & Climate | 17.39 | 37.95 | 6.96 | 22.04 | 20.09 | 6.70 | 17.39 | 24.05 | 19.41 | 6.03 | 12.06 | 20.66 | 17.63 |
| Geo & Soil | 17.09 | 36.32 | 4.91 | 14.33 | 24.74 | 5.28 | 14.22 | 21.11 | 16.04 | 4.48 | 14.00 | 12.28 | 13.75 |
| All variables (Geo & Climate & Soil) | 20.11 | 39.63 | 7.46 | 22.38 | 28.28 | 8.15 | 17.97 | 24.86 | 19.53 | 6.58 | 14.87 | 21.61 | 17.85 |
| Unexplained | 79.89 | 60.37 | 92.54 | 77.62 | 71.72 | 91.85 | 82.03 | 75.14 | 80.47 | 93.42 | 85.13 | 78.39 | 82.15 |
| Climate alone | 3.03 | 3.31 | 2.55 | 8.06 | 3.54 | 2.87 | 3.75 | 3.75 | 3.49 | 2.10 | 0.87 | 9.32 | 4.10 |
| Soil alone | 2.72 | 1.68 | 0.50 | 0.35 | 8.18 | 1.44 | 0.58 | 0.81 | 0.12 | 0.55 | 2.81 | 0.95 | 0.22 |
| Geo alone | 1.10 | 4.49 | 0.08 | 1.79 | 3.71 | 0.06 | 2.66 | 2.88 | 0.88 | 0.04 | 2.38 | 0.13 | 1.26 |
| Climate + Soil (excl. Geo) | 1.44 | 2.71 | 2.39 | 1.76 | 0.94 | 0.99 | 0.21 | 3.24 | 4.05 | 1.98 | 0.64 | 2.22 | 4.39 |
| Geo + Climate (excl. Soil) | 3.54 | 6.72 | 0.51 | 5.49 | 3.12 | 1.06 | 4.58 | 4.57 | 2.90 | 0.37 | 1.52 | 3.28 | 3.39 |
| Geo + Soil (excl. Climate) | 0.46 | 0.00 | 0.10 | 0.25 | 2.90 | 0.16 | 0.99 | 1.21 | 0.25 | 0.07 | 1.18 | 0.16 | 0.00 |
| Soil + Climate + Geo | 7.83 | 20.91 | 1.34 | 4.68 | 5.89 | 1.58 | 5.20 | 8.39 | 7.84 | 1.48 | 5.47 | 5.55 | 4.53 |

**Table S4.2.** Variance partitioning results for each plant group. Rows show the percentage of deviance explained by each combination of predictors (climate, soil, and geographic). "Unexplained" corresponds to the proportion of deviance not accounted for by the predictors. Intersection terms indicate shared explanatory power among predictor groups.

#### Appendix 5: Reference numbers of occurrence datasets

Reference numbers that refer to datasets are not available for iDigBio, but associated metadata is provided at Zenodo (https://doi.org/10.5281/zenodo.17941129).

##### GBIF

- *Calliandra*: GBIF.org (2024) GBIF Occurrence Download 10.15468/dl.78dj83
- *Cattleya*: GBIF.org (2024) GBIF Occurrence Download 10.15468/dl.jvbdk8
- *Chamaecrista*: GBIF.org (2024) GBIF Occurrence Download 10.15468/dl.nyt4an
- *Diplusodon*: GBIF.org (2024) GBIF Occurrence Download 10.15468/dl.wnptfg
- *Dyckia*: GBIF.org (2024) GBIF Occurrence Download 10.15468/dl.fj4mu6
- *Habenaria*: GBIF.org (2024) GBIF Occurrence Download 10.15468/dl.4x2zep
- Lychnophorinae: GBIF.org (2024) GBIF Occurrence Download 10.15468/dl.jee4dy
- Marcetieae: GBIF.org (2024) GBIF Occurrence Download 10.15468/dl.zq846h
- *Mimosa*: GBIF.org (2024) GBIF Occurrence Download 10.15468/dl.zvqhtj
- *Myrcia*: GBIF.org (2024) GBIF Occurrence Download 10.15468/dl.5e7yqj
- *Paepalanthus*: GBIF.org (2024) GBIF Occurrence Download 10.15468/dl.tyrsbp
- *Trimezieae*: GBIF.org (2024) GBIF Occurrence Download 10.15468/dl.rvaw6s
- Velloziaceae: GBIF.org (2024) GBIF Occurrence Download 10.15468/dl.bfbdaz

##### speciesLink

- *Calliandra*: 20240510134251-0030065
- *Cattleya*: 20241115132136-0027002
- *Chamaecrista*: 20240510142533-0011966
- *Diplusodon*: 20240510141019-0031806
- *Dyckia*: 20241210144324-0005516
- *Habenaria*: 20241117145848-0029682
- Lychnophorinae: 20240515145733-0026206
- Marcetieae: 20240515155457-0011748
- *Mimosa*: 20240413115159-0014841
- *Myrcia*: 20241223154822-0027910
- *Paepalanthus*: 20240515174114-0004034
- *Trimezieae*: 20241129175444-0021626

- Velloziaceae: 20240515161822-0014656
